# Ungulate substrate use in fauna passages

**DOI:** 10.1101/2024.02.08.579517

**Authors:** Jan Olof Helldin, Milla Niemi

## Abstract

Fauna passages are increasingly constructed at major roads and railways to mitigate the negative effects of infrastructure and traffic on wildlife. The function of such passages depends on design, including the construction materials, soil and vegetation. Providing “naturalness” in fauna passages may entail significant costs, yet its benefits are unclear. By using camera trap data, we examined ungulate substrate use in seven passages serving both fauna and local roads (overpasses) and fauna and watercourses (underpasses) in boreal Sweden and Finland. While all substrates were used, during snow-free periods, ungulates used smoother surfaces (fine-grained topsoil, grass, artificial fiber mat, and dirt road) more than expected based on their availability. Coarser surface (stony/rocky ground), shrub, and water were used less than expected. The results for road and water were, however, inconsistent between passages; in one overpass road was instead used less than expected, and in one underpass the water section was used particularly during winter but also by moose wading or swimming through in summer and autumn. General patterns of use largely remained when analysed byspecies, although sample sizes were limited. Despite limitations, our study offers valuable insights for planning and constructing fauna passages. To our knowledge, this study was the first of its kind describing how ungulates use different substrates in fauna passages. We suggest to conduct further research on this issue, for example by more detailed study of animal trajectories through passages and by experimental modification of substrates.

## Introduction

Transport agencies worldwide are increasingly constructing large fauna passages at major roads and railways to reduce both wildlife-vehicle collisions and the barrier effects of these transportation routes on wildlife (Iuell et al. 2003; Grilo et al. 2010; Smith et al. 2015a; Hlavac et al. 2019; Denneboom et al. 2021). Such fauna passages may be vegetated bridges, viaducts or culverts of different sizes, construction materials and shapes, but have in common that they are constructed or adapted to meet the requirements of the target species or taxa. Many practical guidelines exist to aid the construction of fauna passages (e.g., Clevenger and Ford 2010; Queensland Government 2010; Smith et al. 2015b; Hlaváč et al. 2019; Chrétien et al. 2022; see also national guidelines accessed through Transport Ecology Guidelines Portal 2024). Most of these advocate for, *inter alia*, large openness (maximizing width and height for underpasses, while minimizing length), using natural materials and substrate, providing shelter and connected habitat structures through the passage, screening from traffic disturbance, as well as minimizing other human disturbances.

While the dimensions of the constructions is generally the most cost-driving factor for fauna passages (Sijtsma et al. 2020; Helldin 2022), adaptations to create “naturalness” by vegetation and soil conditions in the passage may also entail significant costs. The degree to which fauna passages can be combined with non-wildlife structures such as minor roads, hiking trails or watercourses may also be a cost-driving factor as it impacts decisions of total number of bridges and fauna passages needed.

Regarding the ground surface or substrate in large fauna passages, the current Swedish construction guideline (Trafikverket 2022) recommends natural vegetation on a bed of natural soil. Minor unpaved roads with limited traffic can be accepted, and a dry bank ≥20 cm above high water level should be kept along watercourses. Planners, however, still express uncertainties about combining passages for larger fauna with local roads or streams, due to the risk of human disturbance and the potential loss of effective passage width to road or water. For ungulates, which are the main target taxon for larger fauna passages in northern Europe, results of avoidance or preference of certain substrates in fauna passages are sparse and inconclusive (Denneboom et al. 2021). To assess the function for ungulates of non-wildlife over- and underpasses in Sweden, Seiler et al. (2015) proposed to exclude road and water surfaces when calculating the effective passage width, thereby implying that these surfaces would not be used by ungulates.

The aim of this study was to describe how ungulates use substrates in fauna passages, as revealed by the animal’s trajectories through the passage. Studies were conducted year-round at three underpasses (viaducts) and four overpasses (green bridges) in boreal Sweden and Finland. While these passages may be utilized by many different species, we perceive that the passages were primarily adapted to meet the requirements by ungulates. Particular focus was put on the potential avoidance of streams (in the case of underpasses) and low-traffic roads (in overpasses) going through the passage. We discuss how the results may guide future construction of passages built or adapted for boreal ungulates.

## Method

### Fauna passages included in the study

We selected seven fauna passages to include in the study (Table 1), out of a total of 35 under- and overpasses that were monitored for ungulate use within larger research projects in Sweden (Håkansson 2020; Raud Westberg and Ellvin 2021) and Finland (Niemi 2021) during 2018–2022. The selection was arbitrarily based on the following criteria:

**Table 1.** Fauna passages monitored for use of substrate by ungulates, with basic design and monitoring period.

|  | Passage type | Infra-structure | Area | Monitoring start | Monitoring end |
| --- | --- | --- | --- | --- | --- |
| **Kvarnbäcken** | Underpass/viaduct | Railway | Sweden | July 2019 | July 2020 |
| **Aavajoki** | Underpass/viaduct | Railway | Sweden | Nov 2019 | Nov 2020 |
| **Keräsjoki** | Underpass/viaduct | Railway | Sweden | July 2019 | July 2020 |
| **Sangijärvi** | Overpass/green bridge | Railway | Sweden | Nov 2019 | Oct 2020 |
| **Sammatti** | Overpass/green bridge | Motorway | Finland | Dec 2019 | Nov 2020 |
| **Loviisa<sup>a</sup>** | Overpass/green bridge | Motorway | Finland | Dec 2019 | Nov 2020 |
| **Kärmekorpi** | Overpass/green bridge | Motorway | Finland | Dec 2019 | Nov 2020 |

<sup>a</sup> Labelled “Loviisa 2” in Niemi (2021).

- Minimum width 20 m and containing a variety of substrates (≥2), to allow animals to actively select substrate.
- Monitored at least ca 1 year.
- Camera trap set up allowed equal (un-biassed) coverage of the entire passage width.

In one case, however, a bridge was included (Loviisa) despite a 5m strip behind a railing not being reliably covered by the camera. We judged the strip being of less interest and limited the analyses to the rest of the passage width.

By this selection, four passages (three viaducts and one overpass) along the Haparandabanan railway in northern Sweden and three passages (all overpasses) along motorway E18 in southern Finland were included in the analyses (Fig. 1). All seven passages are defined by the respective infrastructure administration as multifunctional fauna passages, i.e., the intended function is for both fauna and any other purpose; in these cases watercourses or local, low-traffic roads, and including recreational use. Structural details and frequency of local traffic and other human use of each passage are presented in Supplementary Information no. 3.

**Fig. 1.**
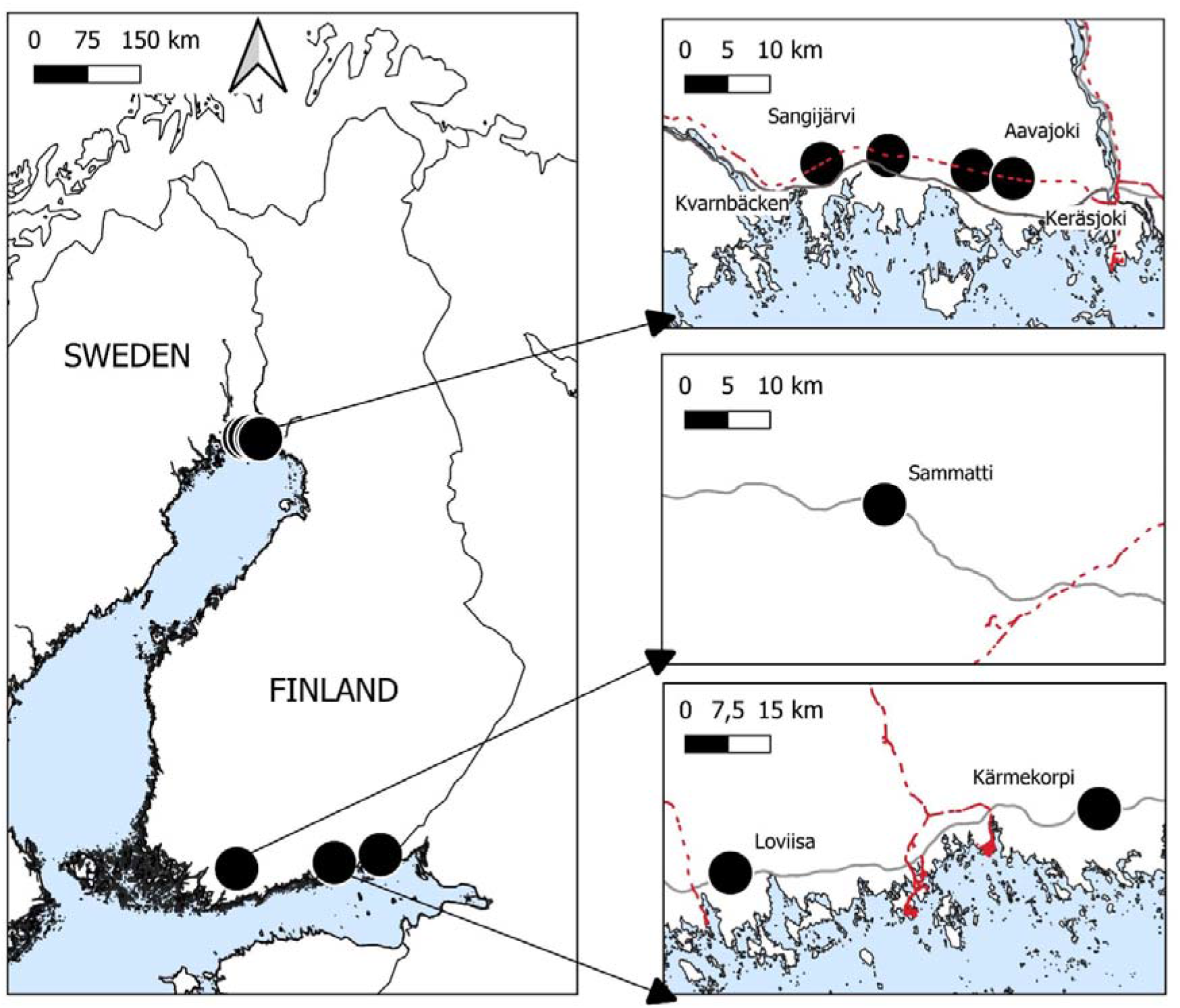
Fauna passages included in the study (black circles) and major roads (grey lines) and railways (red hatched lines). The Swedish study passages (inset map at the top) were all located on railway, while the Finnish study passages (second and third insets) were on motorway. Background information from Trafikverket (2024a, b), Finnish Transport Infrastructure Agency (2024), National Land Survey of Finland (2023), Eurostat (2020) and Flanders Marine Institute (2021)

### Study areas

The study area in northern Sweden is situated at 65°N 23°E, in northern boreal zone dominated by a mix of forest and bogs, and with stable winter conditions from November through April, and average snow cover being 60-70 cm (standardized date March 15, Swedish National Atlas 1995). Ungulate species that occur in the area are moose (*Alces alces*), roe deer (*Capreolus capreolus*) and reindeer (*Rangifer tarandus tarandus*); the latter being semi-domestic but free-ranging most of the year. The study area in southern Finland is situated at 60°N ∼25°E, in southern boreal zone dominated by a mix of forest and farmland, with winter conditions being more erratic, with snow cover typically from January to March, and reaching 10-20 cm (standardized date March 15, Finnish Meteorological Institute 2024). Ungulate species occurring in this area are moose, roe deer, white-tailed deer (*Odocoileus virginianus*; non-native) and wild boar (*Sus scrofa*). The terrain in both study areas is generally smooth and lowland (<100 m above sea level).

### Camera trapping

We monitored ungulates in the passages using motion-triggered cameras (Browning 2017 Spec Ops Advantage Trail in the Swedish passages, Uovision UM785-3G in the Finnish) with infrared (IR) motion sensor, IR flash and IR night vision, and with the detection range of motion sensor and flash specified by manufacturer as minimum 16 m (Uovision) or 24 m (Browning). We set the cameras to rapid fire 3-5 images when being triggered, thereafter with a delay of 5 sec (Browning) or 60 sec (Uovision) to the next triggering.

We mounted the cameras in the middle of the passages (some of the passages also had cameras distributed outside the passage, but these cameras where not used in the present study), at approximately 1 m above ground, attached to pre-existing fence poles or in few cases new poles. We placed the cameras at one or both sides of the passage and facing the passage center (Fig. 2), and in one case we also mounted cameras near a viaduct’s center pillar and facing the sides, accordingly aiming at covering the entire width of the passage. We did some careful vegetation cutting in front of cameras to minimize false “wind triggers”. We revisited cameras once per 1–3 months in order to check functionality and download images. For the aim of this study, ungulate use was recorded for one year (Table 1 and Supplementary Information no. 3).

**Fig. 2.**
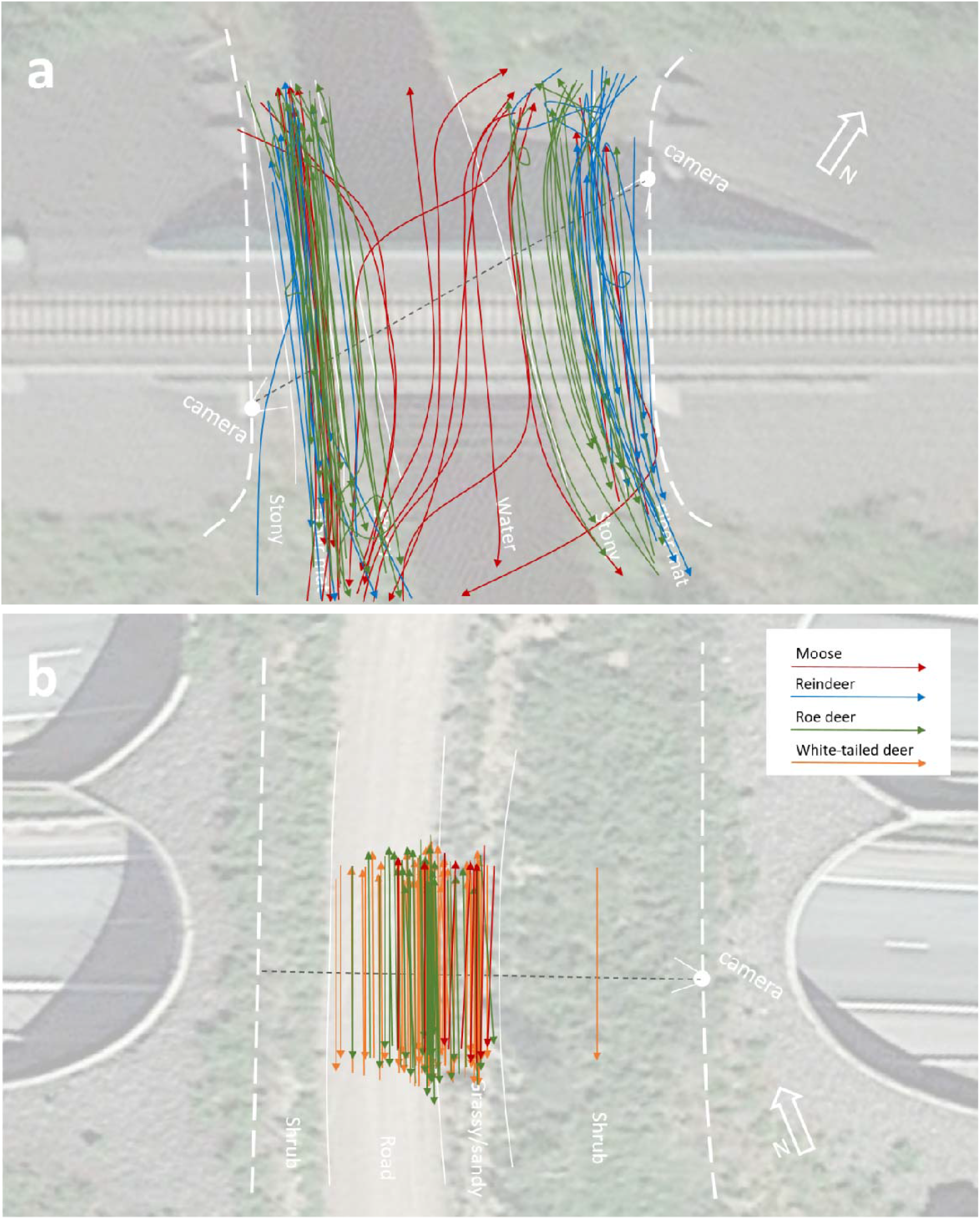
**a-b** Examples of ungulate trajectories over fauna passages. a) Kvarnbäcken underpass, b) Sammatti overpass, both at snow free conditions. Red = moose, blue = reindeer, green = roe deer, orange = white-tailed deer. Dotted line = record line. Background images show camera placement and direction, perimeter fencing, and available substrates

### Image handling

We combined all consecutive images of ungulates (singletons or groups of conspecifics) at a passage separated by ≤10 min into one crossing event. This time limit was arbitrarily set to avoid the most immediate temporal dependence between observations (Burton et al. 2015). We used only events that we judged showing animals successfully crossing the passage, i.e., coming in from one side and leaving towards the other. For each event, we noted the ungulate species, the summed number of individuals, date and time, snow cover (Y/N) and direction of movement (south/north). We excluded a small number of crossing events where species could not be definitively identified (roe/white-tailed deer; Finnish data). For each event, we recorded the substrate where the animal crossed a predefined imaginary line over the passage in front of the cameras (record line; Fig. 2a–b). In the case of groups, we recorded only the first crossing individual. We classified substrates into six types:

- Grassy/sandy (spontaneous herbaceous or graminoid vegetation on sandy soil)
- Stony/rocky (sometimes with sparse herbaceous vegetation)
- Fiber mat (artificial coco liner ca 5 cm thick, rolled out on stony/rocky substrate)
- Shrub (woody vegetation ca 0.5–2 m height, mainly with ground covered by low vegetation)
- Water (river or stream)
- Road (local gravel/dirt road)

Substrates grassy/sandy and road were only represented in the overpasses while stony/rocky, fiber mat and water were only represented in the underpasses (Table 2).

**Table 2.**
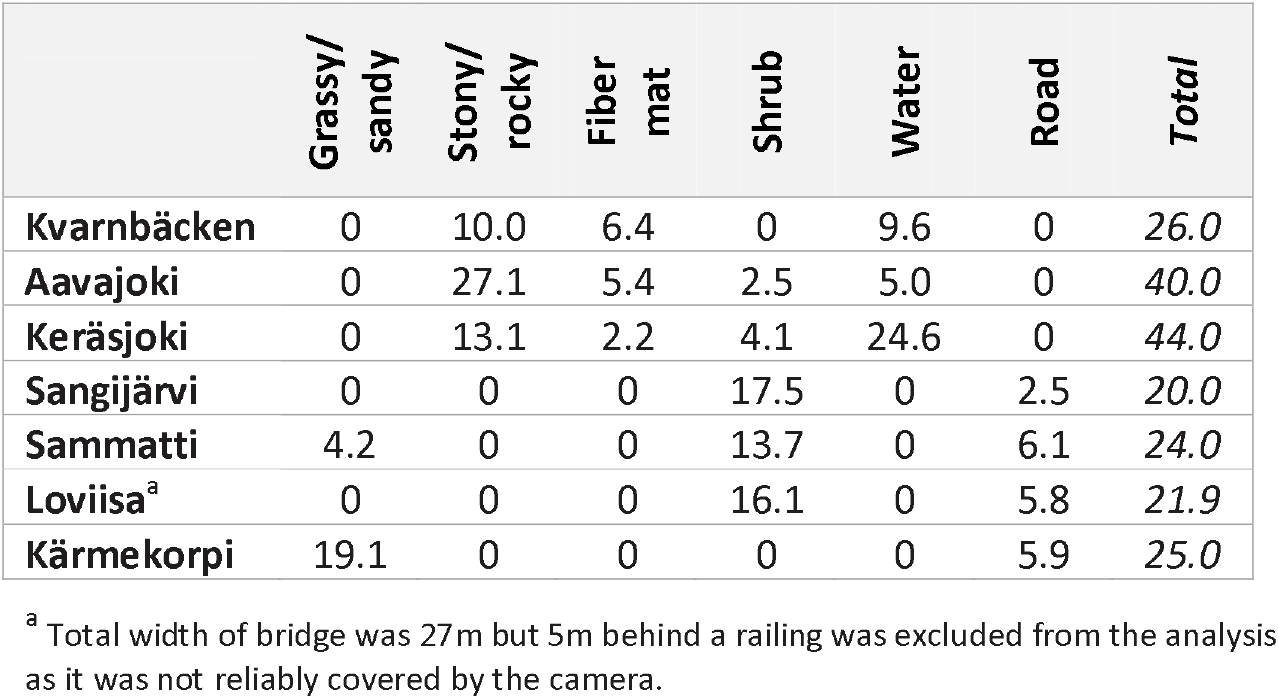
Substrates available in the fauna passages, in width (m) along the record line.

As a complement, we mapped each trajectory on a spatial map (Fig. 2a–b).

### Data analyses

We compared the substrates used by ungulates (when crossing the imaginary line) with available substrates using a chi-square (χ^2^) goodness-of-fit analysis conducted in Excel, separate for each fauna passage and season (snow-free/snow covered ground), and separate for each ungulate species plus all species taken together. We excluded analyses with any cell with an expected value lower than 5 (as recommended by Siegel and Castellan 1988). For all significant analyses (p < 0.05), we recorded the relative contribution (proportion of total χ^2^) of each substrate type, and pointed out preferences (observed > expected) and avoidances (observed < expected).

For the particular focus on watercourses and roads through the passage, we conducted additional chi-square goodness-of-fit analyses of these two substrates compared to other substrates pooled together, separate for each fauna passage and season (snow-free/snow-covered ground), for each ungulate species plus all species taken together, and using the same analysis exclusion criterion (i.e., any cell with an expected value lower than 5).

We separated seasons by records of snow-covered ground on images, which in the Swedish passages corresponded to mid-October to early May (ca 200 days) and in the Finnish a number of shorter periods occurring from end of November through early April (a total of <15 days).

## Results

A total of 983 ungulate crossing events were recorded in the study (Table 3), with moose and roe deer recorded in all passages, reindeer only in the Swedish, white-tailed deer and wild boar only in the Finnish; wild boar only at very few occasions. A majority of crossing events (719) were from the snow-free season. During the period of snow-covered ground, most (90%) crossing events involved moose or reindeer. Only the Swedish passages allowed analyses from snowy period.

**Table 3.**
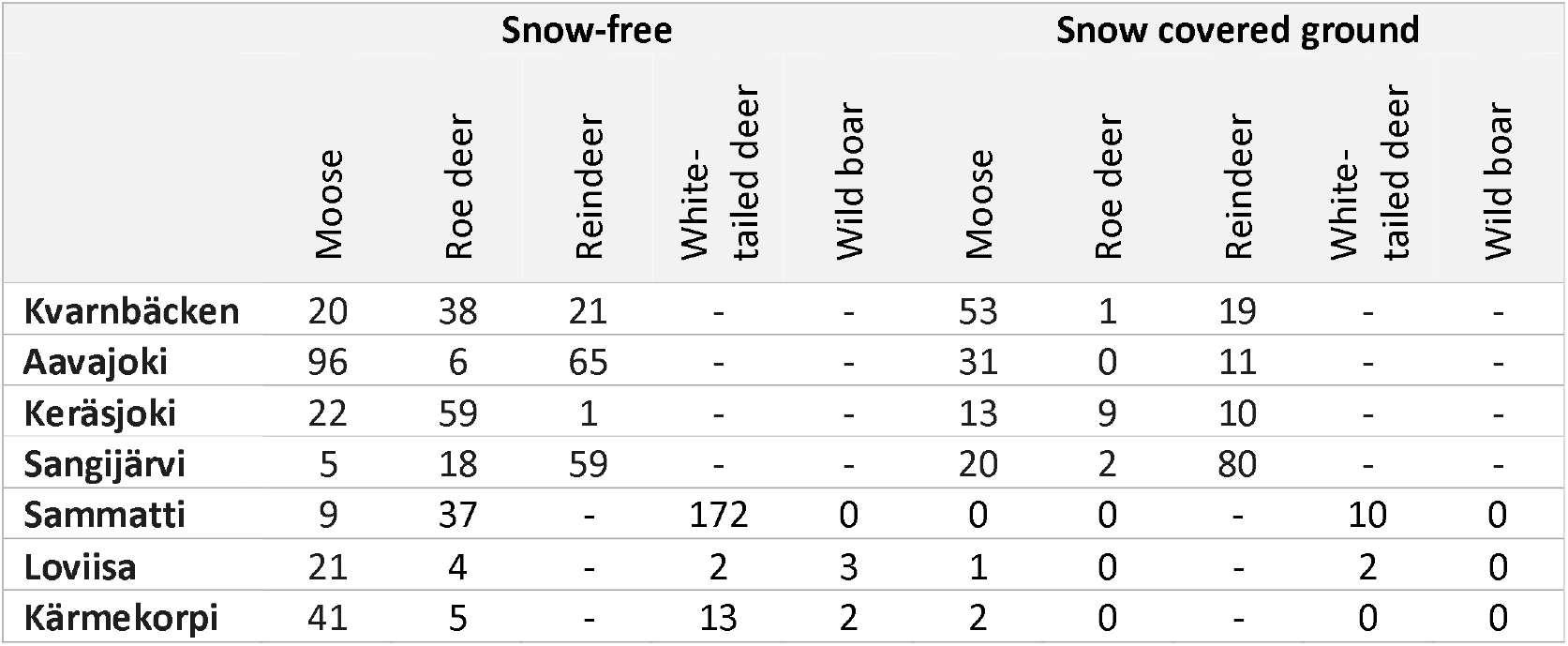
Number of ungulate crossing events included in the study, separated by season (snow-free/snow covered ground) and species.

Although all types of substrates were used, the extent varied, and ungulates displayed a non-random use of substrate in all analyses (Table 4), except for the passage Loviisa.

**Table 4.**
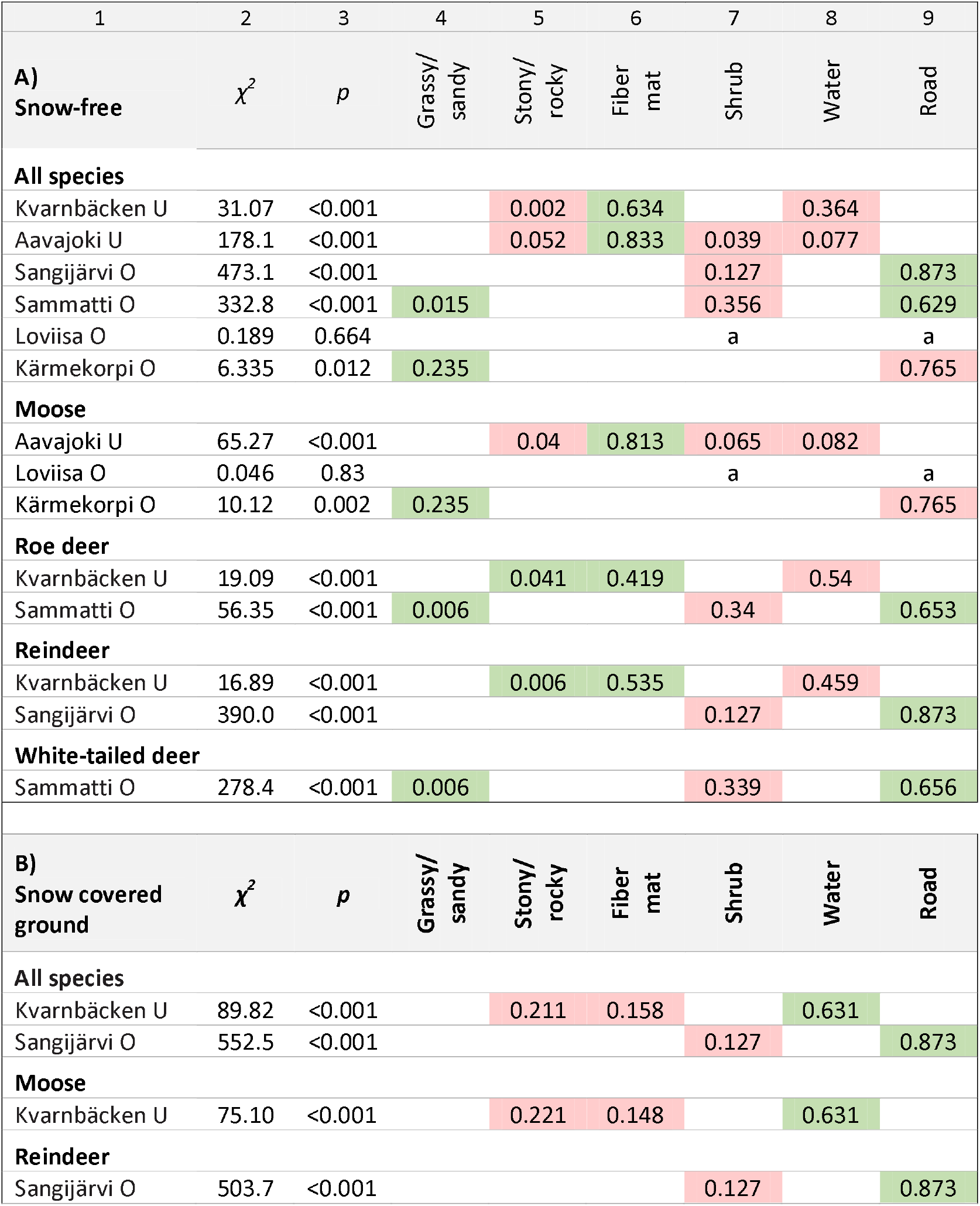
**A-B** Results from analyses of ungulate substrate use in relation to availability, during snow-free season and snow covered ground, respectively. U = underpass, O = overpass. Values in columns 4-9 are the relative contribution to the χ^2^ in column 2; green cells = preferred substrate (observed > expected), red cells = avoided substrate (observed < expected), a = substrate used according to availability, empty cells = substrate not available within that passage. Only analyses reaching required sample size (all expected values ≥5) are presented.

In the *snow-free season* (Table 4A), the general pattern was that fiber mat and grassy/sandy ground (only available in underpasses) and road (only in overpasses) were used more than expected by availability; this was most pronounced for fiber mat (large contribution to χ^*2*^). Conversely, water and stony/rocky ground (in underpasses) and shrub (available in both under- and overpasses) was used less than expected by availability; this was however less well pronounced for stony/rocky ground (≤5% contribution to χ^*2*^). For road, the results were inconsistent, with much more use than available in two passages but much less in one. When analyzed at the species level, these general patterns mainly persisted. However, roe deer and reindeer utilized stony/rocky ground more than available in the Kvarnbäcken underpassage (still with low contribution to χ^*2*^).

In the period of *snow-covered ground* (and, notably, also ice covered water), water and road were used more than expected by availability, while stony/rocky ground, fiber mat and shrub were used less (Table 4B). However, only two passages allowed analysis for snowy season – the underpass at Kvarnbäcken and the overpass at Sangijärvi – and the result was to a large extent attributed to moose in the former case and to reindeer in the latter.

The particular analyses of water and road, respectively, allowed analysing a few more combinations of passages and species (Table 5A–B), and largely underlined results above. *Water* was used less than expected by availability during snow-free (and ice-free) season, in all cases except for moose in one case (Kvarnbäcken) where water was used according to availability. In the period with snow and ice, the use of water differed between passages; more than expected by availability in one case (Kvarnbäcken) and less in the other two cases (Aavajoki and Keräsjoki). For *road*, the results remained inconsistent, with road used more than expected by availability in two passages (Sangijärvi and Sammatti, for the former also with snow-covered ground) but less in one (Kärmekorpi) and according to availability in one (Loviisa).

**Table 5.**
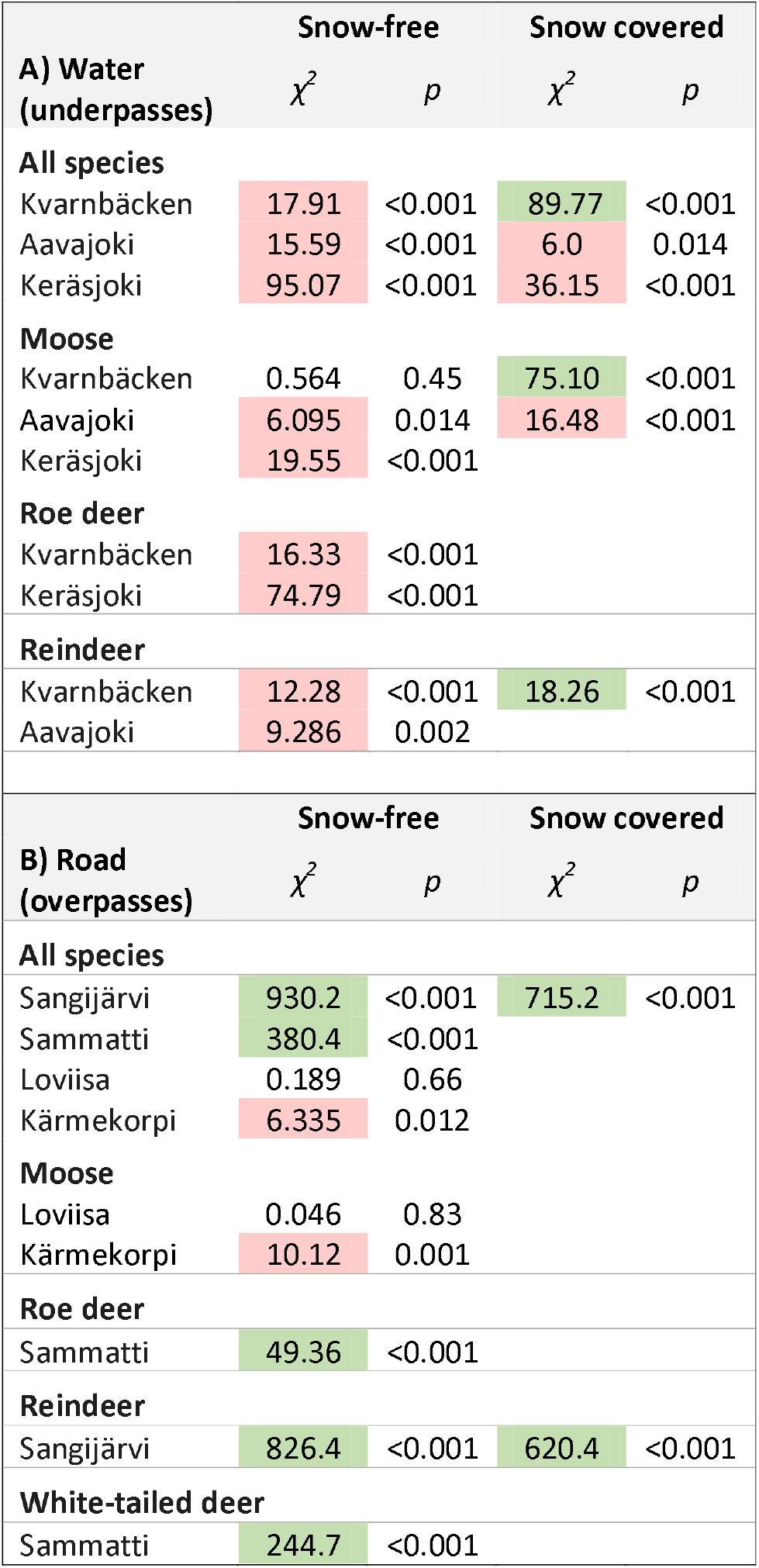
**A-B** Results from analyses of ungulate use of water and road in relation to other substrates, during snow-free season and snow covered ground, respectively. Green cells = preferred substrate (observed > expected), red cells = avoided substrate (observed < expected). Only analyses reaching required sample size (all expected values ≥5) are presented.

## Discussion

Our study indicated that fine-grained topsoil, grass, and fiber mat were preferred by ungulates in fauna passages during snow-free conditions (Fig. 3a–b). Although we can only speculate about the reason for this preference, we acknowledge that these surfaces are smoother, and may be perceived by animals as more comfortable for walking or running. It has been argued that fine-grained soil and smooth surfaces should be used in fauna passages, rather than coarse materials such as macadam or riprap (Smith et al. 2015b; Trafikverket 2022). In line with that, we found that ungulates used coarser surfaces (Fig. 3c) slightly less than expected based on its availability. This was true also in snowy conditions.

**Fig. 3.**
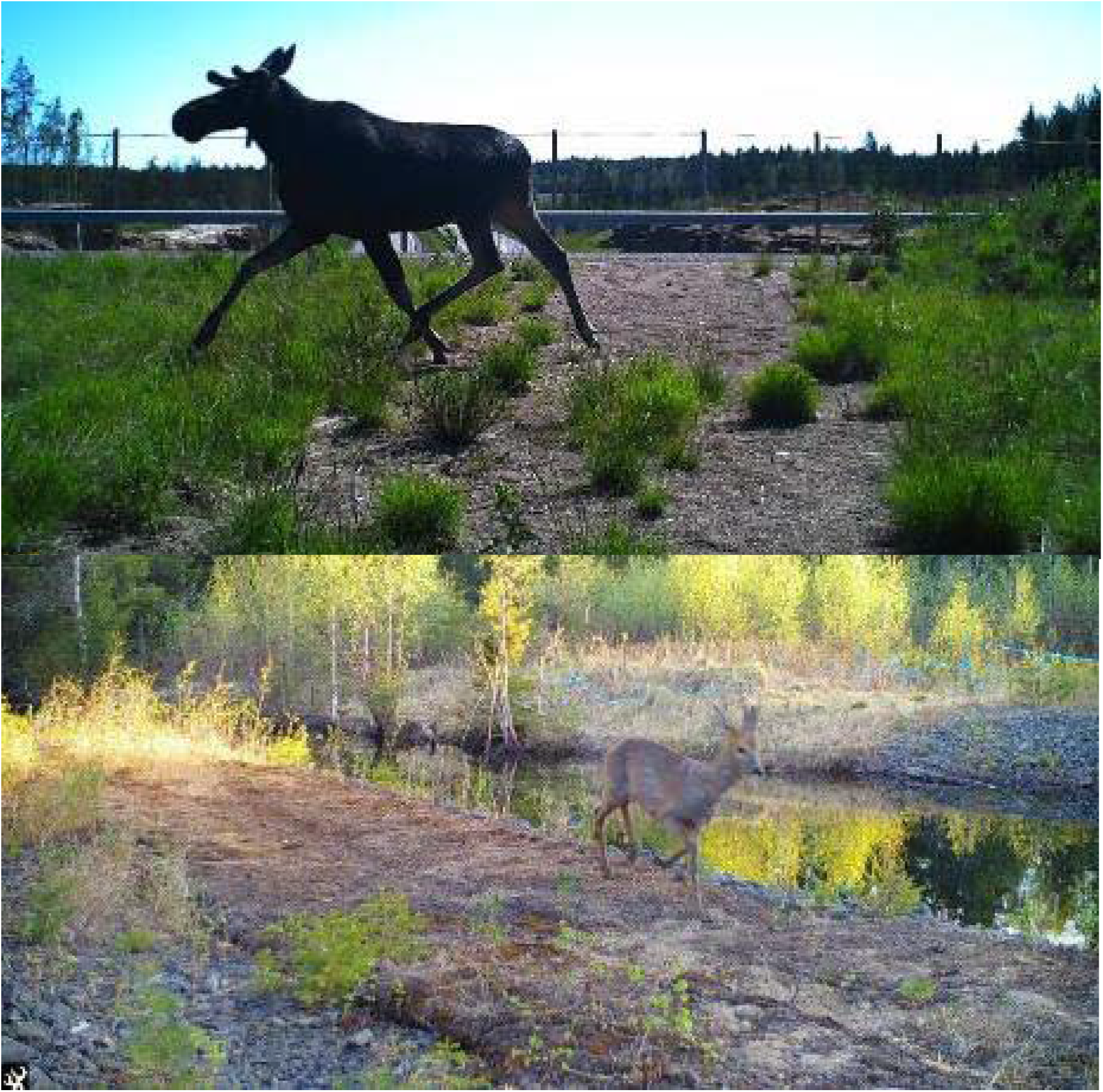
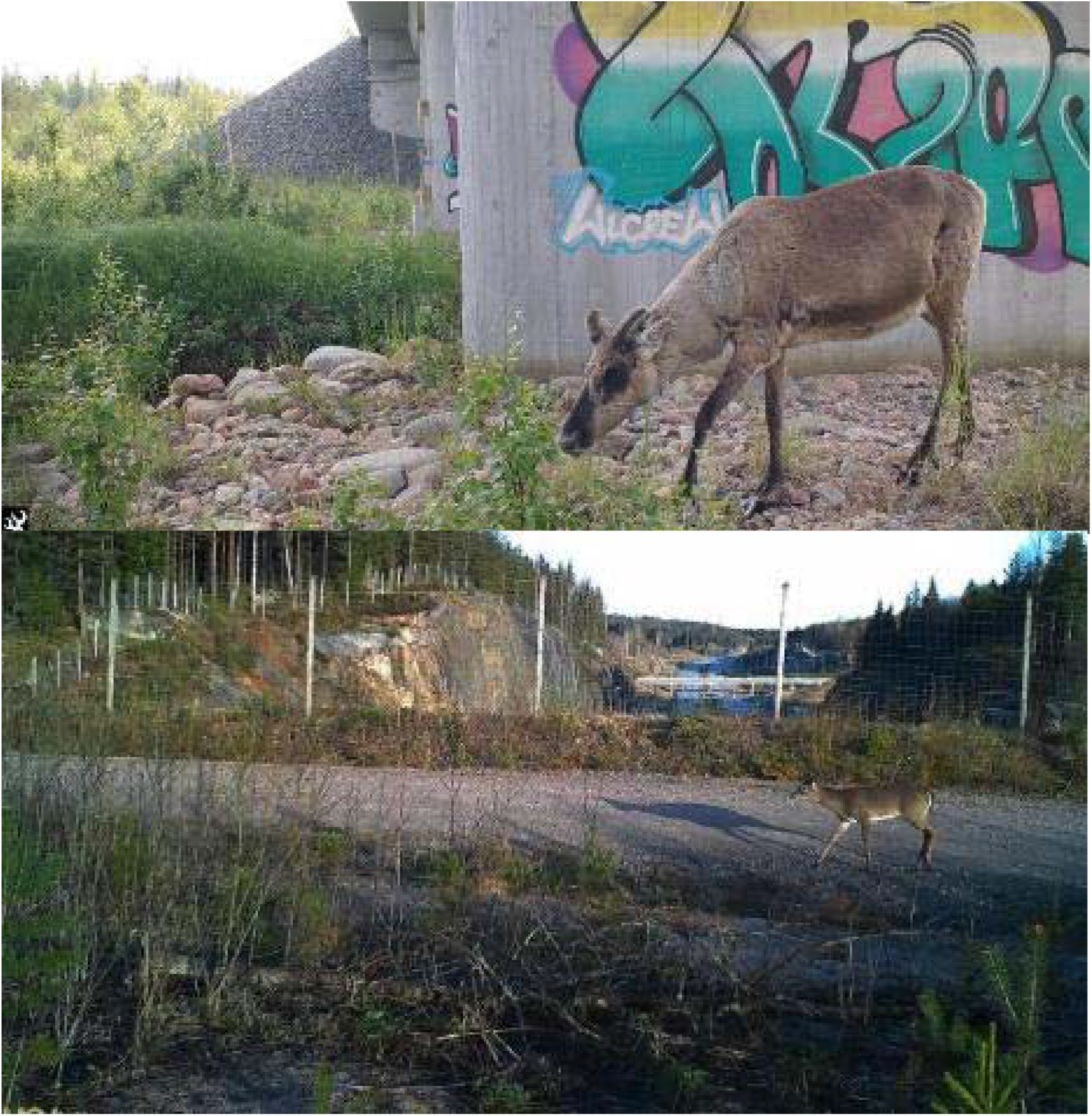
**a-d** Examples of records of ungulates crossing through fauna passages on different substrates. a) moose on grassy/sandy substrate (Kärmekorpi, 12 Jun, 2020); b) roe deer on fiber mat (Kvarnbäcken, 2 Jun 2020), c) reindeer on stony/rocky substrate (Aavajoki, 25 Jun 2020), d) white-tailed deer on road (Sammatti, 6 May 2020)

Smooth and easy-to-walk-on surface may apply also to road (local gravel or dirt road), which was clearly preferred in two of the overpasses in our study (of which one included data also from snowy conditions; Fig. 3d), and used according to availability in one. The analyses on species level showed that this preference of roads was attributed to roe deer, reindeer, and white-tailed deer. In the overpass where road was instead avoided, the alternative substrate was grassy/sandy, i.e., another smooth surface, but also the site where moose was the most frequent species. It has been proposed that ungulates (reindeer) may use minor, low-traffic roads as transportation corridors (e.g. Strand et al. 2018). Although some studies indicate that ungulates in forest landscapes avoid minor roads (Laurian et al. 2008; Mathisen et al. 2018), this is likely due more to higher risk of hunting mortality and predation by large carnivores than to the character of the road surface.

Furthermore, the results indicated that shrub as a substrate (or habitat) in fauna passages was avoided by ungulates, although shrub should appear natural and provide shelter and even food (depending, of course, on species), and although shrub in the current cases forms more or less continuous habitat corridors through these passages and thereby connects habitats across the infrastructure. In contrast to shrub, all other solid-ground substrates (grassy/sandy, stony/rocky, fiber mat, road) are more or less open habitats, and may be preferred for that reason.

As a general conclusion, we propose that ungulates may require view rather than shelter when crossing through fauna passages, and accordingly avoid denser habitats such as shrub. Moving through a fauna passage is likely to be an aggravated situation for wildlife, with an elevated risk of predation or strife (Mata et al. 2015), and open areas near passages may facilitate predator avoidance and escape (Clevenger and Waltho 2005; Denneboom et al. 2021). However, we cannot exclude the possibility that the shrub in our cases may have provided shelter from the side (for example from traffic disturbance) or functioned as a guiding habitat structure, and therefore may have served a purpose for ungulates even when not used directly.

Water was generally avoided by ungulates in fauna passages, but with the prominent exception of Kvarnbäcken underpass where moose used the water section year-round and reindeer in winter (Fig. 4a–c). In this passage, much of its span is dedicated to a gently flowing, deep but narrow river (ca 10 m wide). In winter, the river froze and the ice created a solid floor used by the vast majority of moose and reindeer that crossed in the six winter months (November-April). However, also in the rest of the year, moose used the river for crossing, either by wading along the river bank or swimming through the deeper part. We recorded the same behaviors in the other two underpasses, however less frequently. Interestingly, as far as we could observe (by additional records from cameras outside of the passages), in all these occasions moose approached on land, went into the river some 10 m before the passage, swam or waded through, and got up again on land shortly after crossing.

**Fig. 4.**
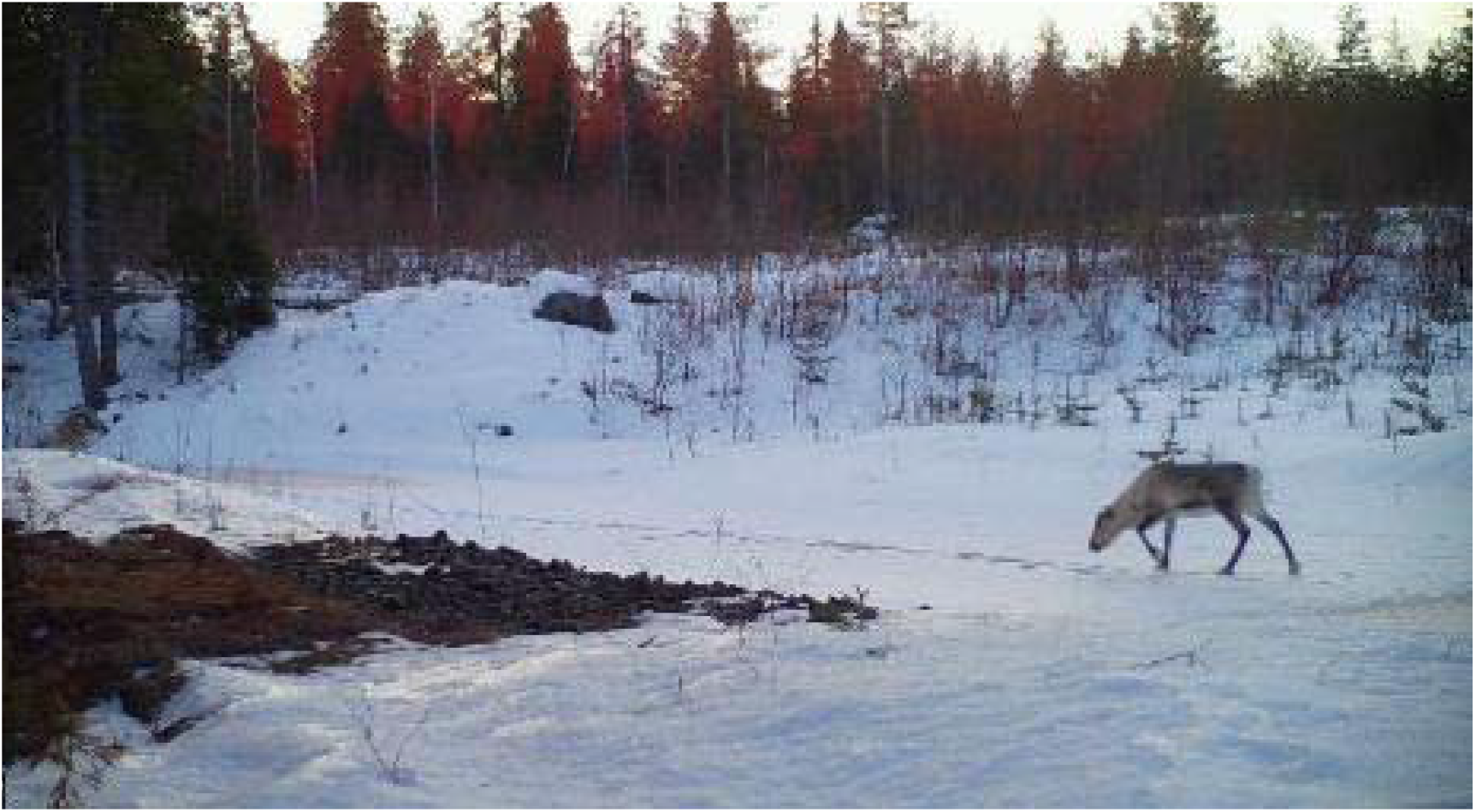
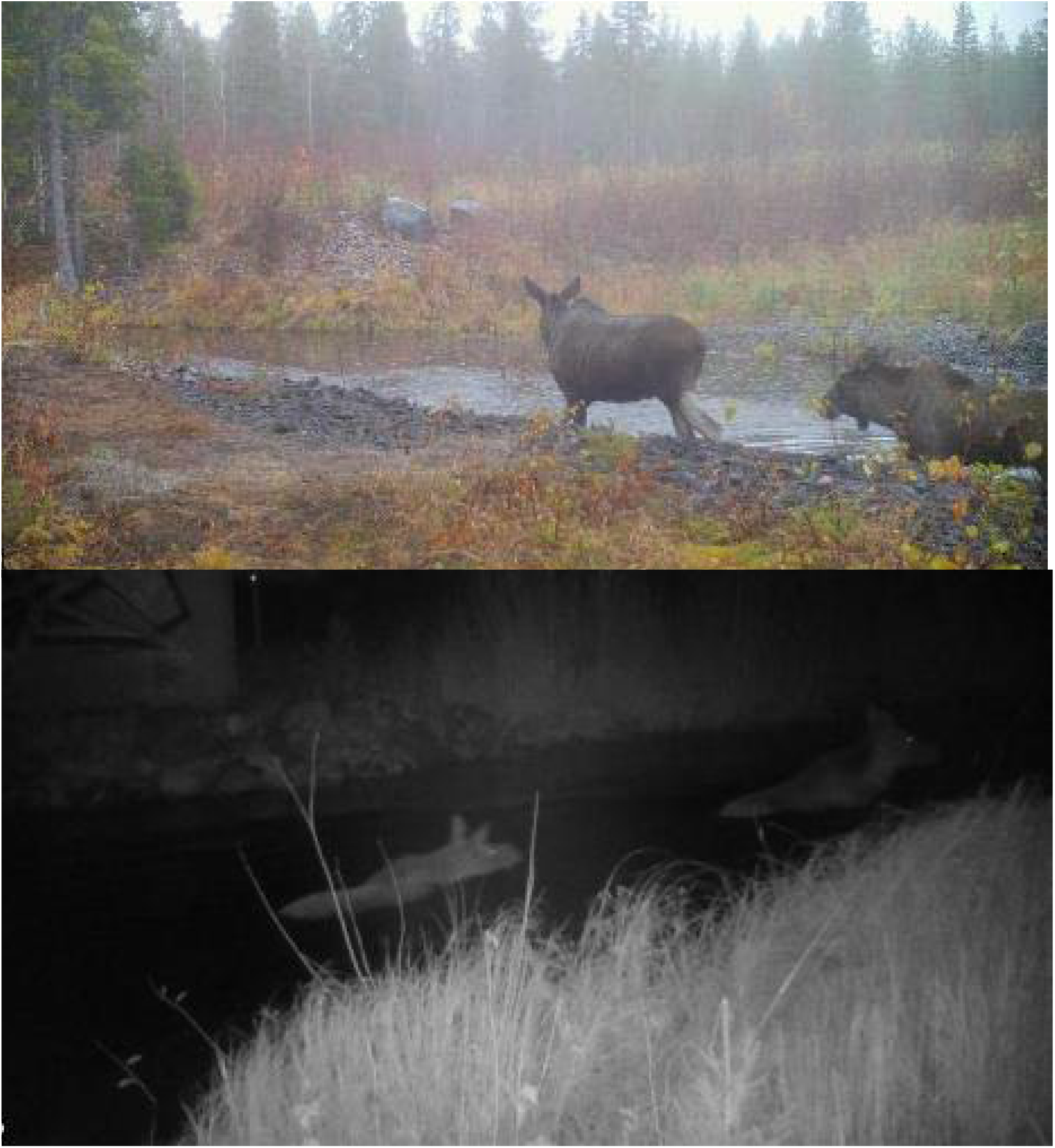
**a-c** Ungulates crossing fauna passages via a watercourse. a) reindeer walking on river ice (Kvarnbäcken, 31 Dec 2019); b) moose wading along river bank (Kvarnbäcken, 24 Oct 2019); c) moose swimming along river (Aavajoki, Oct 13 2020)

Our study had a number of limitations, related to the study setup and confounding factors, so conclusions must be drawn with care. We could only include a limited number of fauna passages in our study. These passages varied in type, size and substrate composition, and were located in two areas with different species abundance. Therefore, the observed patterns might be influenced by these local circumstances and may not necessarily be universal. Also, other structural factors than substrate could potentially have impacted the trajectory along which ungulates crossed through the passages, for example if they approached and entered the passage along preferred guiding structures in the surroundings (for example perimeter fencing or established wildlife trails), or if they avoided moving in the direct proximity to technical elements in the passage such as the concrete bridge abutments.

We also acknowledge the risk of pseudo-replication (non-independent data) within passages, deriving from a smaller number of individuals returning to a passage, and therefore that results may reflect individual choices or habits. We did not systematically record individuals but find it most likely that local residents use fauna passages repeatedly, and we could incidentally recognize a few returning individual roe deer bucks by their antler shape and reindeer by their fur patterns. Another aspect to pseudo-replication is that ungulates may follow each other’s trails based on scent and tracks, probably even after much longer period than the 10 min we set to define a separate crossing event.

Despite these limitations, our work provides insight in a topic on which surprisingly little research has been conducted. Given the significant implications for planning and costs associated with fauna passages, there’s a clear need for more applied research in this area. The importance of substrate and microhabitats in fauna passages have been studied for smaller vertebrate species such as small mammals, reptiles and amphibians (Georgii et al. 2011; Connolly-Newman 2013; Andrews et al. 2015), but much less so for large mammals (Denneboom et al. 2021). While authors have stressed the importance for ungulates of vegetation, natural soil, level ground surface, and dry pathways along watercourses through fauna passages, this has usually been addressed by analyzing crossing rates vs. crossing failures (Smith 2003; Eco-Kare International 2020; Denneboom et al. 2021), hence comparisons on a between-passage level. Our study may serve as a complement to these previous studies, by illustrating also how ungulates use, and possibly select, substrates within passages.

## Conclusions and recommendations

Our results largely support the current, broad recommendation to provide natural soil and vegetation in fauna passages (Väre et al. 2003; Chrétien et al. 2022; Rosell et al. 2022; Trafikverket 2022).

However, we point out some opportunities for clarifications on this matter. The results indicate that open habitats are more important for ungulates than previously acknowledged, and accordingly, encroaching woody vegetation may need to be thinned or cleared. We are aware that dense vegetation in fauna passages may benefit smaller species and thus make the passages functional for a broader array of species. This should, however, not be at the cost of the effectiveness for the primary target species. More emphasis may be put in the guidelines on the value of a smooth ground surface, such as that created by fine-grained topsoil and fiber mats, and on the potential value of minor roads with low traffic, as they can function as conduits of ungulate movement. We also recommend to consider the function of watercourses for some ungulate species, not only as a large scale guiding structure but also as a unique substrate providing the opportunity to wade or swim through the passage, and to walk through in winter in regions with stable winter cold.

All these conclusions are tentative, and we hope that our study will open up for further research of vegetation, soil and structures in and around fauna passages. We suggest to plan study setups that allow recording of entire trajectories of animals through the fauna passage, taking into consideration the substrates, guiding structures and microhabitats both in the passages and in their direct vicinity; i.e., in the area planned and managed by the road or railway agency. We find this field particularly suitable for experimental analyses, by modifying substrates within a passage and monitoring any changes in the use by animals. For example, coarse substrate could be covered with fine-grained soil or fiber mat, and shrub could be planted or removed, in a Before-After setup. Further study of watercourses and minor roads should aim at sorting out whether these facilitate or obstruct animal movements, since both our results and previous research (as reviewed in Denneboom et al. 2021) are inconclusive on these points.

## Supporting information

Data suppl 1 Crossing events

Data suppl 2 Individual trajectories

Data suppl 3 Fauna passages info

## Supplementary Information

The following supplementary information is supplied:

- Suppl Info 1. Basic data of all crossing events used in the analyses.
- Suppl Info 2. Spatial maps of individual trajectories for all crossing events used in the analyses.
- Suppl Info 3. General information on fauna passages included in the study.

## Acknowledgements

We are grateful to the many colleagues and assistants involved in data collection and analysis (Ida Anomaa, Christine Godeau, Charlotte Hansson, Emma Håkansson, Jan-Erik Innala, Victor Johansson, Fabian Knufinke, Torbjörn Nilsson, Lars-Gunnar Nyström, Mattias Olsson, Andreas Seiler, Andreas Öhlund), and to Marcus Elfström, Fabian Knufinke, Roy Rea and an anonymous reviewer for valuable comments on earlier drafts of this paper.

## Declarations

## Ethical approval

The collection and processing of the photographic data complied with the EU’s General Data Protection Regulation (GDPR) and following the national regulations for Sweden (SFS no. 2018:1200) and Finland (FFS no. 5.12.2018/1050).

No animals were handled, forced or held in captivity for the research and no ethical approval needed.

## Author contributions

J.O. Helldin conceptualized and designed the study, performed the analyses, and wrote the first draft of the manuscript. Both authors contributed in collecting and preparing data, and in revising the manuscript.

## Availability of data and materials

All data supporting the findings of this study are available within the paper and its Supplementary Information files.

## Statements and declarations

The authors declare no competing interests.

