## Supplementary material for "Ungulate substrate use in fauna passages": Data suppl 2 Individual trajectories

**Data supplement no. 2 to manuscript Ungulate substrate use in fauna passages**

Spatial maps of individual trajectories (events), separated by site, species and season (snow-free or snowy)
Figures in maps refer to column F in Data supplement no. 1.
Camera placement and record lines (dotted) are denoted.

*J.O. Helldin & M. Niemi*
*Submitted to European Journal of Wildlife Research*

### Kvarnbäcken moose, snow free

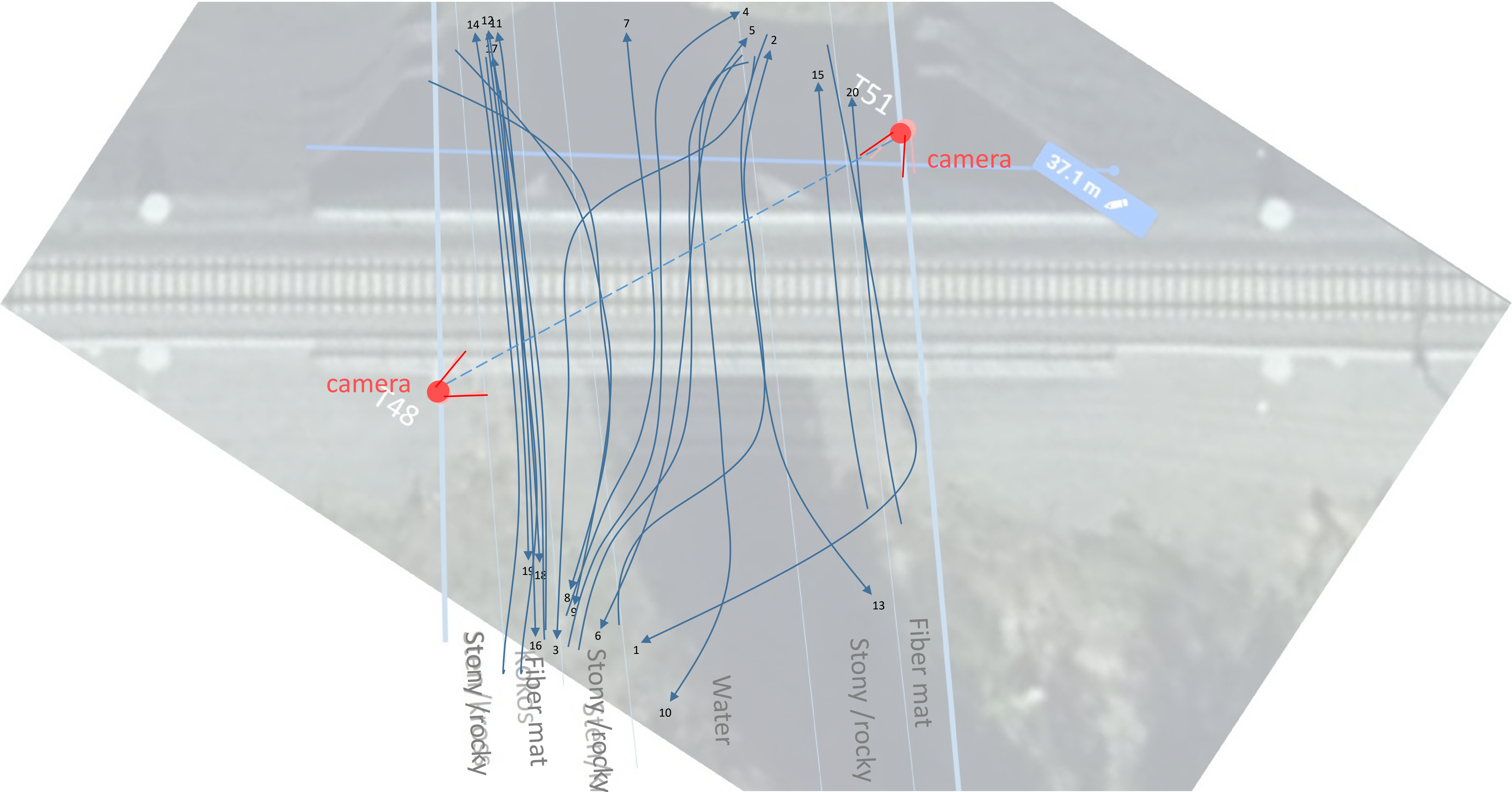

### Kvarnbäcken moose, snowy ground

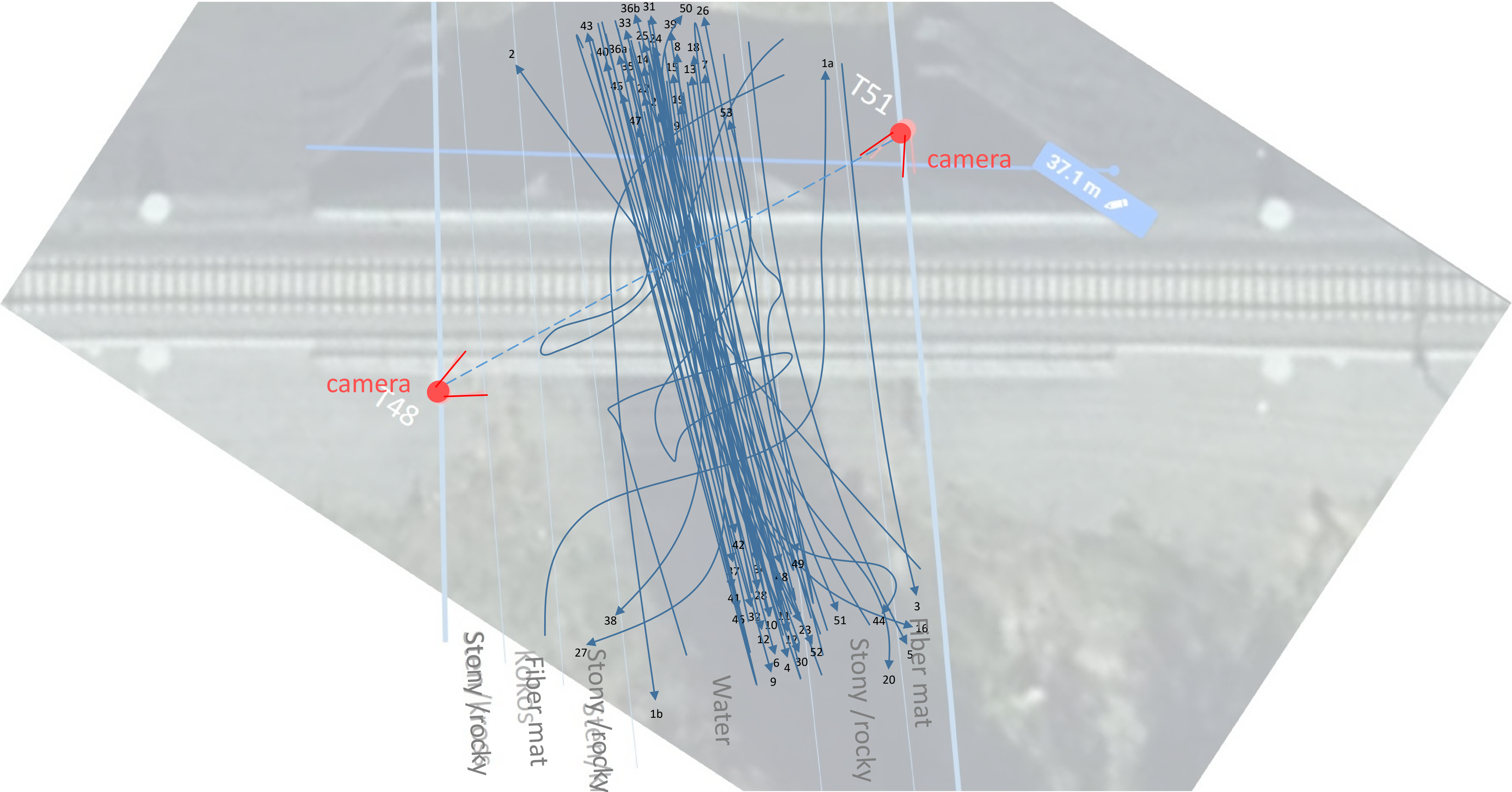

### Kvarnbäcken reindeer, snow free

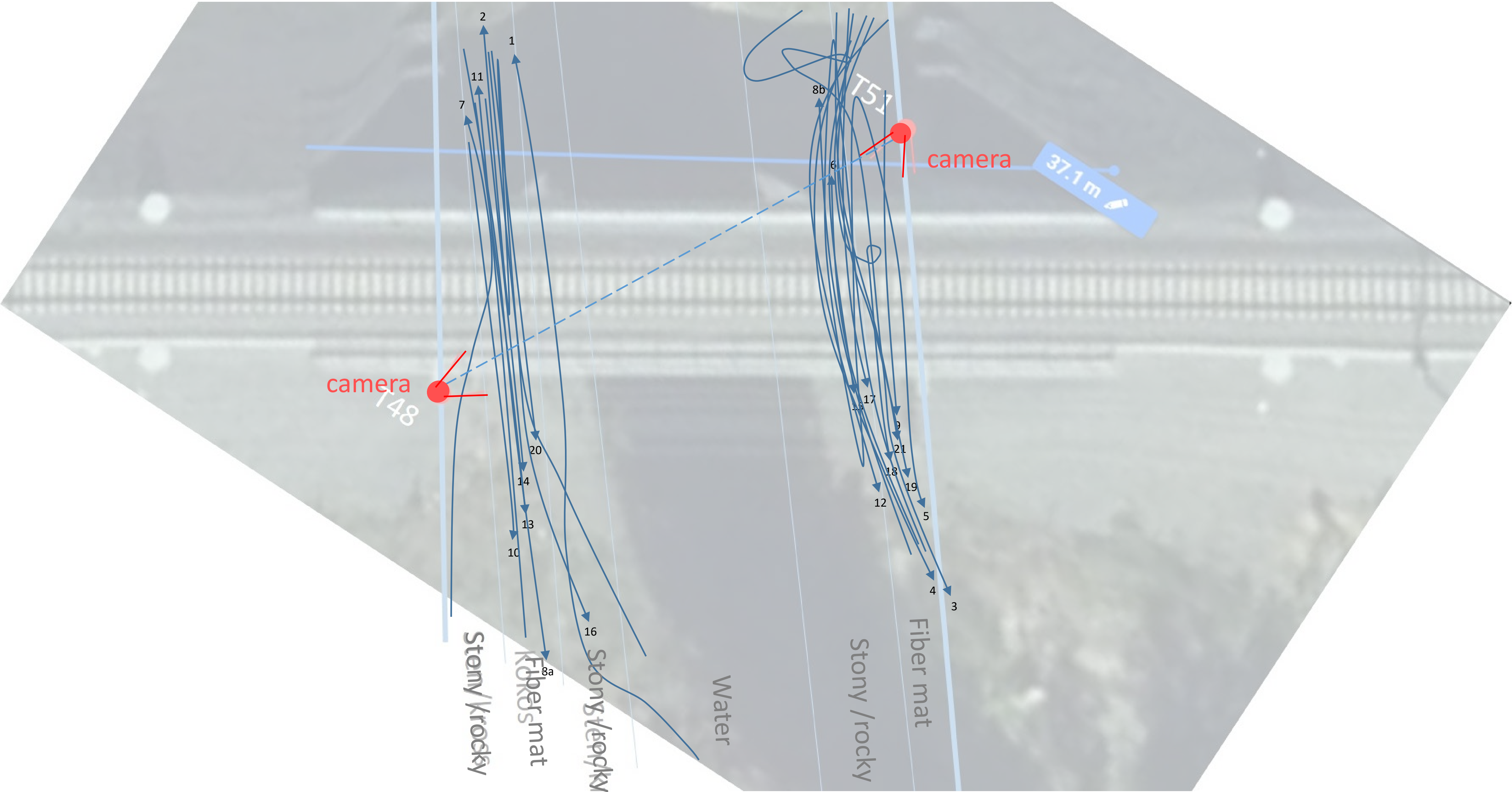

### Kvarnbäcken reindeer, snowy ground

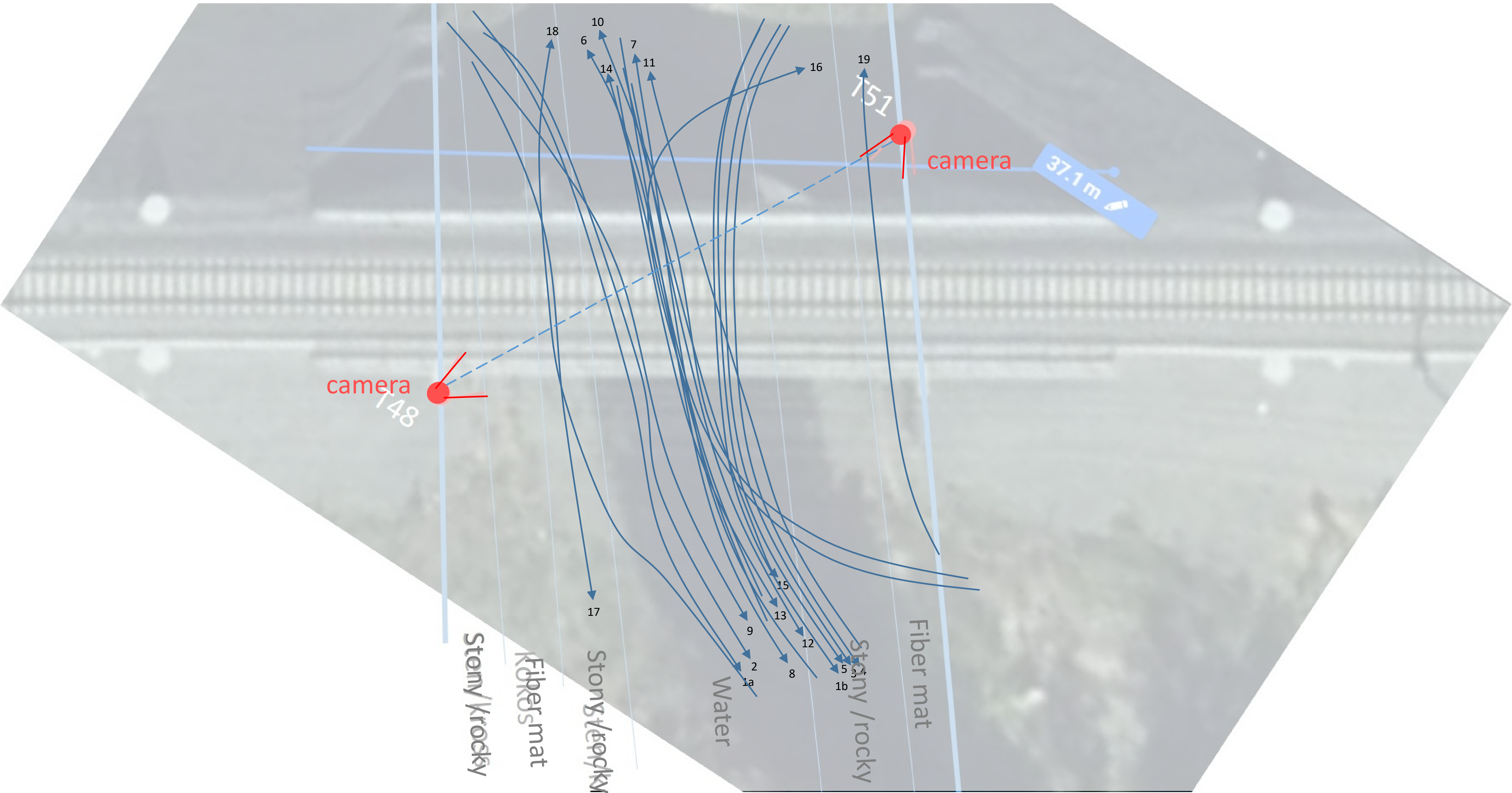

### Kvarnbäcken roe deer, snow free

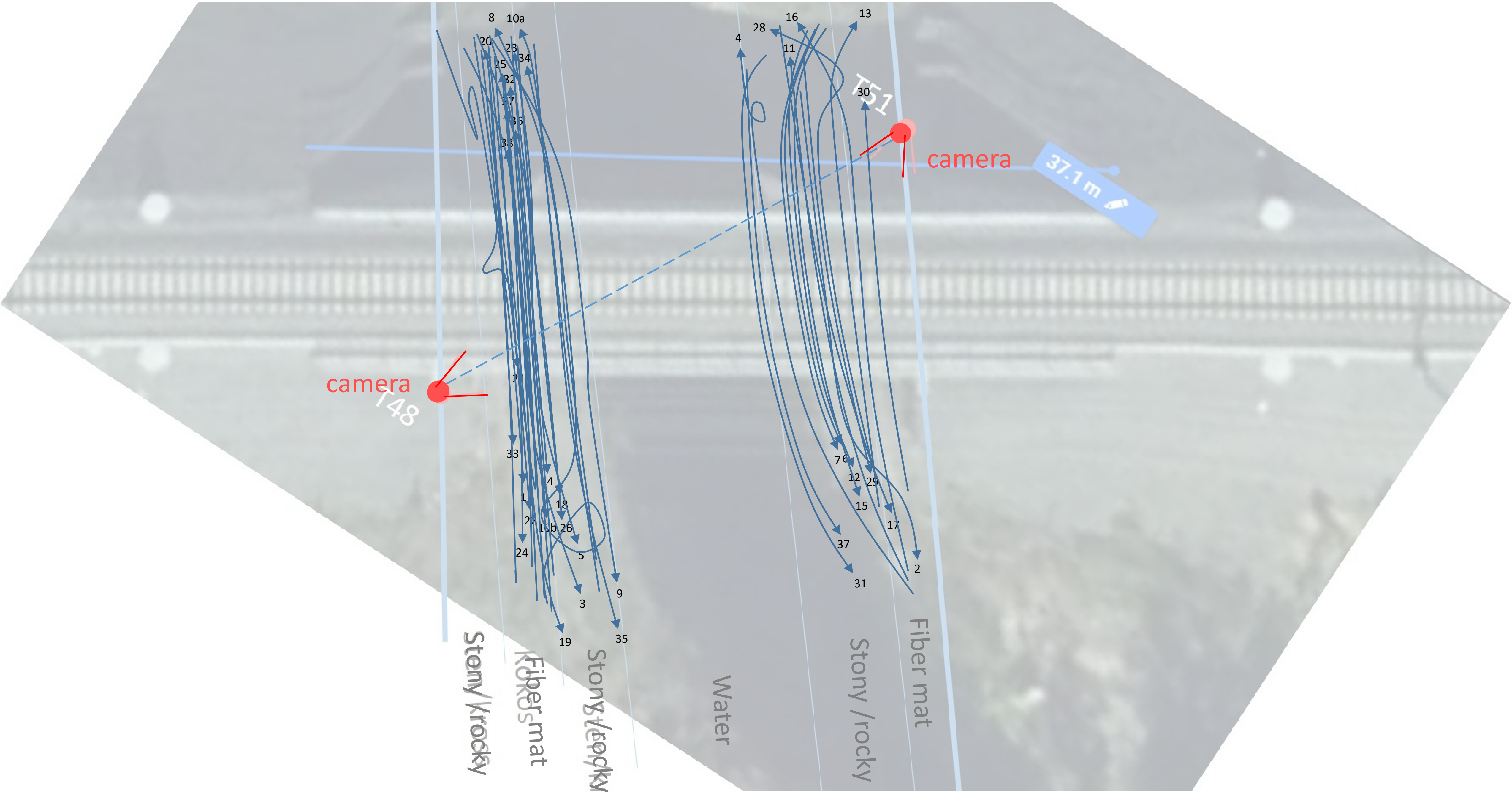

### Kvarnbäcken roe deer, snowy ground

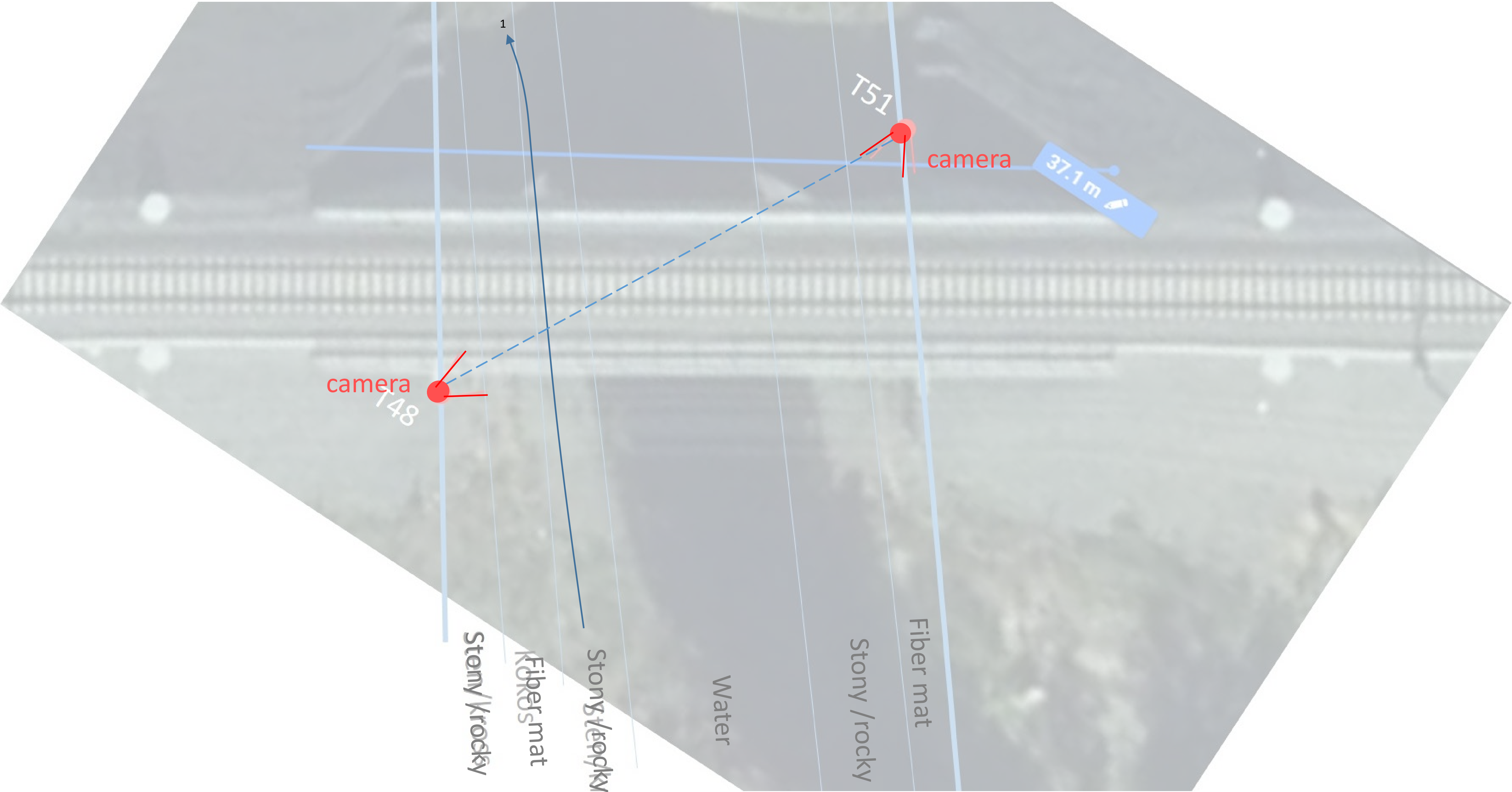

### Aavajoki moose, snow free

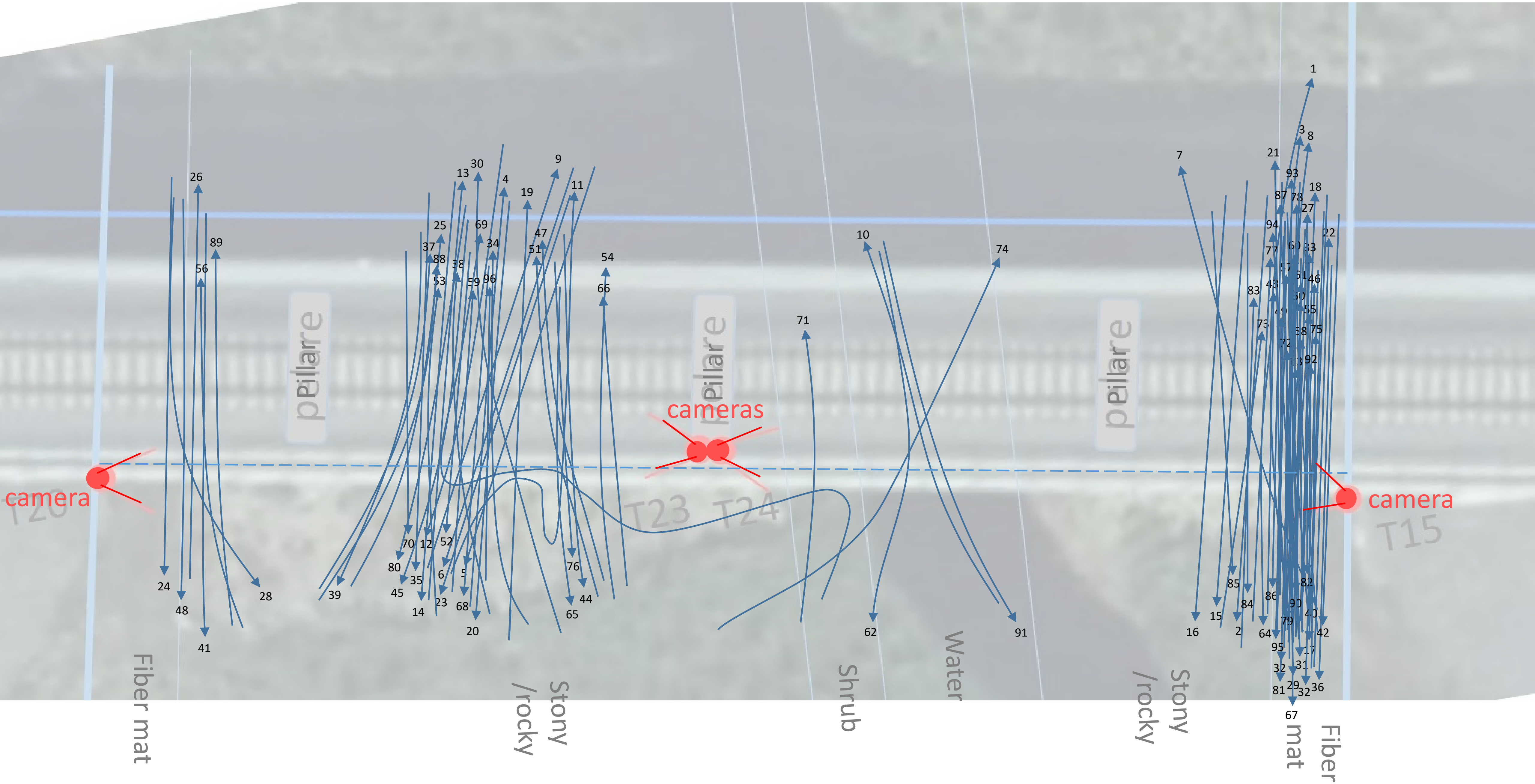

### Aavajoki moose, snowy ground

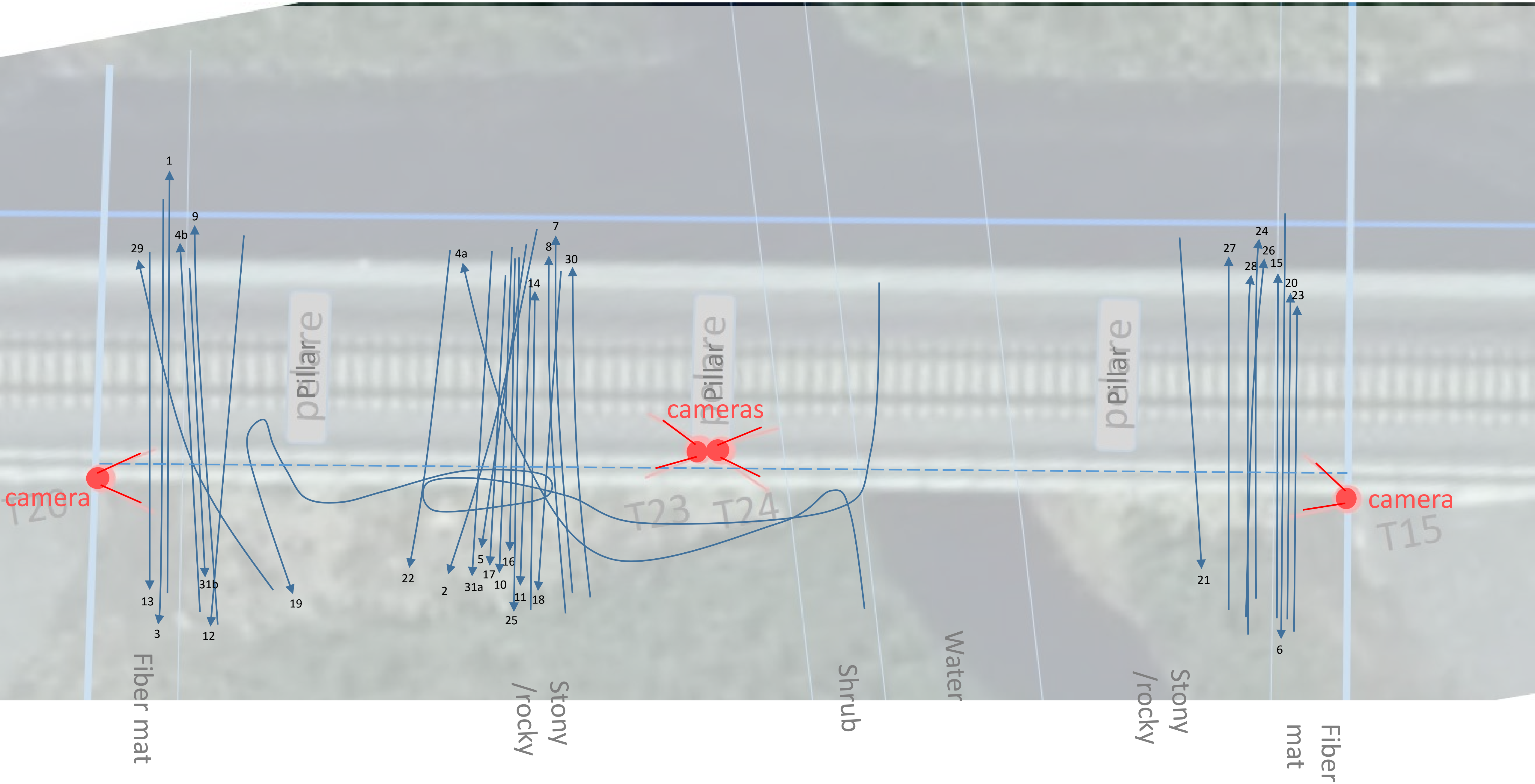

Aavajoki reindeer, snow free

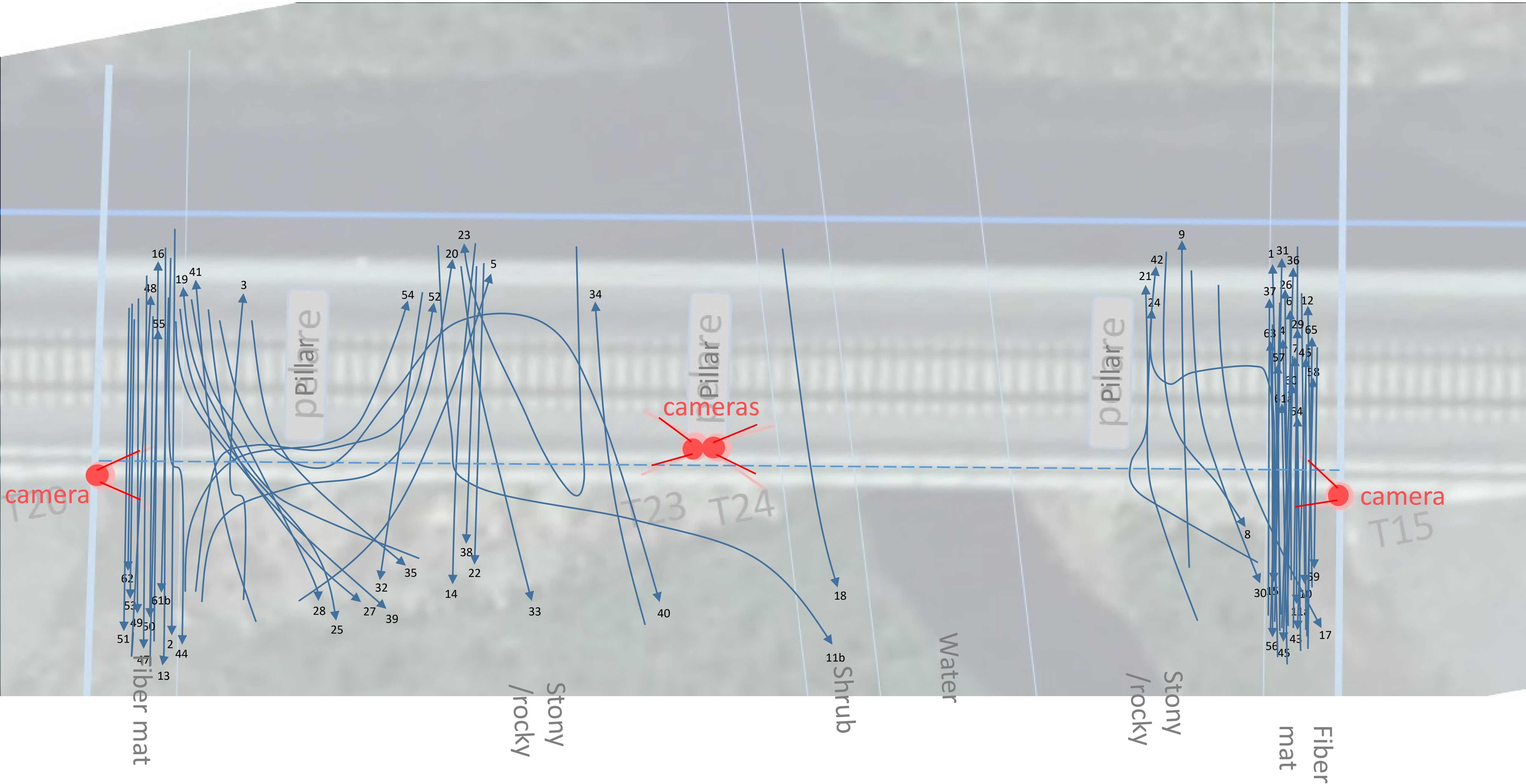

### Aavajoki reindeer, snowy ground

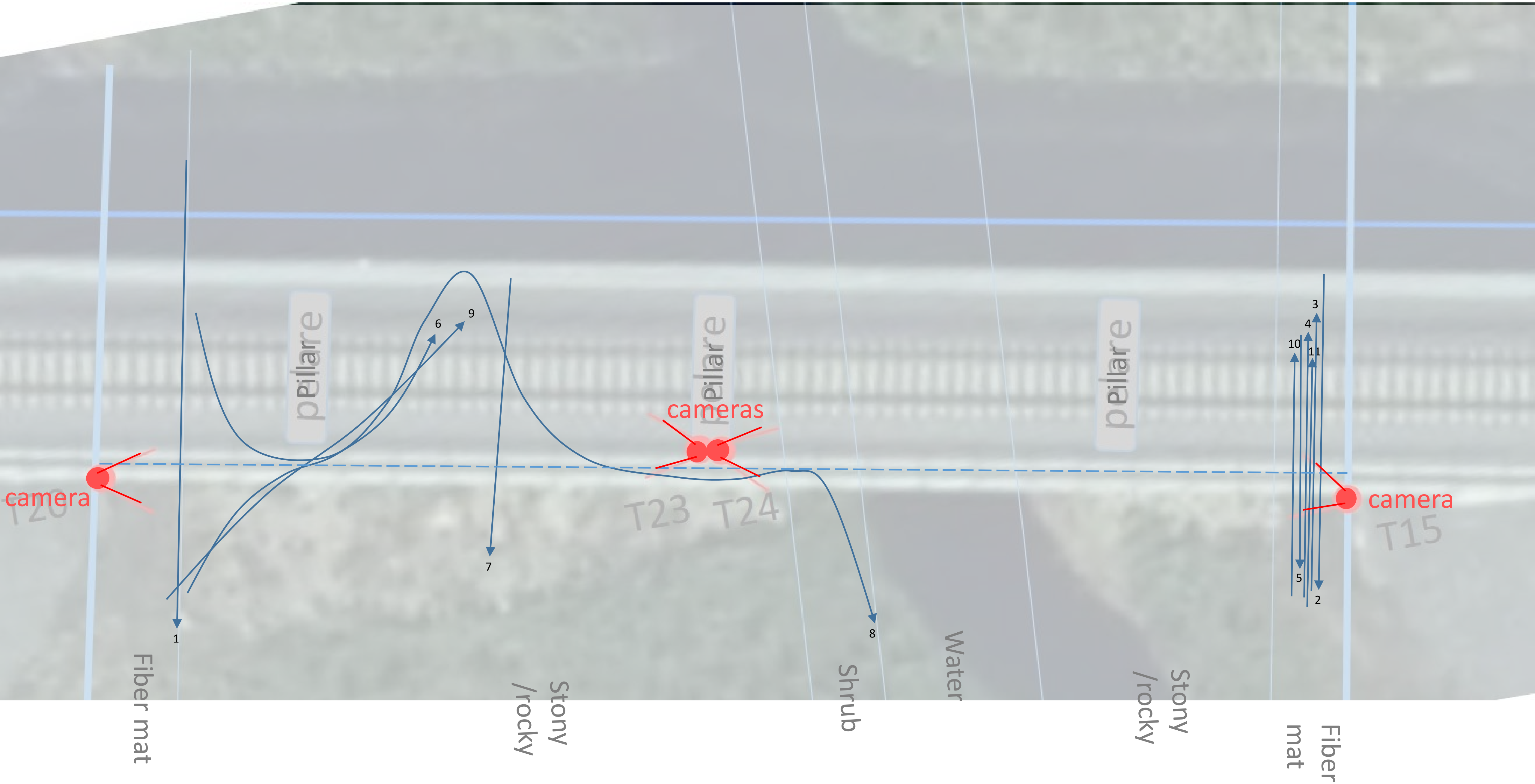

### Aavajoki roe deer, snow free

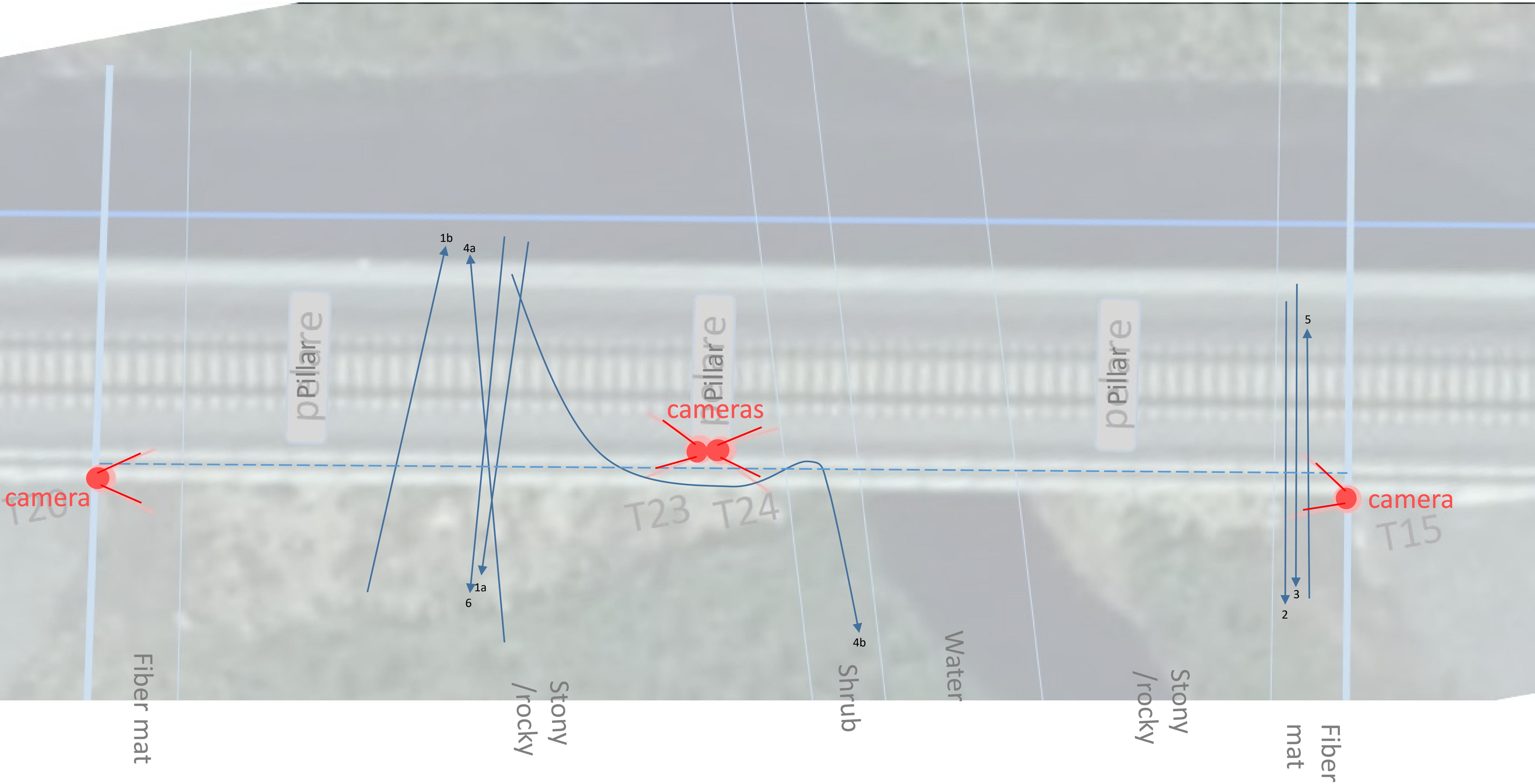

### Keräsjoki moose, snow free

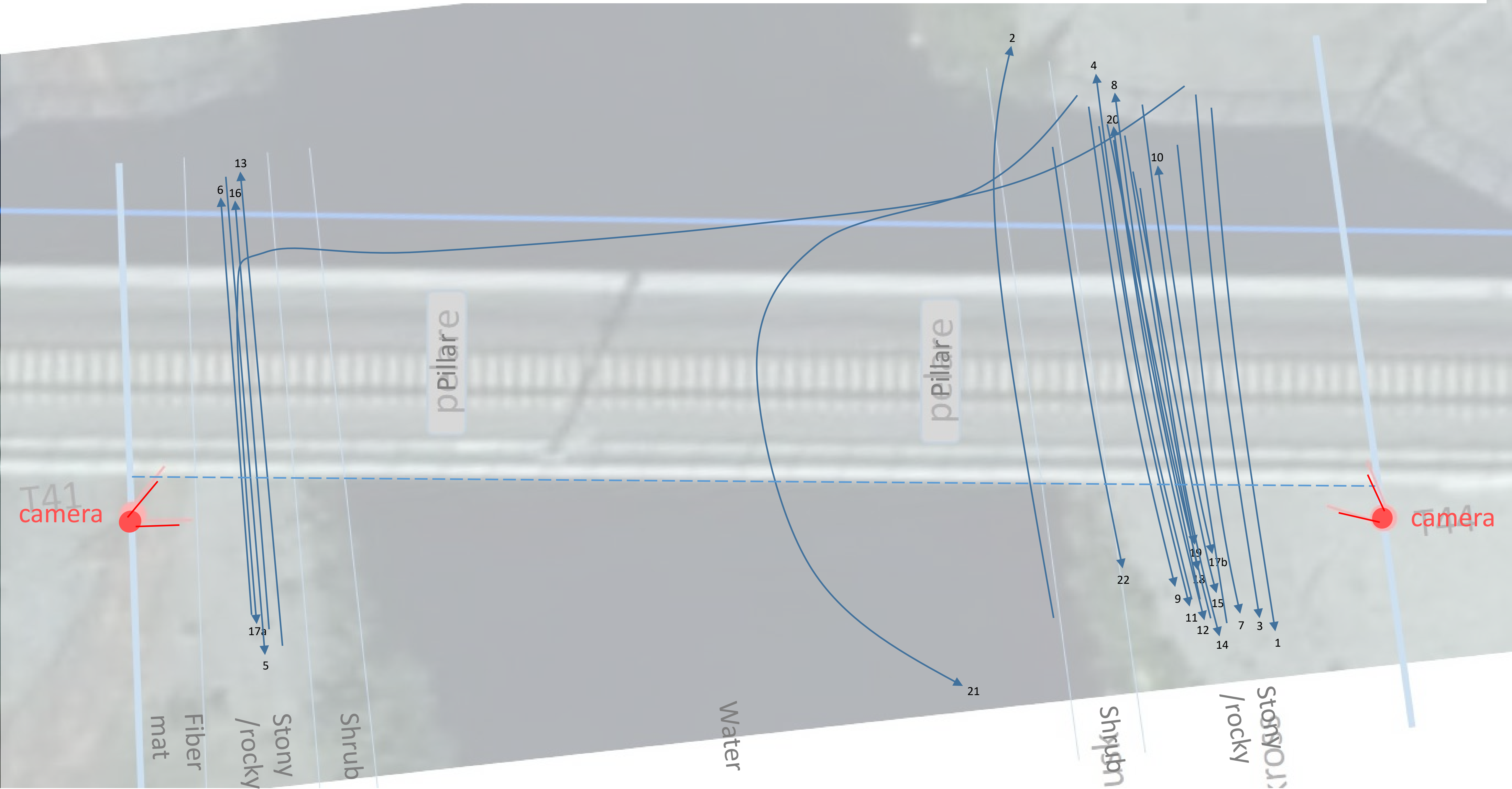

Keräsjoki moose, snowy ground

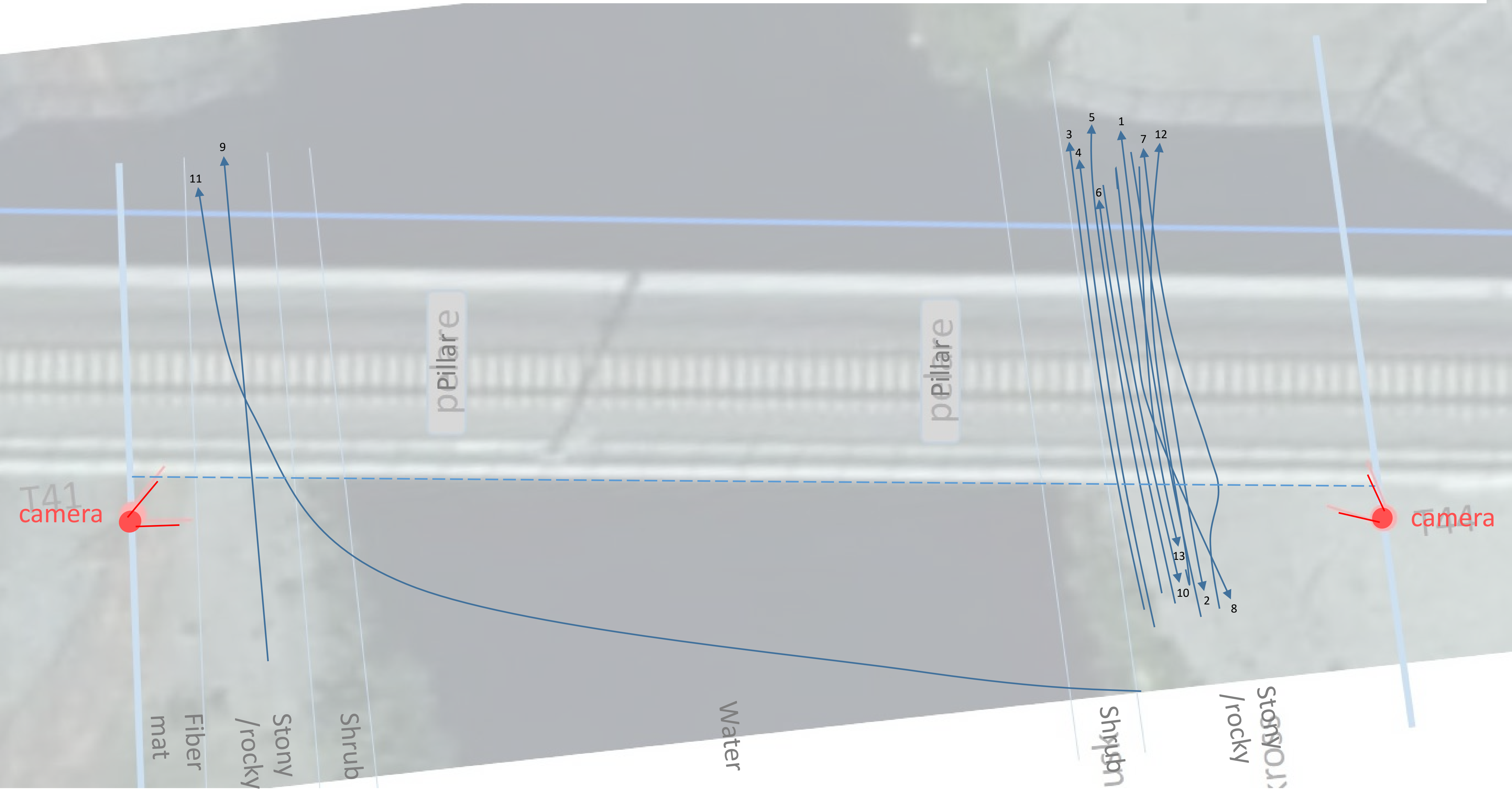

Keräsjoki reindeer, snow free

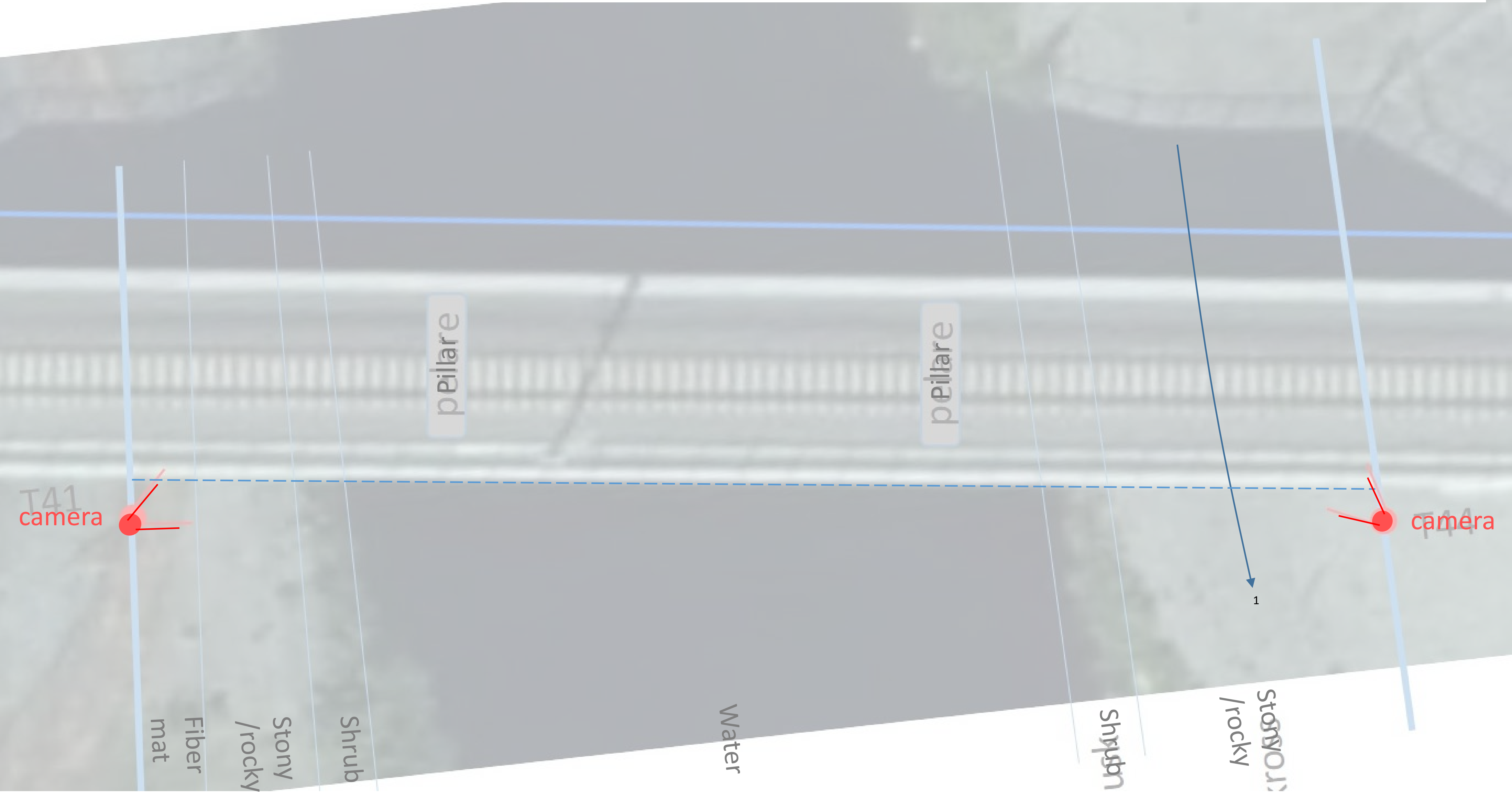

### Keräsjoki reindeer, snowy ground

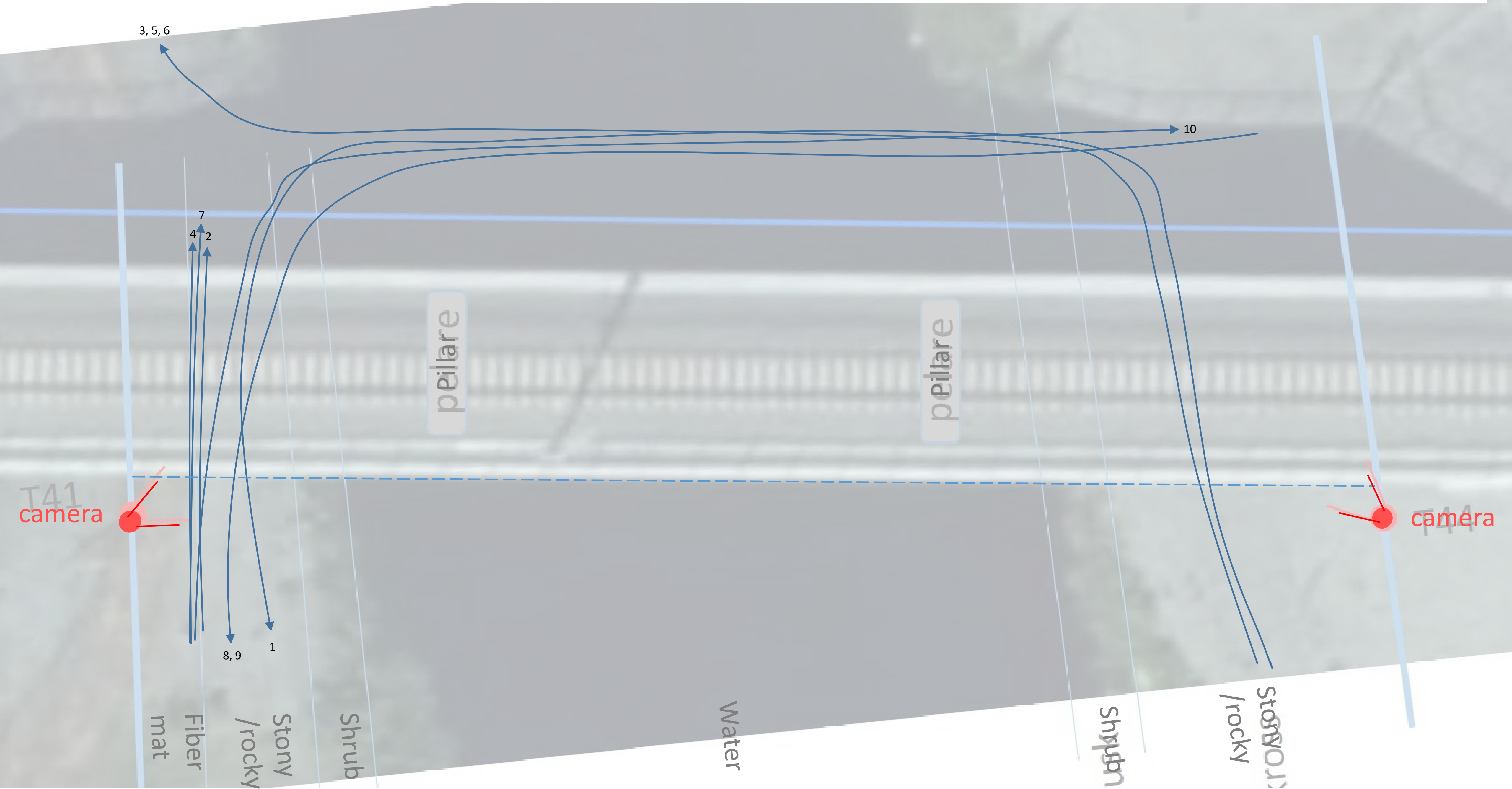

### Keräsjoki roe deer, snow free

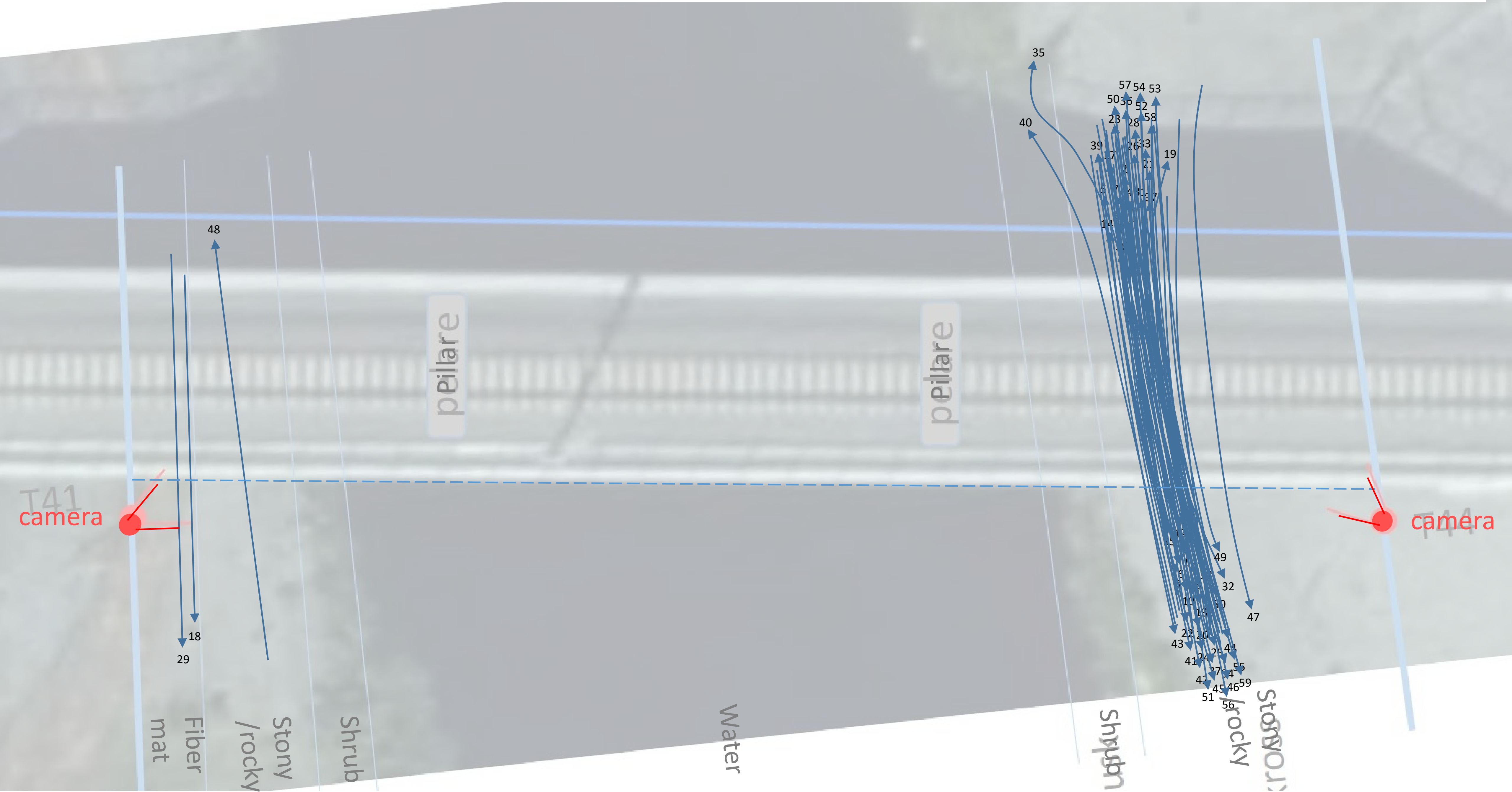

### Keräsjoki roe deer, snowy ground

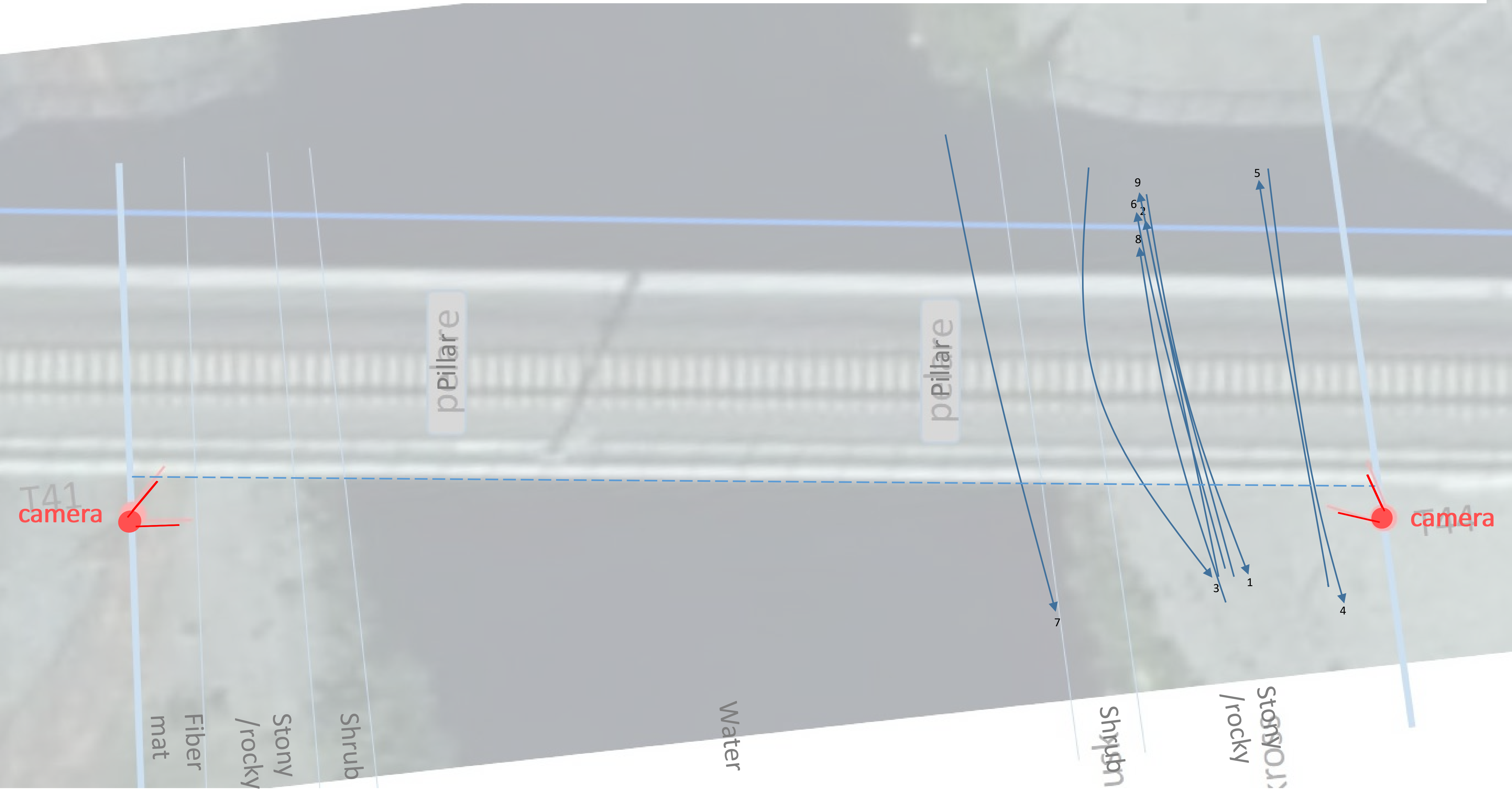

### Sangijärvi moose, snow free

Only trajectories no. 7-11 were included in analyses

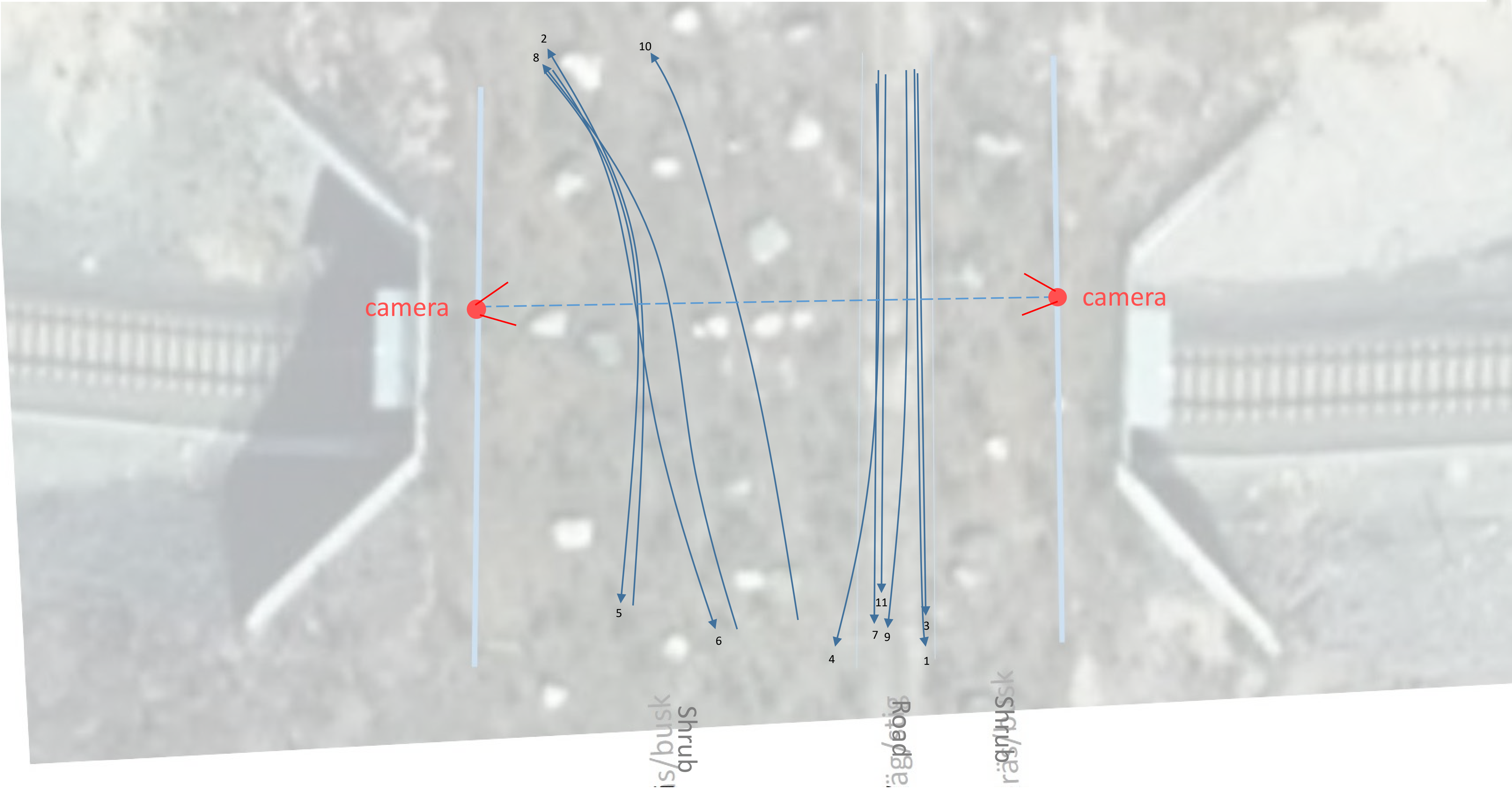

### Sangijärvi moose, snowy ground

Only trajectories no. 9-28 were included in analyses

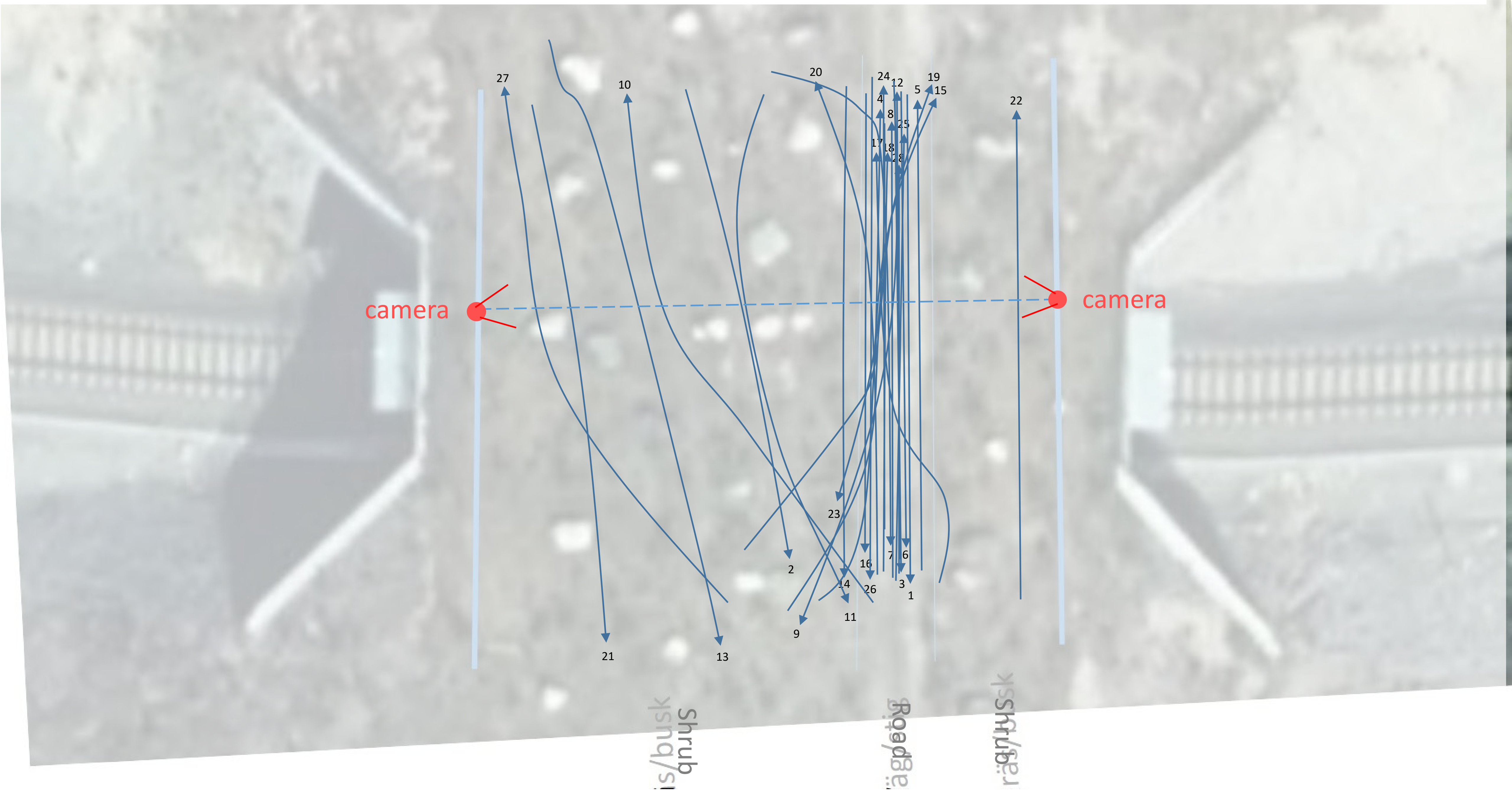

### Sangijärvi reindeer, snow free

Only trajectories no. 69-127 were included in analyses

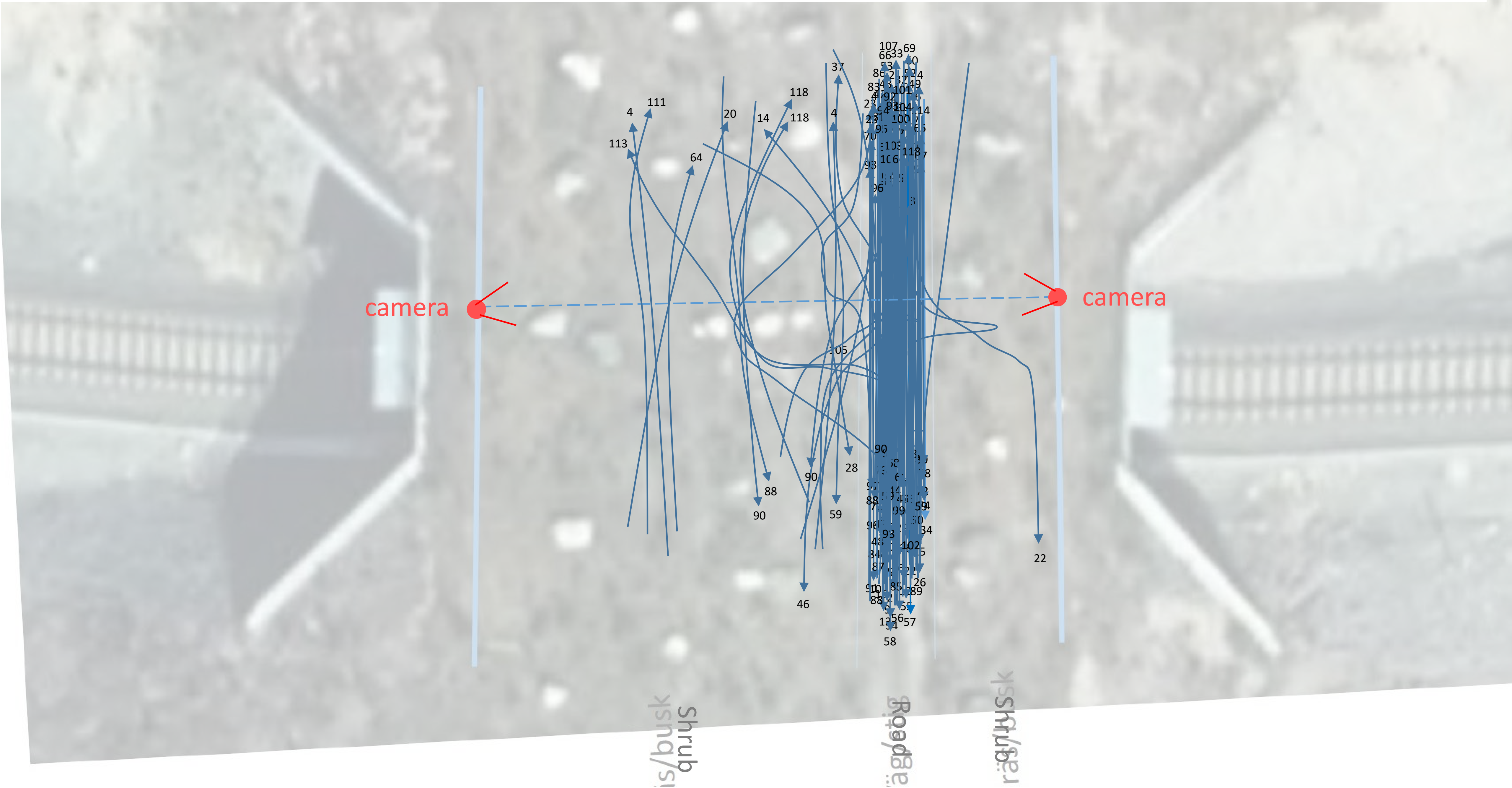

### Sangijärvi reindeer, snowy ground

Only trajectories no. 18-99 were included in analyses

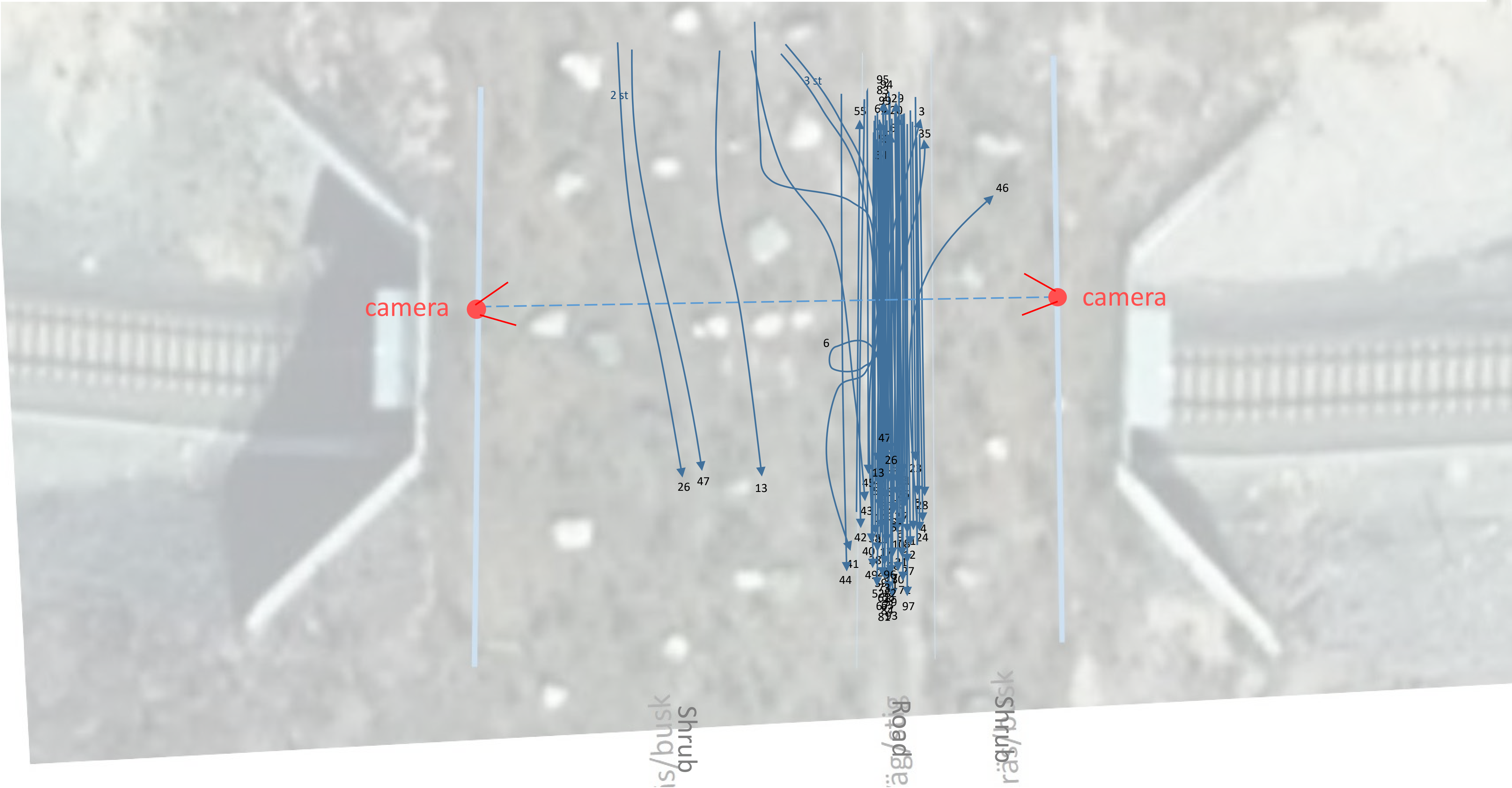

### Sangijärvi roe deer, snow free

Only trajectories no. 15-32 were included in analyses

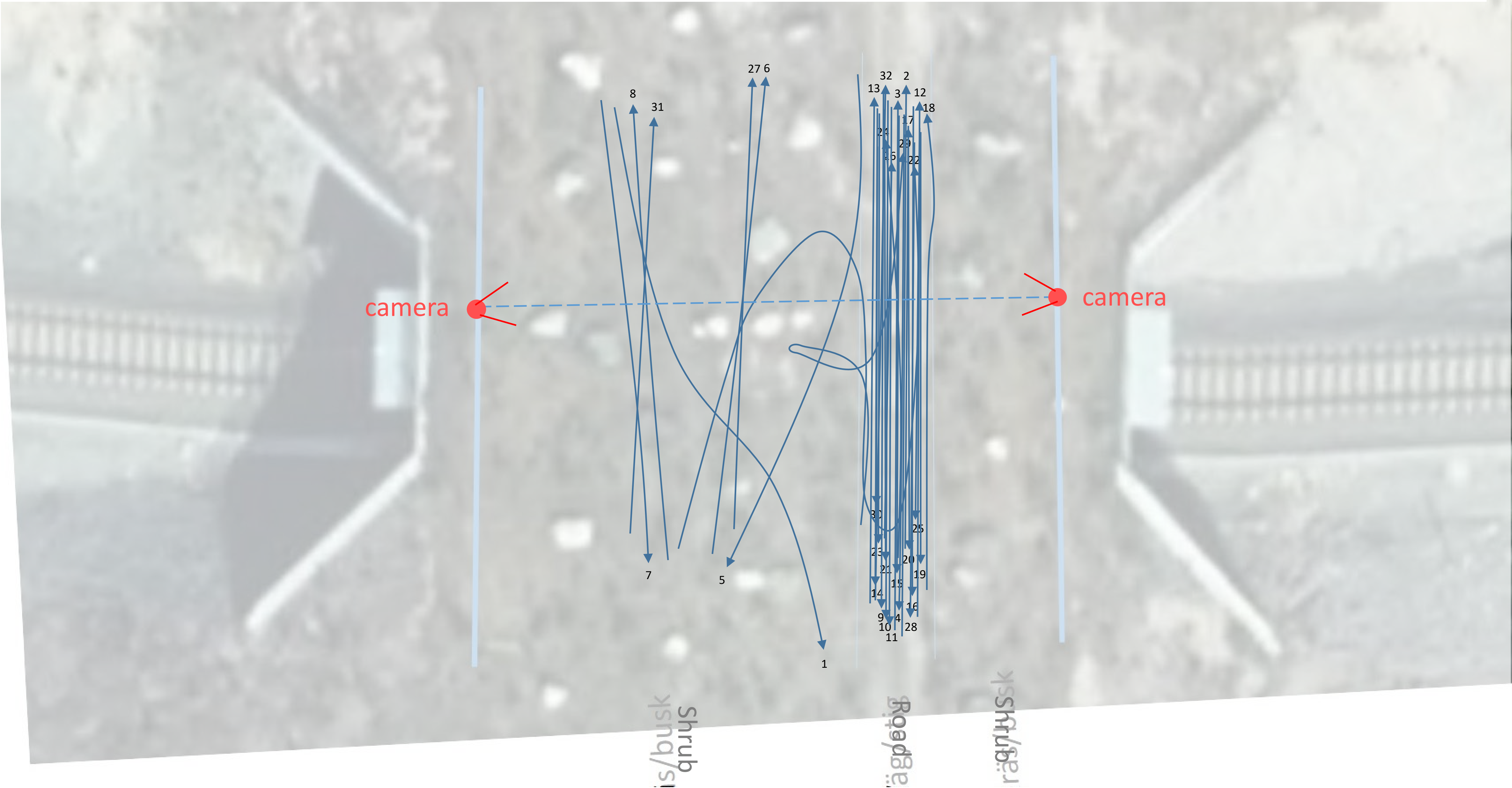

### Sangijärvi roe deer, snowy ground

Only trajectories no. 2-3 were included in analyses

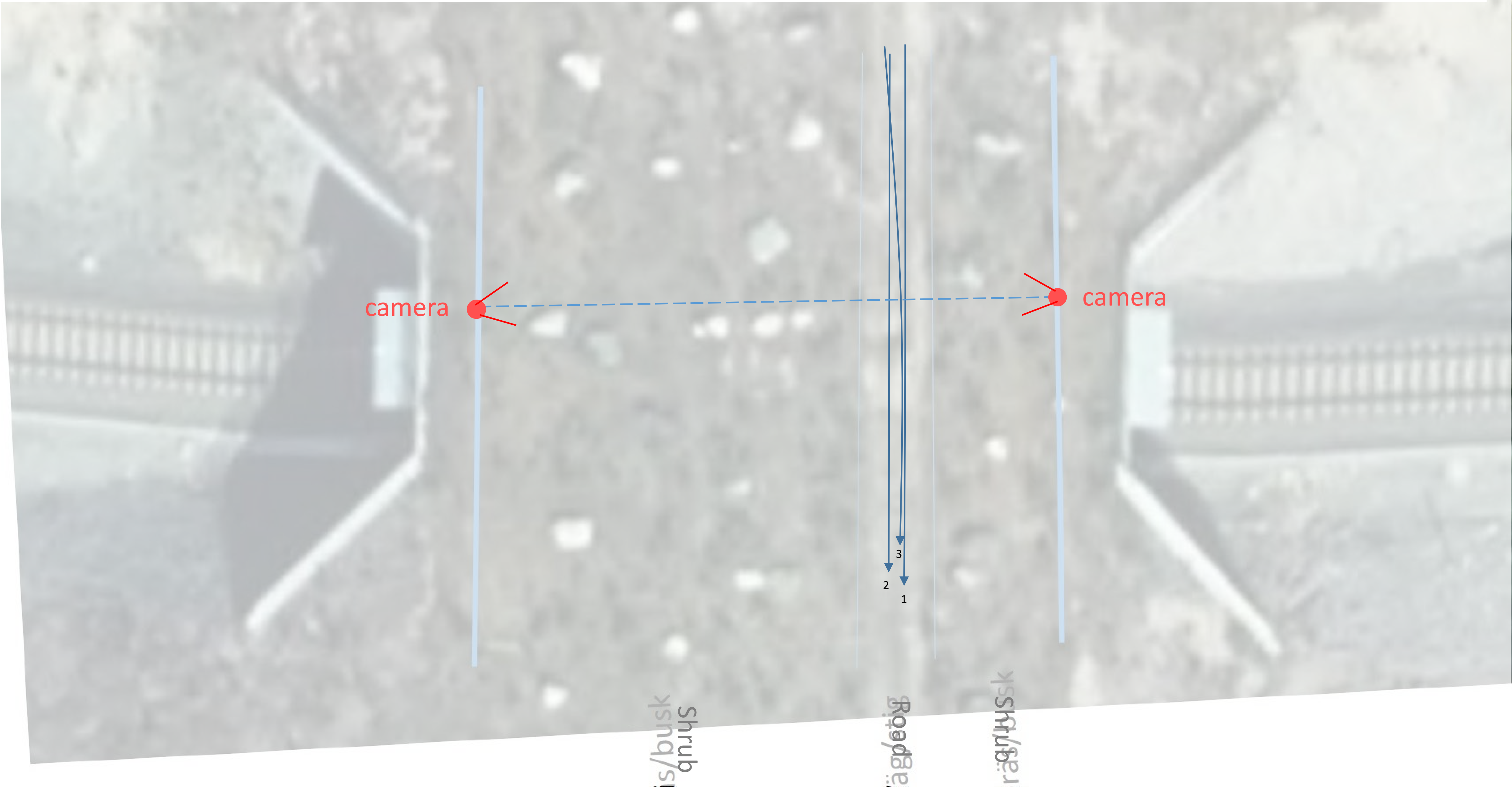

### Sammatti moose, snow free

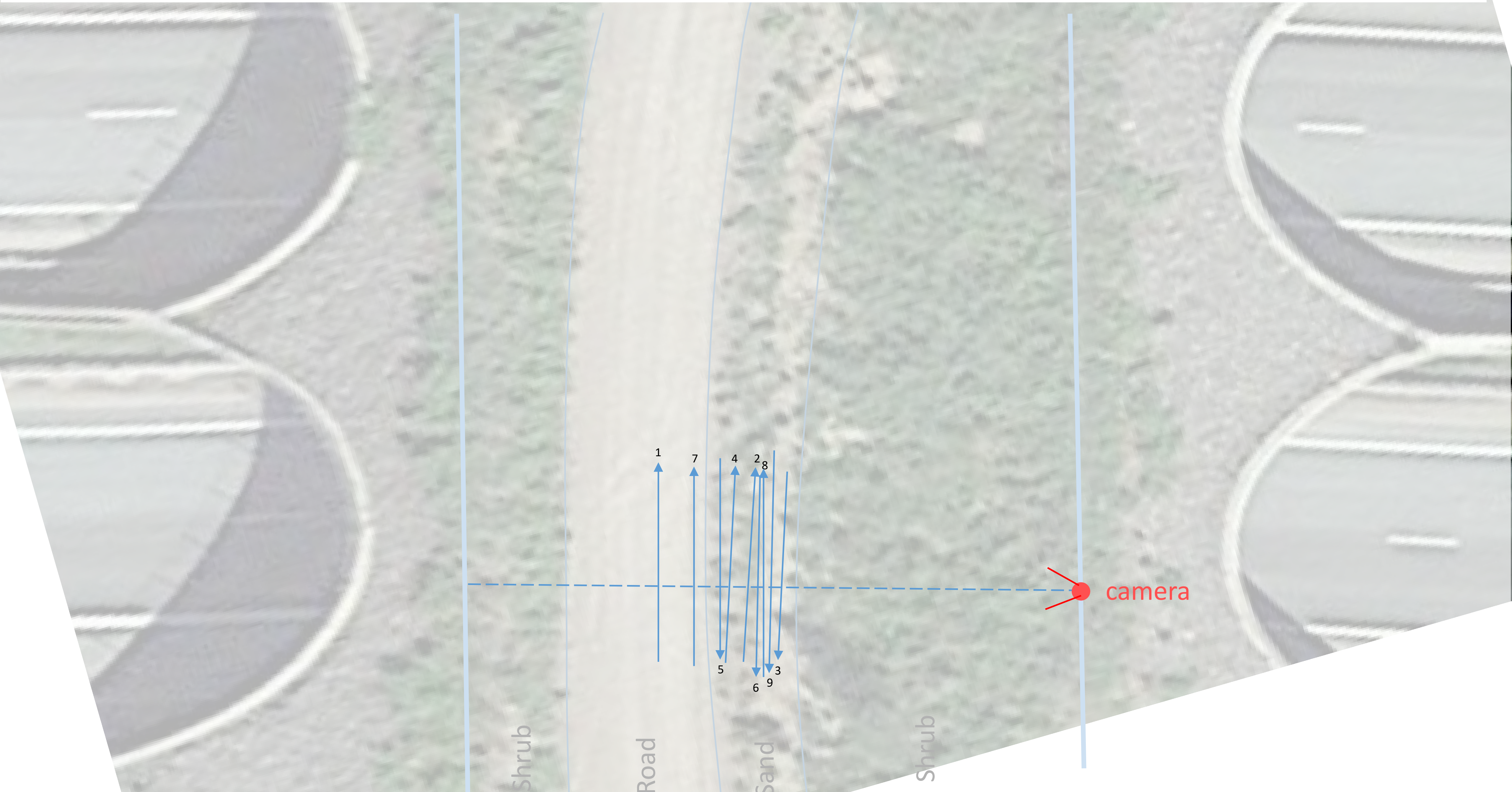

### Sammatti roe deer, snow free

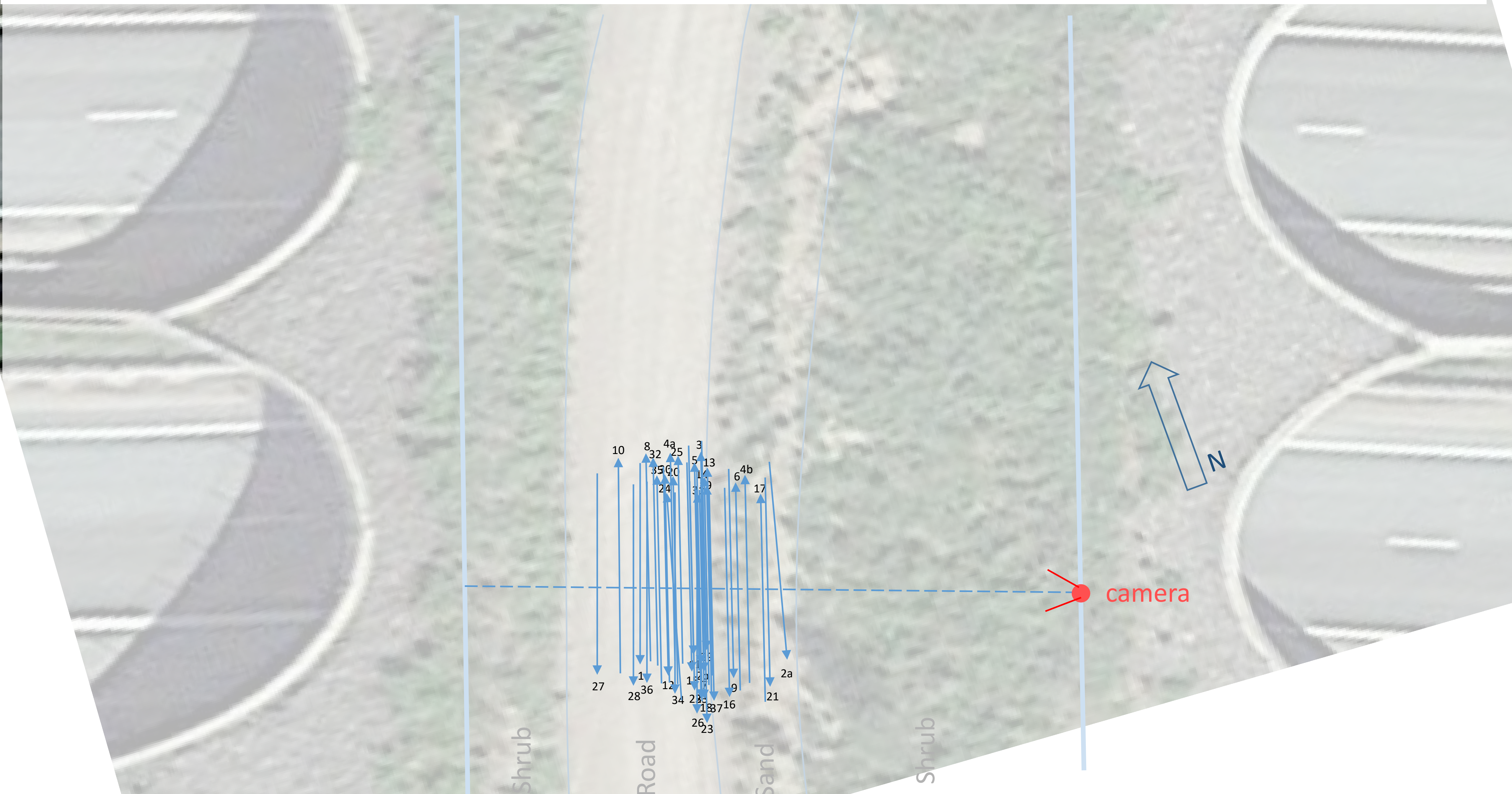

### Sammatti white-tailed deer, snow free

Subsample with 110 of 172 events

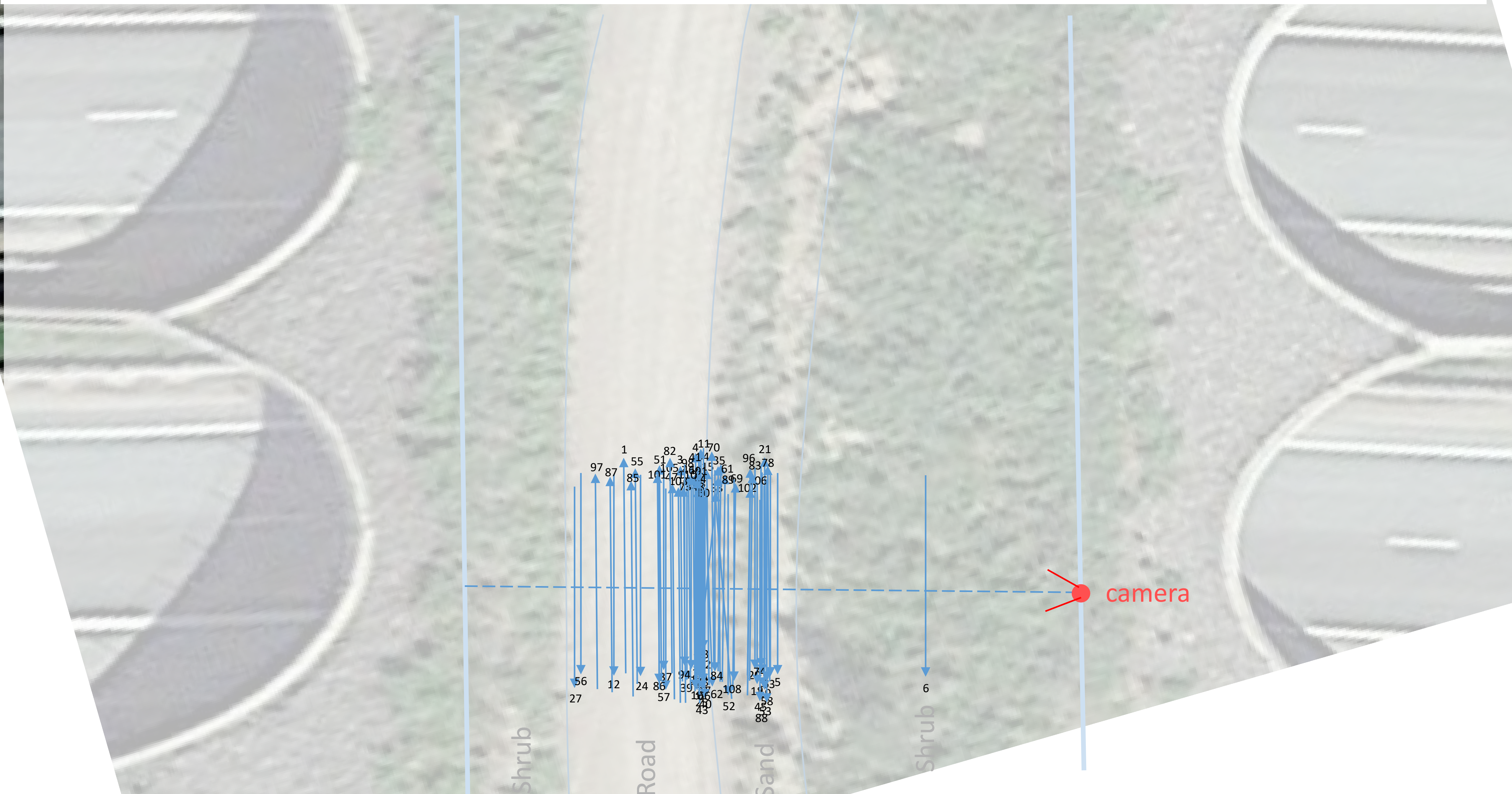

### Sammatti white-tailed deer, snowy ground

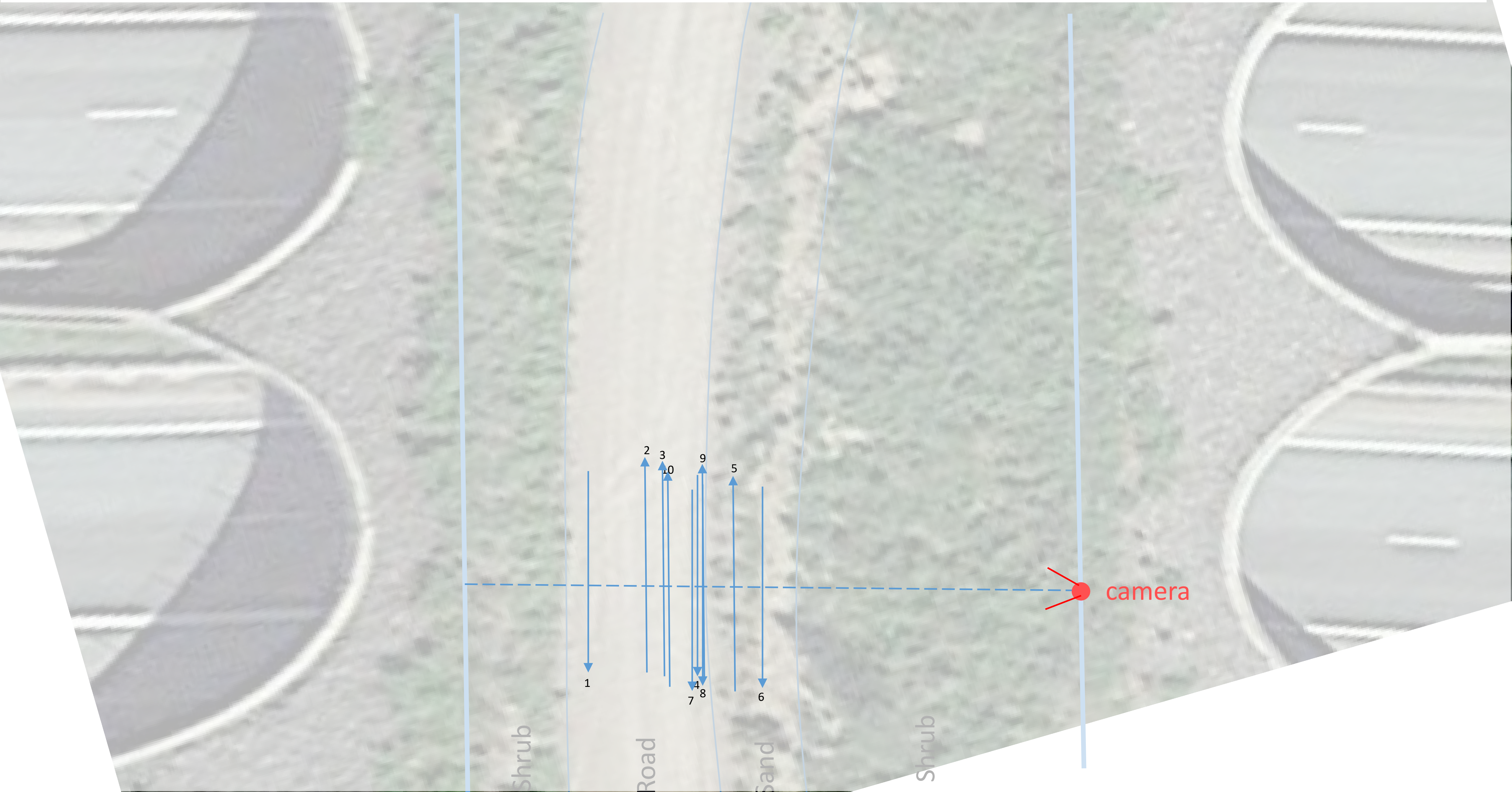

### Loviisa 2, moose, snow free

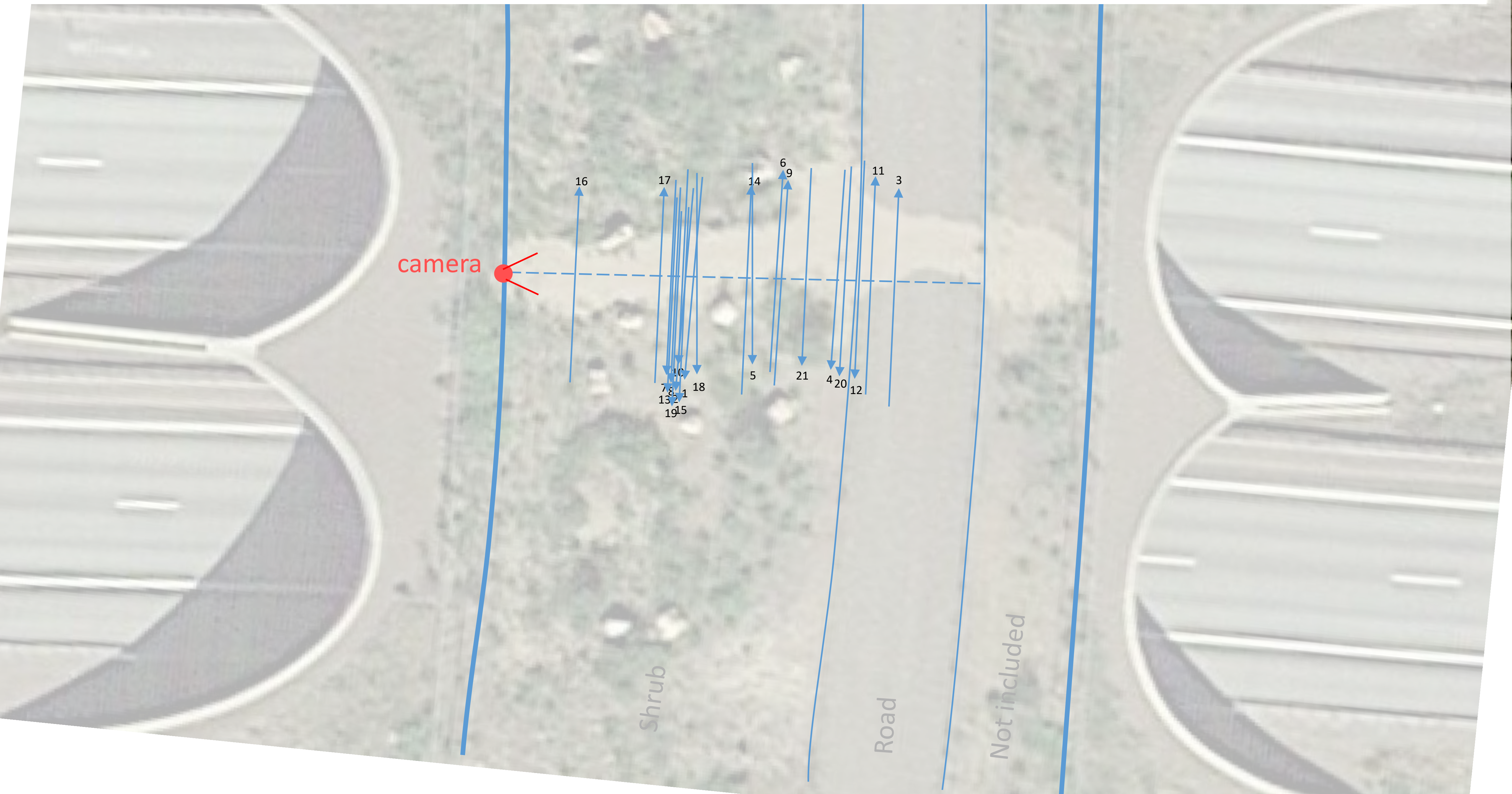

### Loviisa 2, moose, snowy ground

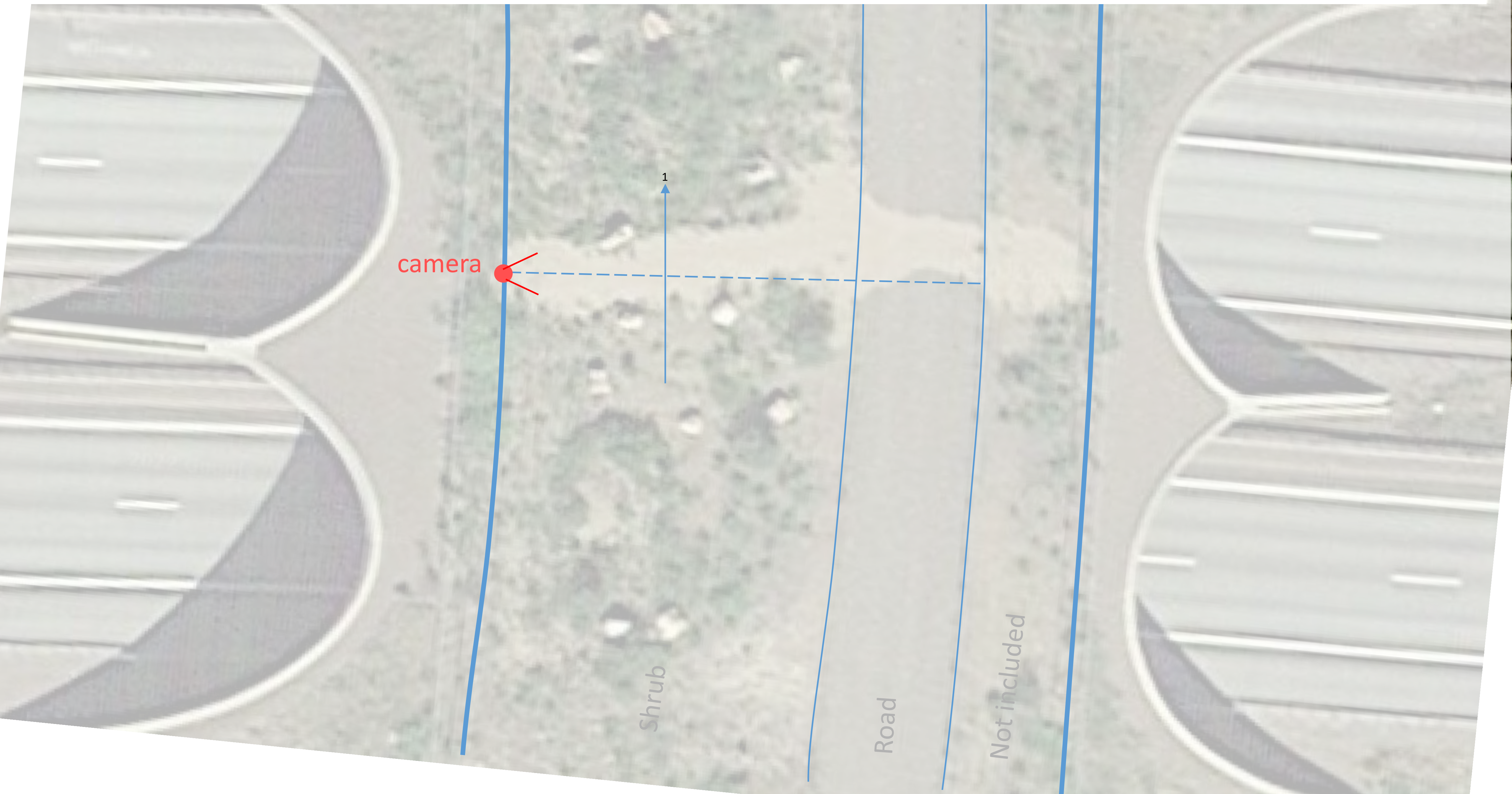

### Loviisa 2, roe deer, snow free

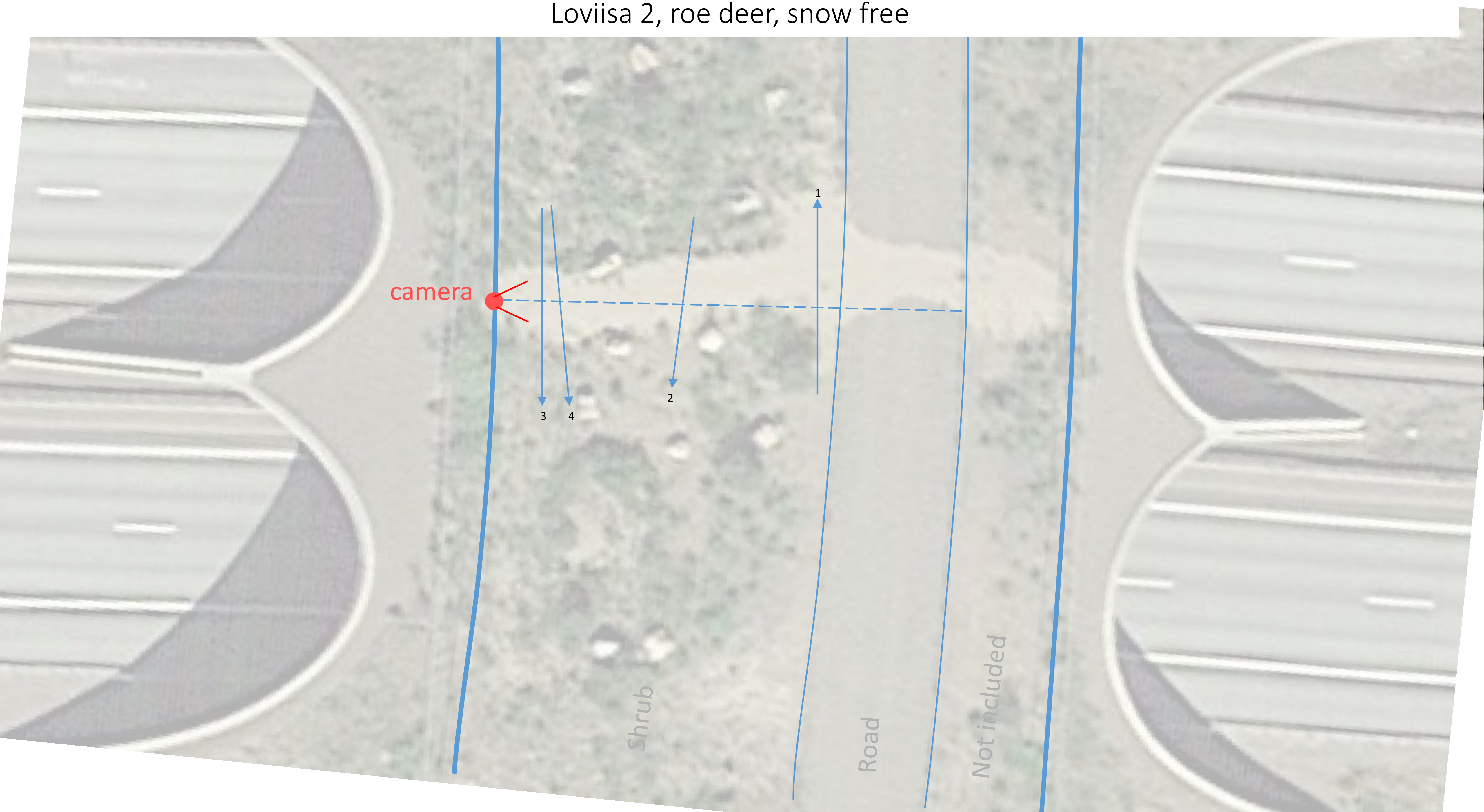

### Loviisa 2, white-tailed deer, snow free

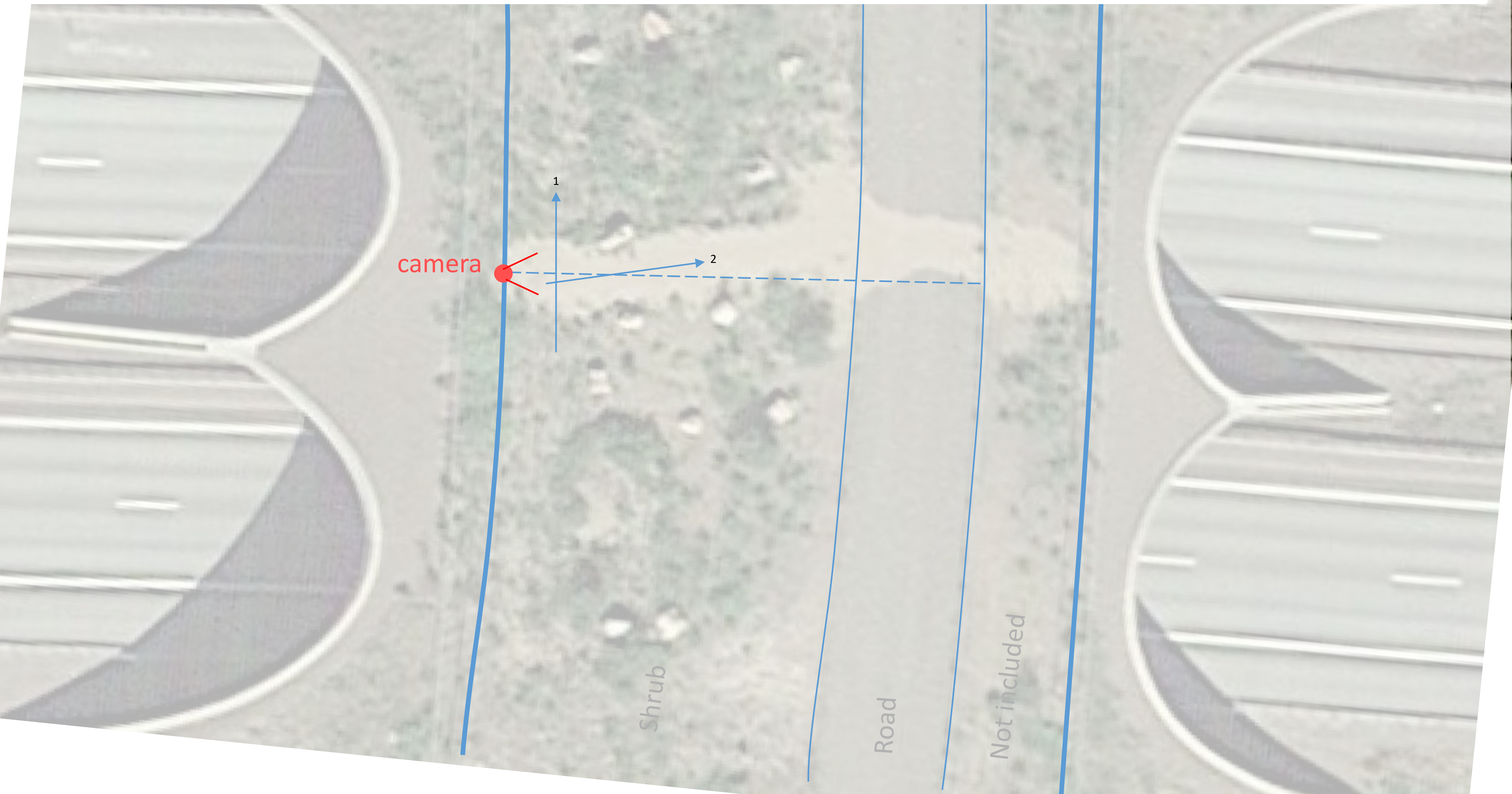

### Loviisa 2, white-tailed deer, snowy ground

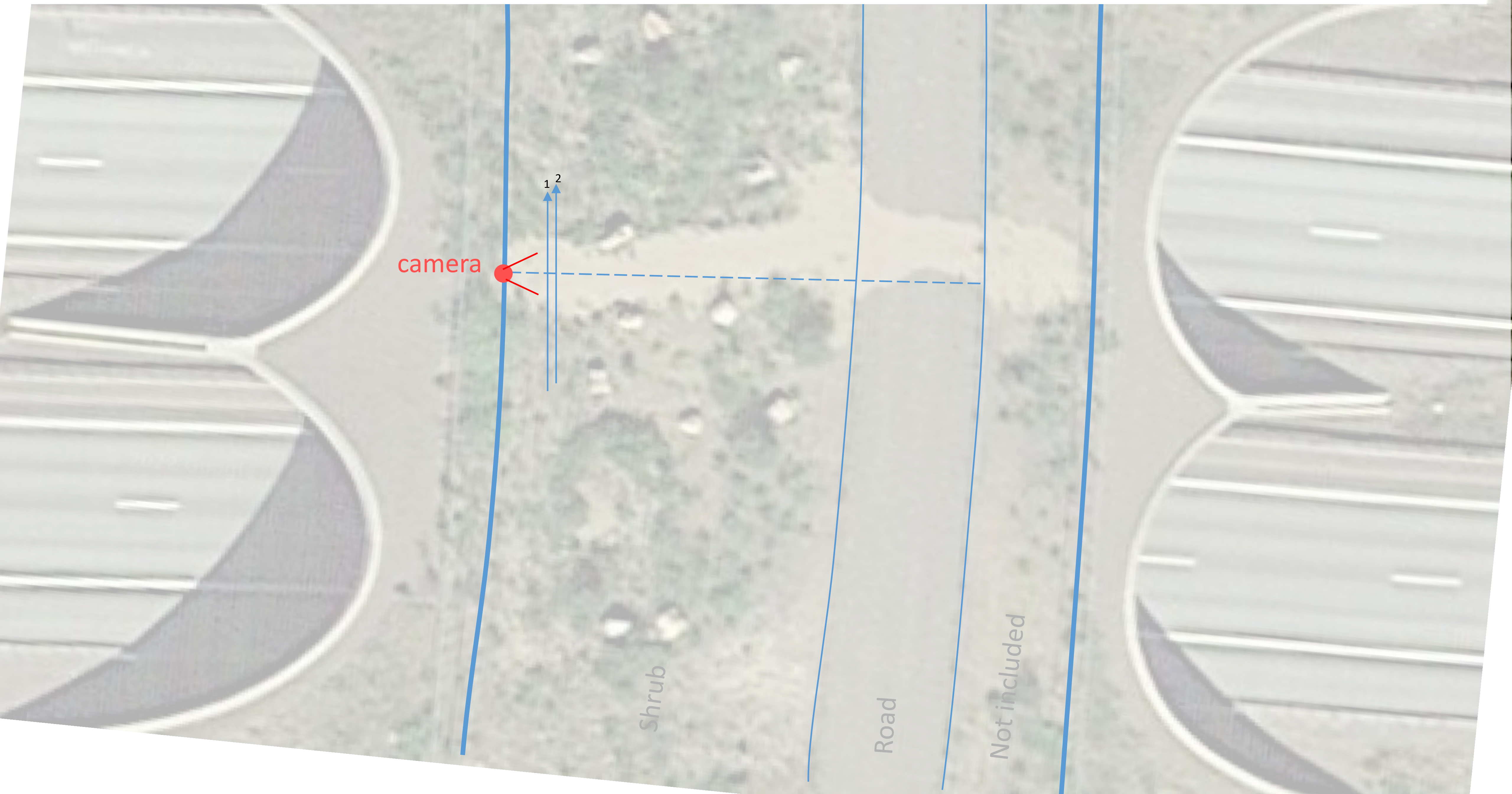

### Loviisa 2, wildboar, snow free

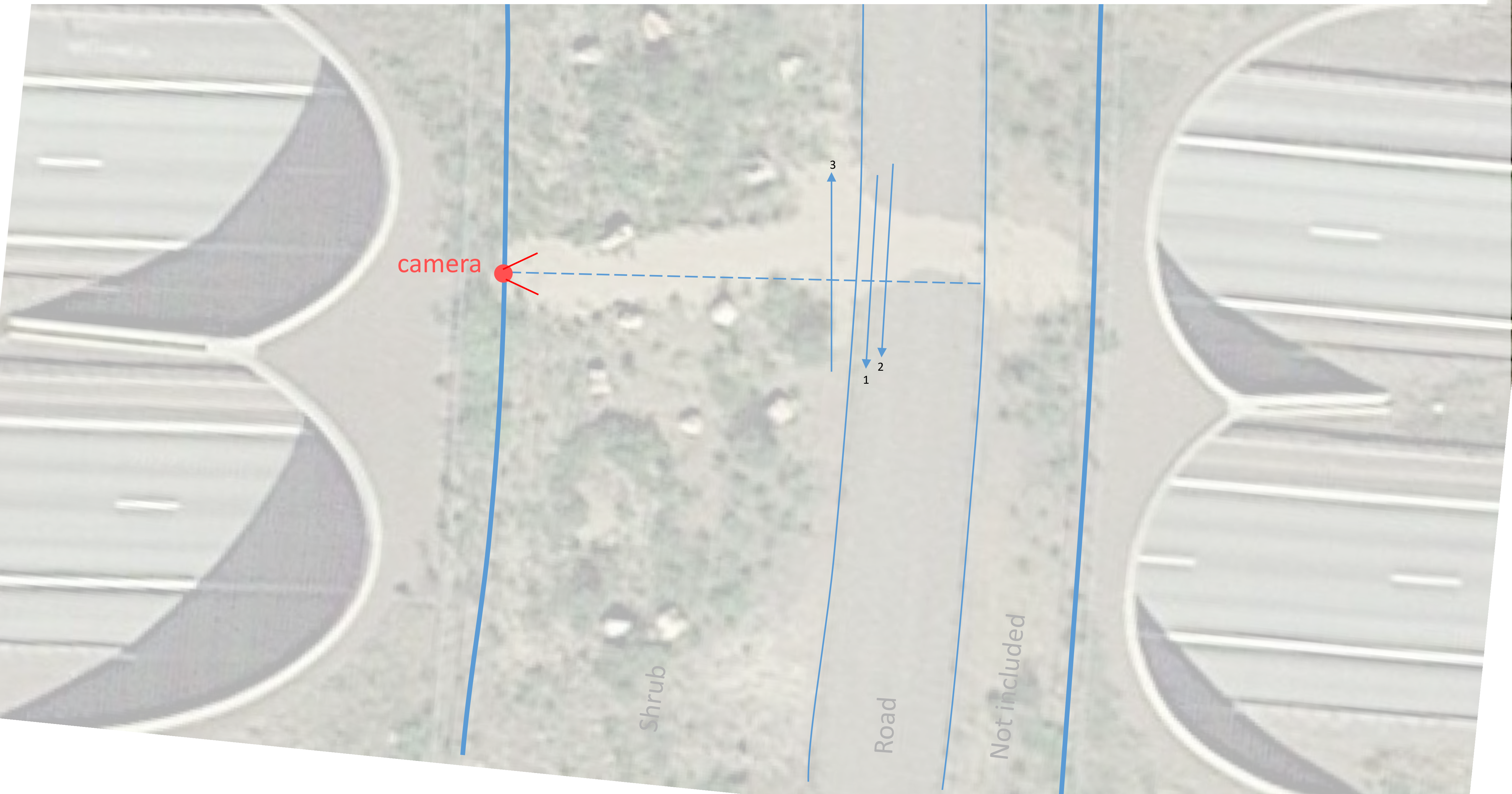

### Kärmekorpi, moose, snow free

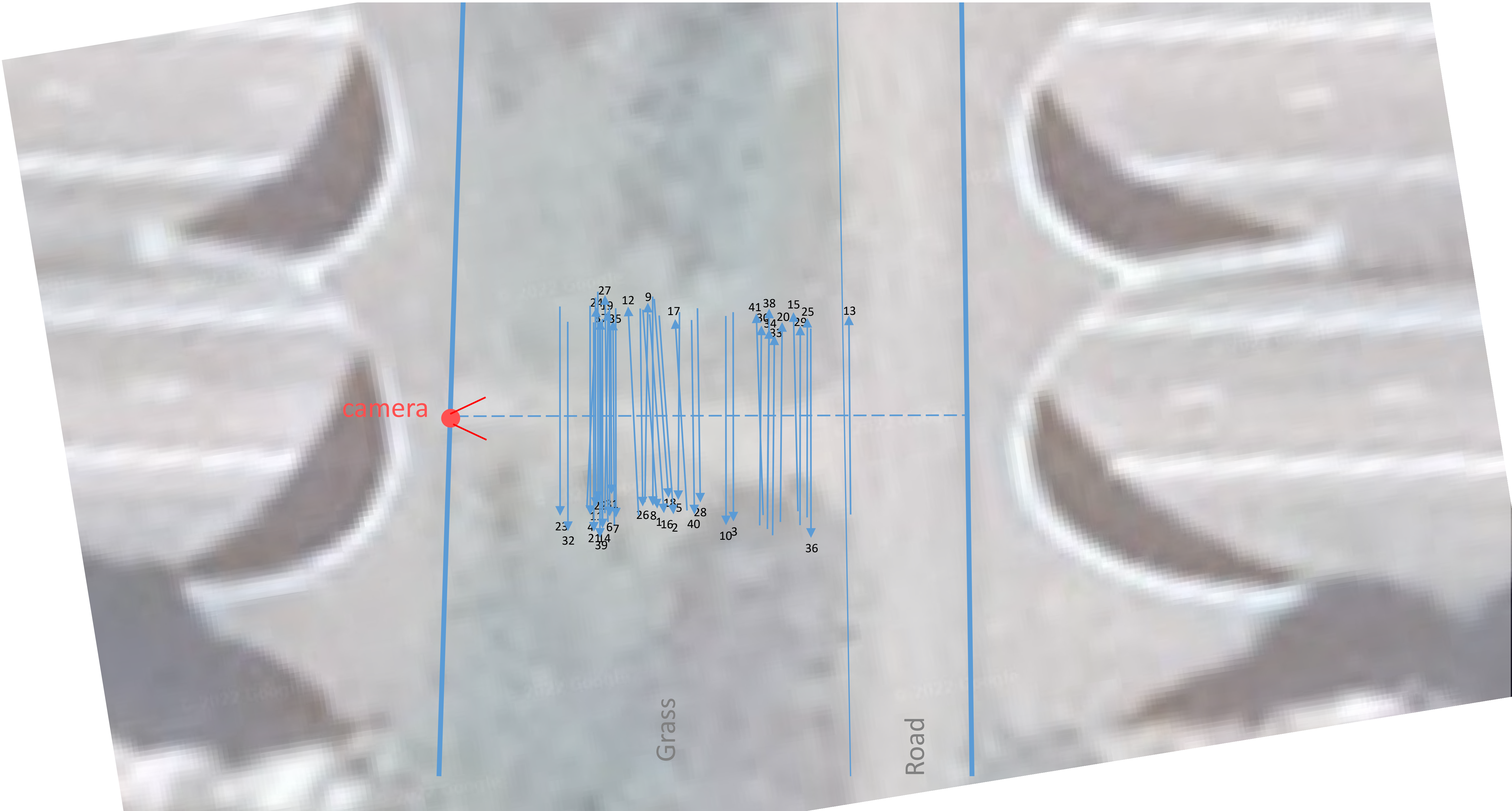

### Kärmekorpi, moose, snowy ground

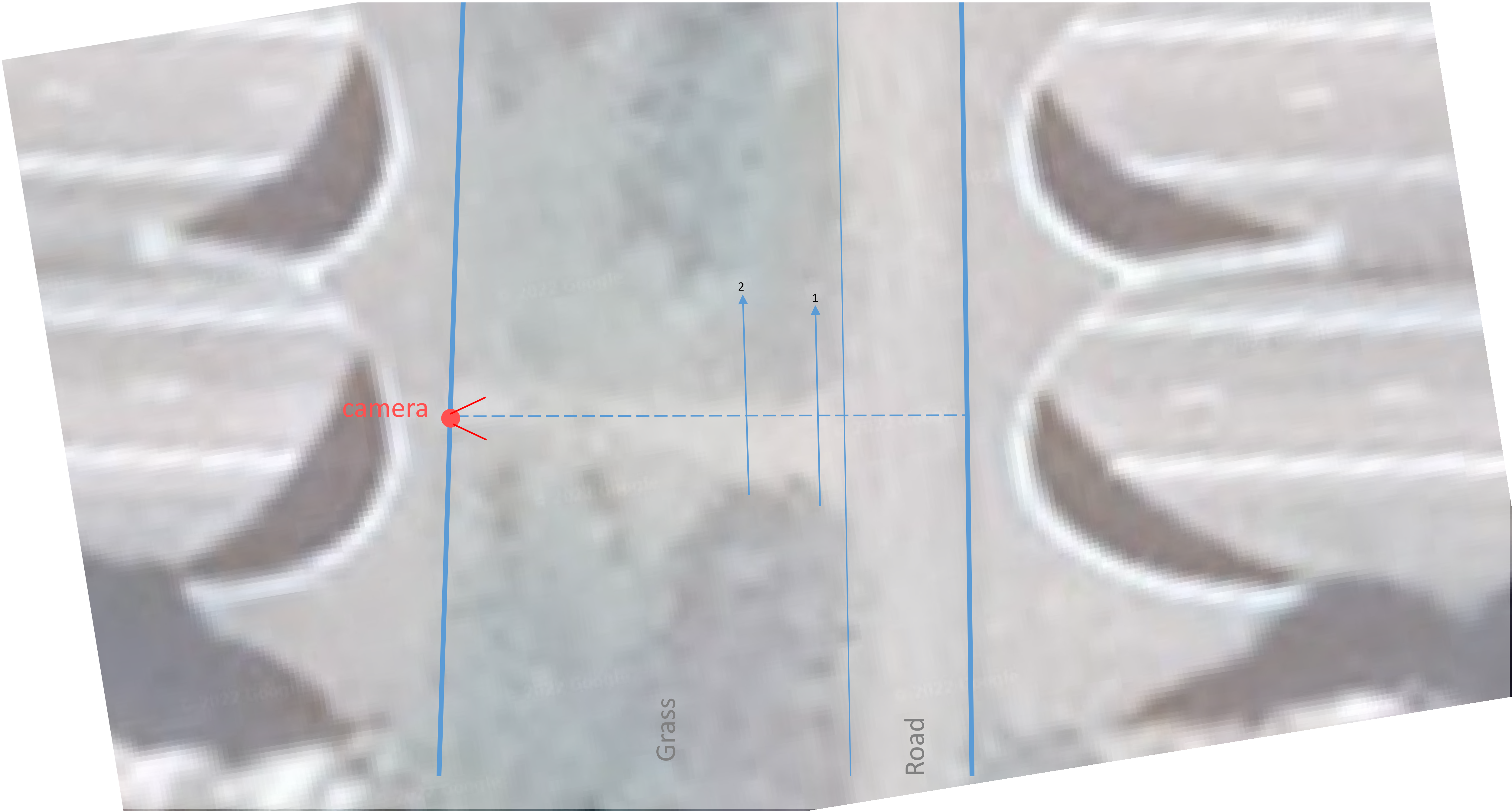

### Kärmekorpi, roe deer, snow free

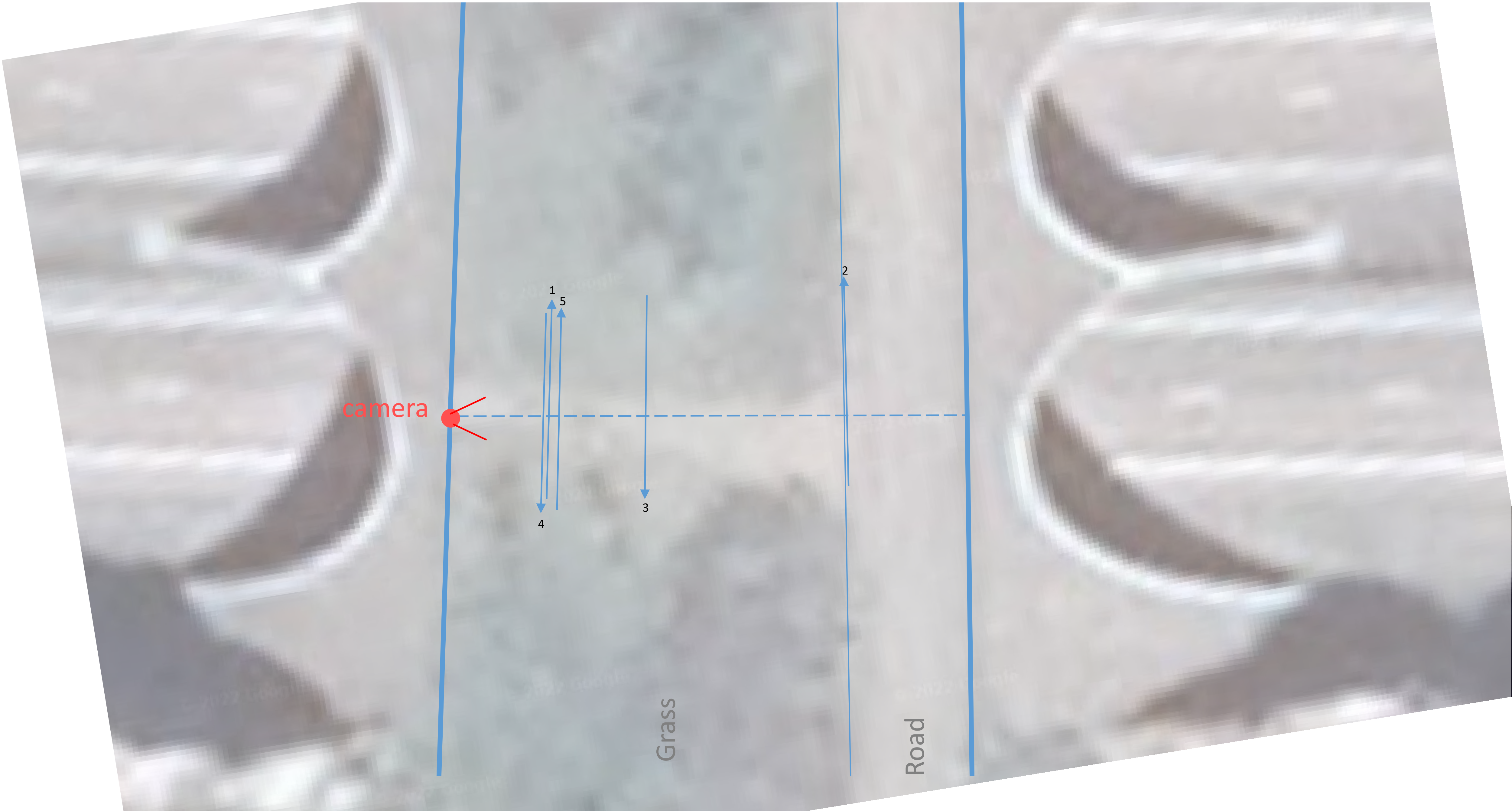

### Kärmekorpi, white-tailed deer, snow free

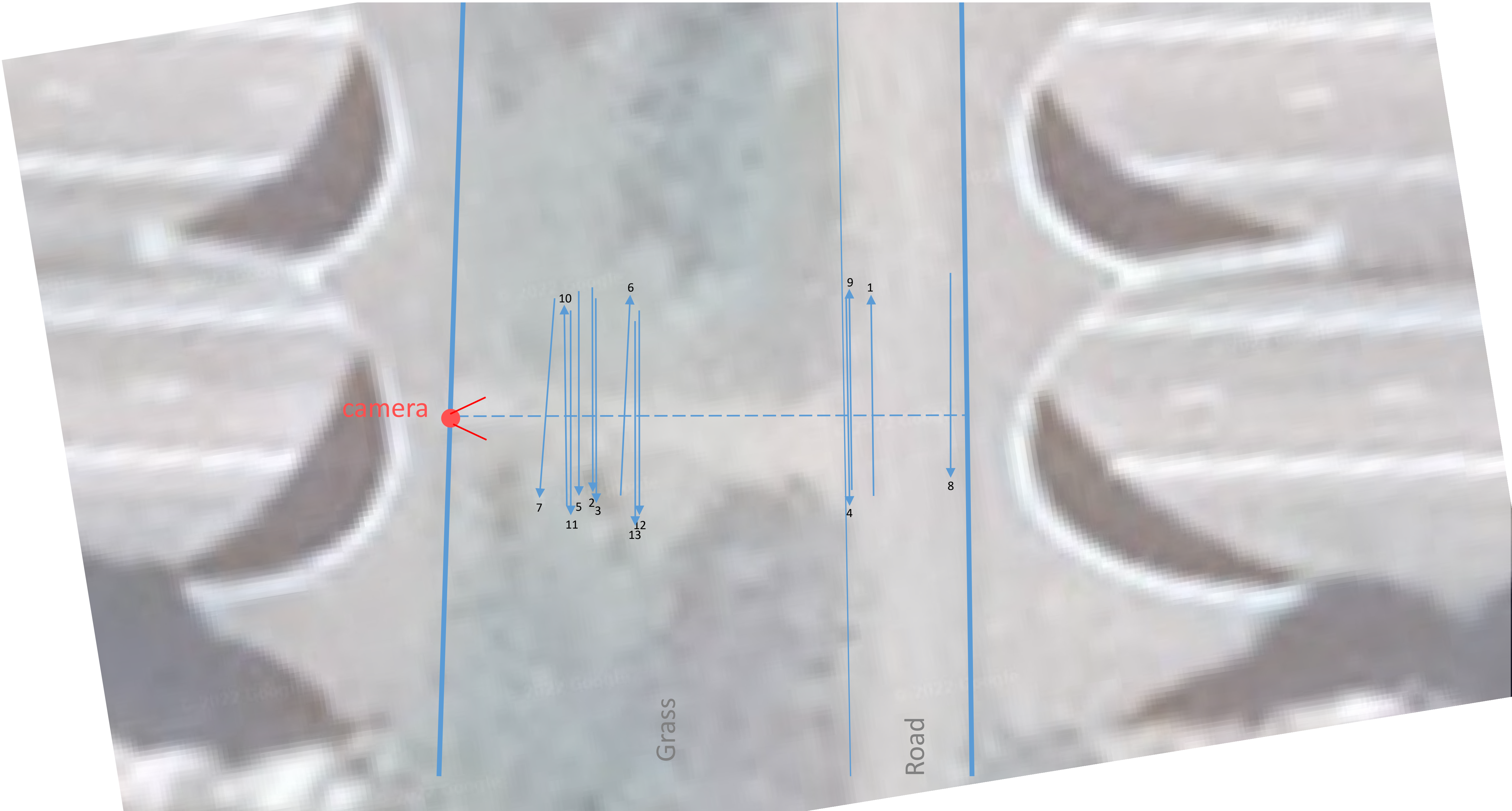

### Kärmekorpi, wildboar, snow free

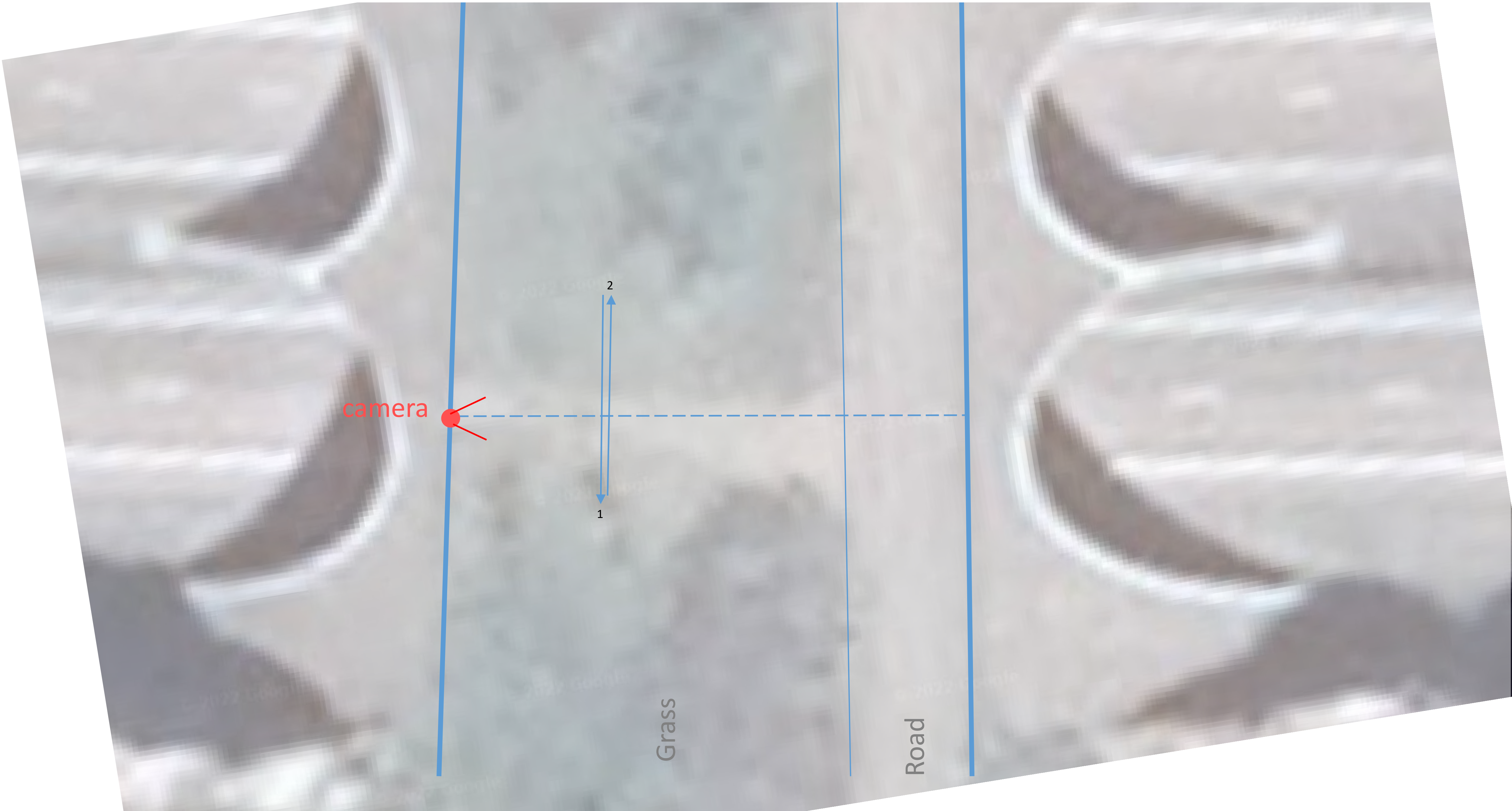
