## Supplementary material for "Ungulate substrate use in fauna passages": Data suppl 3 Fauna passages info

### Data supplement no. 3 to manuscript Ungulate substrate use in fauna passages

#### General information on fauna passages included in the study

*J.O. Helldin & M. Niemi*
*Submitted to European Journal of Wildlife Research*

Notes:

- The dimensions width and length were measured as defined by the wildlife fencing. Since the fences through the passages in most cases follow an hourglass shape, the width refers to the narrowest place, and the length is approximate.
- Frequency of traffic and other human activities were calculated from observations in camera traps during the study.

#### Kvarnbäcken underpass

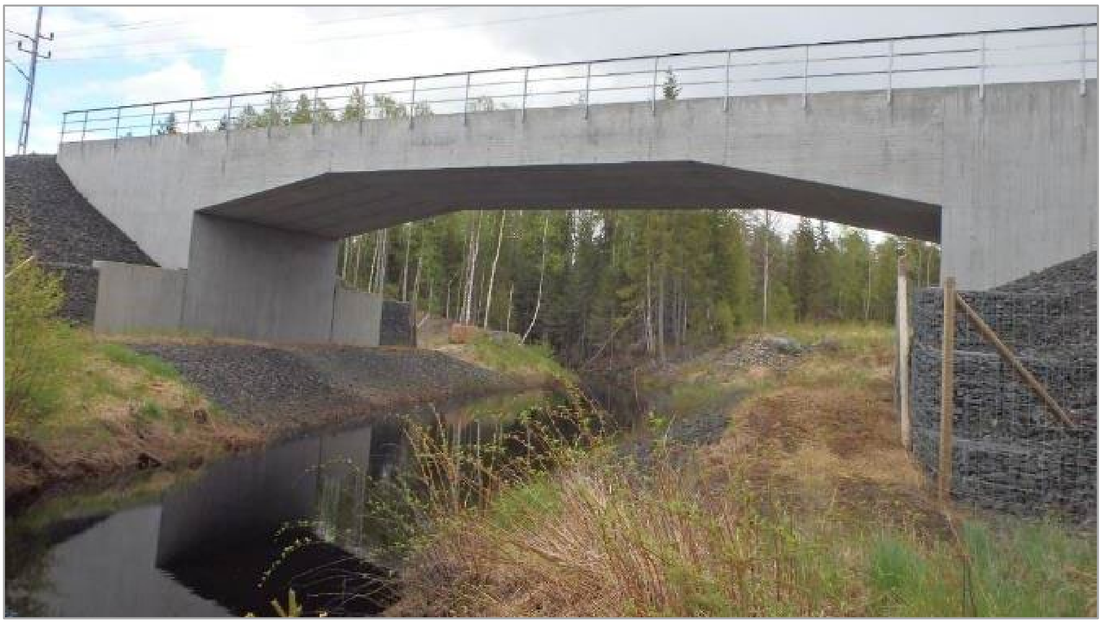

*Photo: Swedish Transport Administration*

| | |
|---|---|
| Location (WGS84) | 65°51'13.7"N 23°24'2.1"E |
| Technical label (Swedish Transport Administration) | Knr 3500-10003-1 |
| Construction year | 2012 |
| Infrastructure crossed | Haparandabanan railway |
| Width (between fences) | 22 m |
| Height (ground to ceiling) | 4-5 m |
| Length | ca 35 m |
| Traffic through passage (off-road vehicles) | <0.1 events per day |
| Other human activities in passage (pedestrians, bicycles etc.) | <0.1 events per day |
| Monitoring start (y-m-d) | 2019-07-14 |
| Monitoring end (y-m-d) | 2020-07-21 |
| Number of days (24h) effectively monitored | 372 |

#### Aavajoki underpass

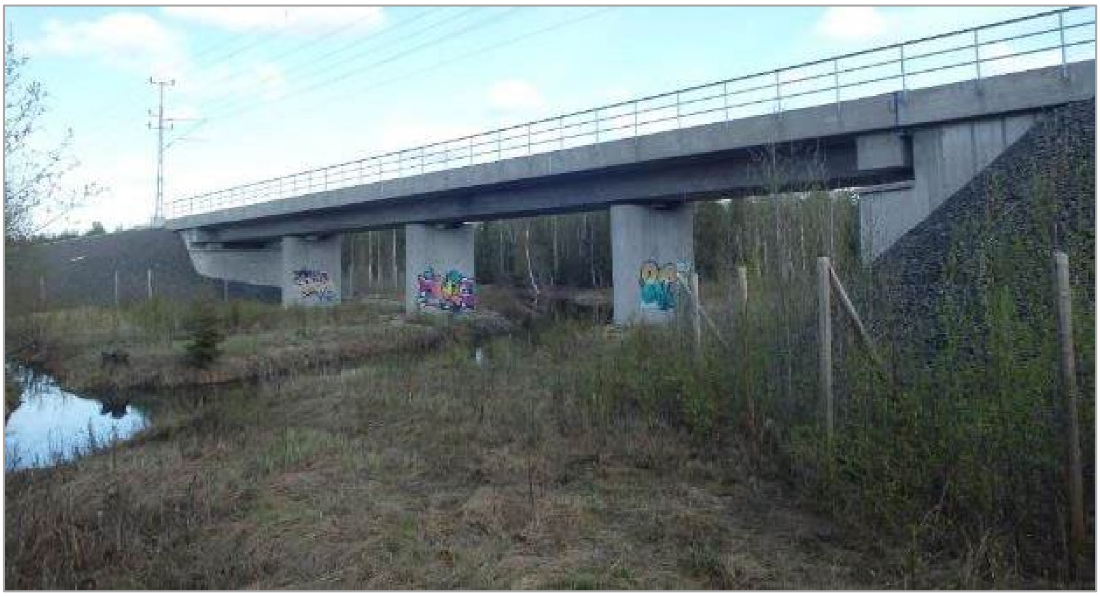

*Photo: Swedish Transport Administration*

| | |
|---|---|
| Location (WGS84) | 65°51'19"N 23°47'23"E |
| Technical label (Swedish Transport Administration) | 3500-5759-1 |
| Construction year | 2012 |
| Infrastructure crossed | Haparandabanan railway |
| Width (between fences) | 40 m |
| Height (ground to ceiling) | 5 m |
| Length | ca 45 m |
| Traffic through passage (off-road vehicles) | 0.1 events per day |
| Other human activities in passage (pedestrians, bicycles etc.) | 0.1 events per day |
| Monitoring start (y-m-d) | 2019-11-16 |
| Monitoring end (y-m-d) | 2020-11-12 |
| Number of days (24h) effectively monitored | 362 |

#### Keräsjoki underpass

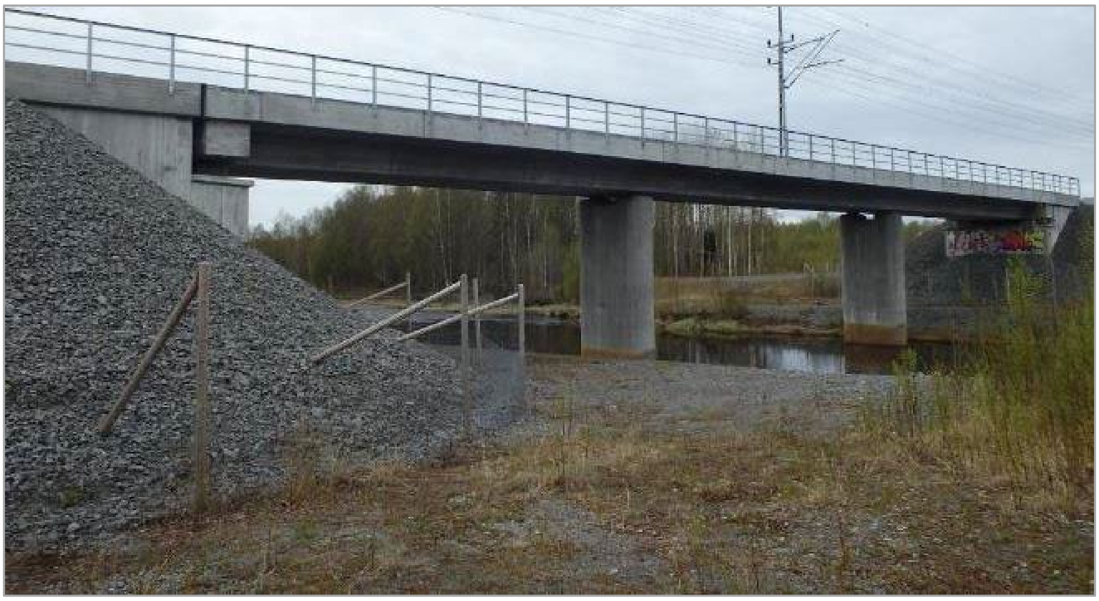

*Photo: Swedish Transport Administration*

| | |
|---|---|
| Location (WGS84) | 65°50'55"N 23°53'45"E |
| Technical label (Swedish Transport Administration) | 3500-5761-1 |
| Construction year | 2012 |
| Infrastructure crossed | Haparandabanan railway |
| Width (between fences) | 44 m |
| Height (ground to ceiling) | 5-6 m |
| Length | ca 35 m |
| Traffic through passage (off-road vehicles) | 0.2 events per day |
| Other human activities in passage (pedestrians, bicycles etc.) | 0.3 events per day |
| Monitoring start (y-m-d) | 2019-07-14 |
| Monitoring end (y-m-d) | 2020-07-20 |
| Number of days (24h) effectively monitored | 371 |

#### Sangijärvi overpass

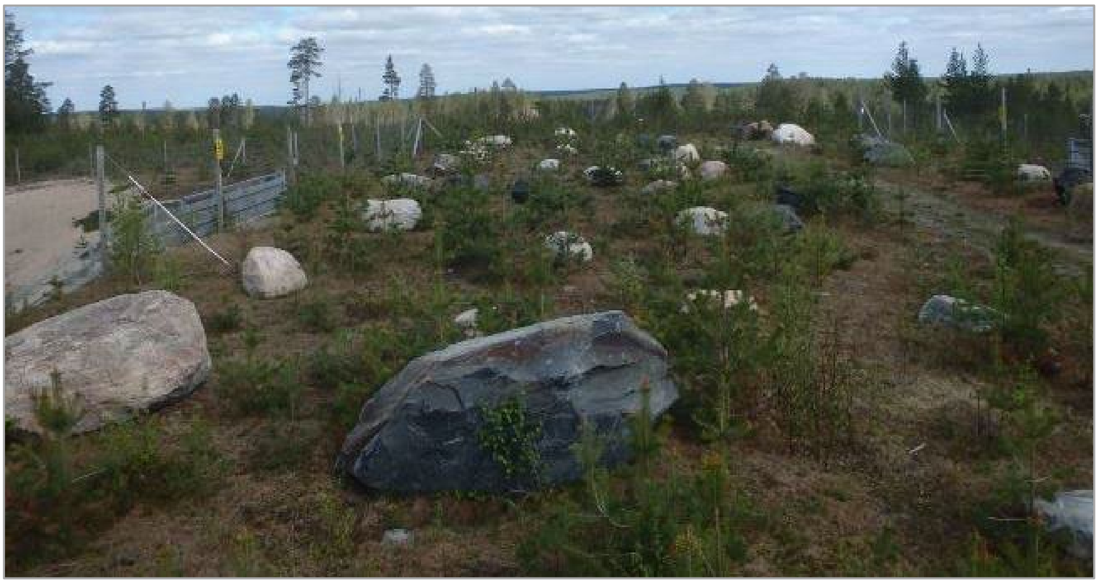

*Photo: Swedish Transport Administration*

| | |
|---|---|
| Location (WGS84) | 65°51'58"N 23°34'12"E |
| Technical label (Swedish Transport Administration) | 3500-10032-1 |
| Construction year | 2012 |
| Infrastructure crossed | Haparandabanan railway |
| Width (between fences) | 20 m |
| Length | ca 50 m |
| Traffic through passage (including off-road vehicles) | 0.4 events per day |
| Other human activities in passage (pedestrians, bicycles etc.) | 0.4 events per day |
| Monitoring start (y-m-d) | 2018-11-18[a] |
| Monitoring end (y-m-d) | 2020-11-12[a] |
| Number of days (24h) effectively monitored | 366 |

a) Only from 2019-11-01 to 2018-10-31 used in the current study

#### Sammatti overpass

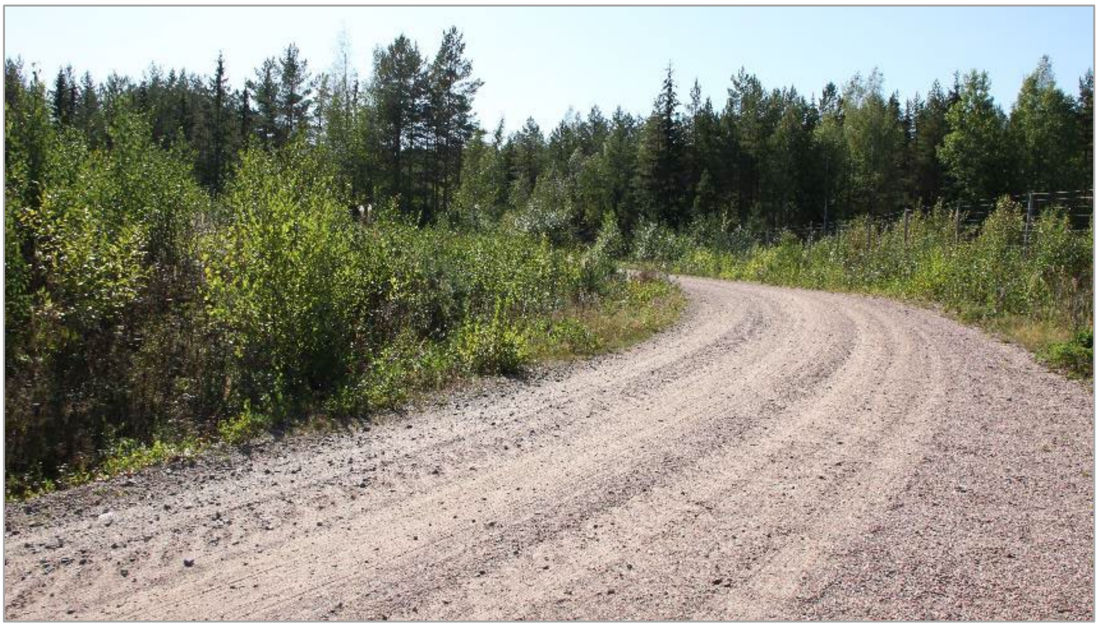

*Photo: Milla Niemi*

| | |
|---|---|
| Location (WGS84) | 60°22'37"N 23°49'7"E |
| Technical label (Finnish Transport Infrastructure Agency) | U-3650 |
| Construction year | 2008 |
| Infrastructure crossed | Road E18 (Finnish road 1) |
| Width (between fences) | 24 m |
| Length | ca 60 m |
| Traffic through passage (including off-road vehicles) | 5.2 events per day |
| Other human activities in passage (pedestrians, bicycles etc.) | 0.8 events per day |
| Monitoring start (y-m-d) | 2019-12-01 |
| Monitoring end (y-m-d) | 2020-11-30 |
| Number of days (24h) effectively monitored | 353 |

#### Loviisa overpass

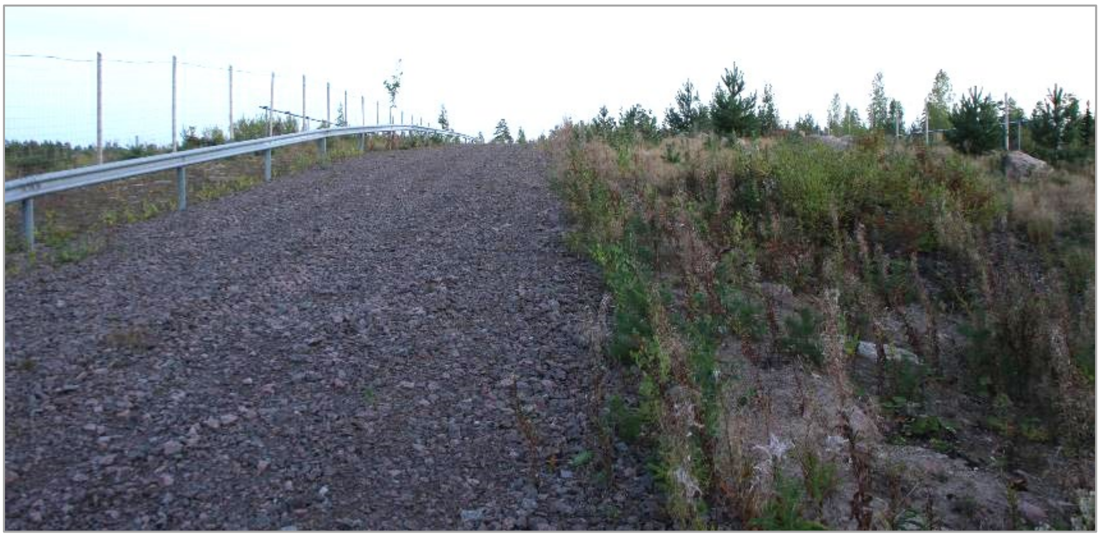

*Photo: Milla Niemi*

| | |
|---|---|
| Location (WGS84) | 60°29'23"N 26°19'38"E |
| Technical label (Finnish Transport Infrastructure Agency) | U-3740 |
| Construction year | 2014 |
| Infrastructure crossed | Road E18 (Finnish road 7) |
| Width (between fences) | 27 m |
| Length | ca 80 m |
| Traffic through passage (including off-road vehicles) | 0.3 events per day |
| Other human activities in passage (pedestrians, bicycles etc.) | 0.3 events per day |
| Monitoring start (y-m-d) | 2019-12-01 |
| Monitoring end (y-m-d) | 2020-11-30 |
| Number of days (24h) effectively monitored | 330 |

#### Kärmekorpi overpass

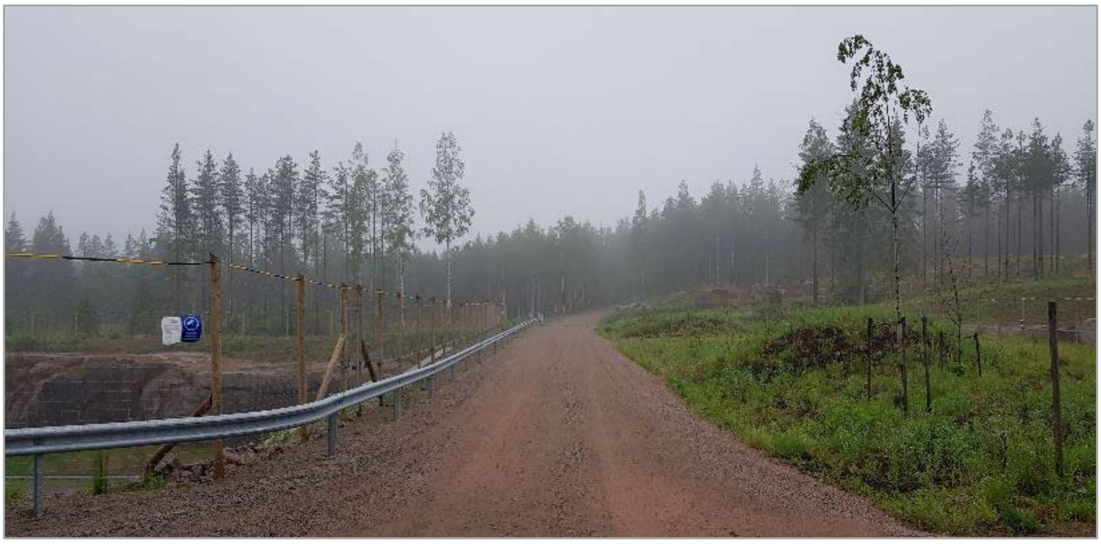

*Photo: Milla Niemi*

| | |
|---|---|
| Location (WGS84) | 60°35'33"N 27°30'48"E |
| Technical label (Finnish Transport Infrastructure Agency) | KaS-1452 |
| Construction year | 2018 |
| Infrastructure crossed | Road E18 (Finnish road 7) |
| Width (between fences) | 29 m |
| Length | ca 60 m |
| Traffic through passage (including off-road vehicles) | 2.7 events per day |
| Other human activities in passage (pedestrians, bicycles etc.) | 0.3 events per day |
| Monitoring start (y-m-d) | 2019-12-01 |
| Monitoring end (y-m-d) | 2020-11-30 |
| Number of days (24h) effectively monitored | 358 |
